# Exploring Degradation of Intrinsically Disordered Protein YAP induced by PROTACs

**DOI:** 10.1101/2023.09.19.556013

**Authors:** Chen Zhou, Chunbao Sun, Liya Pi, Chenglong Li

## Abstract

Yes-associated protein (YAP), a potent oncogene and a key player in the Hippo tumor suppression pathway, has long been considered challenging to target due to its partially intrinsically disordered nature. However, recent advances in High-throughput Screening (HTS) have led to the discovery of a few YAP binders. Building upon this progress, a novel approach utilizing Proteolysis-Targeting Chimera (PROTAC) technology was employed to design and synthesize a series of YAP degraders. Here, our degraders were created by linking NSC682769, a previously reported YAP binder, with either VHL ligand 2 or pomalidomide using various linkers of different lengths and types. The most promising degrader **YZ-6** recruits the E3 ligase VHL, inducing rapid and sustained YAP degradation leading to suppression of YAP/TEAD-led transcription in both YAP-dependent NCI-H226 and Huh7 cancer cell lines. In addition to its degradation capabilities, **YZ-6** also exhibited potent antiproliferative activity in both cell lines. Importantly, **YZ-6** efficiently suppresses tumor development in the Huh7 xenograft mouse model without adverse effects on the mice. These findings highlight the potential of PROTAC-mediated degradation as a viable strategy for reducing oncogenic YAP levels and attenuating downstream signaling in cancer cells. Moreover, the development of PROTACs based on NSC672869 holds promise for treating YAP-driven malignancies.

## Introduction

Yes-associated protein (YAP) and transcriptional co-activator with PDZ-binding motif (TAZ), prominent downstream regulators of the Hippo pathway, play crucial roles in regulating tissue homeostasis, organ size, regeneration and tumorigenesis.^1–3^ The Hippo signaling pathway regulates the co-activator function of YAP/TAZ by controlling their cellular localization through phosphorylation.^4^ Activation of the upstream Hippo signaling pathway leads to the phosphorylation of large tumor suppressor 1/2 (Lats1/2) by mammalian STE20-like protein kinase 1/2 (Mst1/2) or activation by neurofibromin 2 (NF2), which acts as a scaffolding protein in the pathway.^5^ Subsequently, phosphorylated Lats1/2 phosphorylates YAP/TAZ in the cytoplasm. Importantly, phosphorylated YAP/TAZ are recognized by protein 14-3-3, which sequesters these co-activators in the cytoplasm, resulting in the suppression of Hippo-induced gene transcription.^3, 6^ Conversely, when YAP/TAZ are not phosphorylated, the co-activators are able to translocate into the nucleus by an unknown mechanism, bind the TEAD transcription factors and drive transcriptional targets, including genes involved in cell growth and proliferation, most notably connective tissue growth factor (CTGF) and Cysteine-rich angiogenic inducer 61 (Cyr61). ^7–9^

More importantly, YAP and TAZ are considered oncogenes, as their amplification or over-expression has been observed in various human cancers, and reports of gene amplification and epigenetic modulation of the YAP and TAZ loci in cancer are similarly prevalent.^9–11^ Studies have demonstrated that elevated YAP expression can trigger multiple oncogenic characteristics in mammalian cells.^4^ Additionally, the nuclear localization of YAP in tumor biopsies has been correlated with a poor prognosis for cancer patients.^6, 12, 13^

YAP and its paralogue TAZ are characterized as intrinsically disordered proteins.^14, 15^ Due to their inherent flexibility, they have the ability to adopt diverse conformations, making them challenging targets for intervention.^16^ However, recent advances have demonstrated successful targeting of intrinsically disordered proteins using small molecules.^17^ In fact, clinical trials are currently underway for potential therapeutics that targets the disordered region of androgen receptor.^17^ Although targeting YAP/TAZ presents significant challenges, progress in targeting other intrinsically disordered proteins provides hope for potential strategies aimed at modulating YAP/TAZ activity.

High-throughput screening (HTS) is indeed the primary method used for targeting disordered proteins.^18^ HTS involves experimental screening of large libraries of small molecules to identify compounds that can interact with the target protein. In the case of YAP/TAZ, HTS has been successfully employed to identify several small molecules that have the potential to bind to and modulate the activity of YAP/TAZ.^19^ Verteporfin (VP) (**Figure 1**), a drug commonly used in the treatment of macular degeneration, is the first and most renowned YAP inhibitor obtained through a luciferase reporter assay screening.^20^ It has been demonstrated to bind to YAP, leading to a conformational change in the protein and disrupting its interaction with TEAD. However, further investigations have revealed the presence of off-target effects that are independent of YAP.^21, 22^ These studies have identified additional effects of verteporfin that are not directly related to its inhibition of YAP, highlighting the complexity and potential limitations of using verteporfin as a specific YAP inhibitor.^23^ In the study conducted by Oku et al., a screening was performed to identify small molecules capable of inhibiting the nuclear localization of YAP/TAZ.^24^ The researchers discovered that three drugs, namely dasatinib, fluvastatin, and pazopanib, were effective in inhibiting the nuclear localization and targeting gene expression of YAP and TAZ (**Figure 1**). All three drugs induced the phosphorylation of YAP and TAZ, which is an important regulatory modification. Additionally, pazopanib specifically triggered the proteasomal degradation of YAP/TAZ, contributing to their transcriptional suppression. Interestingly, Taccioli et al., using the Mutations and Drugs Portal (MDP) at http://mdp.unimore.it, also found that individually, dasatinib and fluvastatin were able to partially reduce the nuclear localization of YAP/TAZ. However, when these two drugs were combined, they observed a complete exclusion of YAP/TAZ from the nucleus. This combination therapy demonstrated a synergistic effect in inhibiting the nuclear translocation of YAP/TAZ, which is important for their oncogenic activity.^25^ Utilizing a high-throughput yeast two-hybrid based screen, NSC682769 (**Figure 1**) was identified to inhibit the association of the co-transcriptional activator YAP1 and the TEA domain family member 1 (TEAD1) transcription factor protein–protein interaction.^26^ NSC682769 potently blocked association of YAP and TEAD *in vitro* in GBM cells treated with submicromolar concentrations of drug. To investigate the mechanism of action, the researchers performed surface plasmon resonance (SPR) analysis to study the binding of NSC682769 to immobilized YAP. They observed that NSC682769 bound to YAP in a concentration-dependent manner and quickly reached binding equilibrium. The dissociation constant (K_D_) was determined to be 738 nM, indicating a direct interaction between NSC682769 and YAP. Further confirmation was obtained through inhibitor-coupled bead pull-down assays, which further demonstrated that NSC682769 binds directly to YAP. Collectively, these findings provide evidence that NSC682769 directly binds to YAP, preventing its association with TEAD and inhibiting their protein-protein interaction. This study highlights NSC682769 as a potential therapeutic compound for disrupting YAP/TEAD-induced oncogenic signaling.

**Figure 1.**
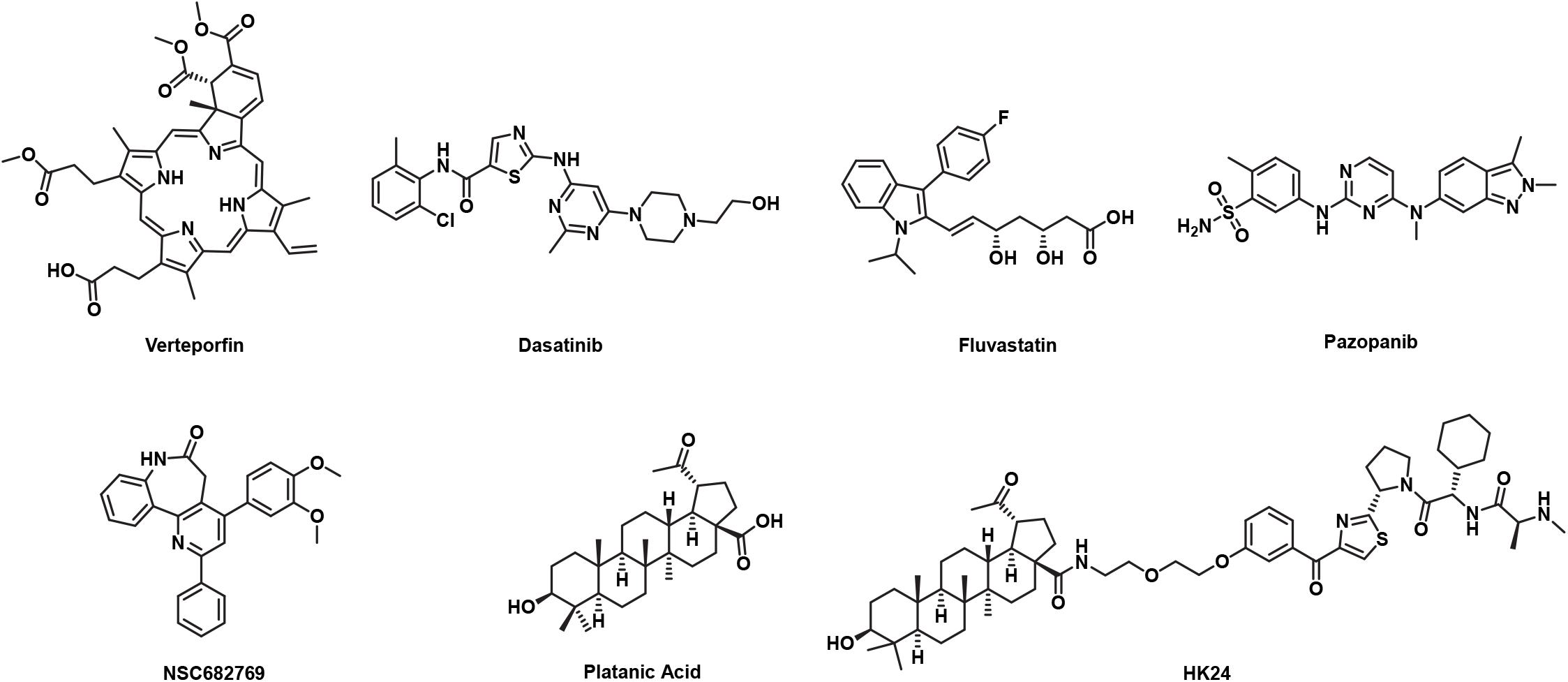
Representative compounds targeting YAP.

In the meanwhile, computational methods can also be utilized to identify binders for disordered proteins.^27, 28^ However, the process of identifying molecules that interact with disordered regions requires additional steps similar to traditional drug design approaches that focus on well-ordered pockets. To reliably identify binders for disordered proteins, the first step involves characterizing the conformational ensembles of the target protein. This can be achieved through a combination of computational techniques and experimental methods. Once the conformational ensembles are established, the next step is to select druggable conformers. Ligands that bind to druggable conformers can then be identified using virtual screenings based on ensemble docking or receptor-based pharmacophore models. However, it is worth noting that the shift from the traditional drug design pipeline that primarily targets well-ordered pockets to the identification of binders for disordered proteins has been relatively slow and has not gained significant momentum.

More recently, an increasing interest in PROteolysis TArgeting Chimeras (PROTACs) has become apparent, spurred by research interests into ‘undruggable’ targets.^29, 30^ These bifunctional molecules simultaneously interact with a protein of interest (POI) and an E3 ligase, resulting in the formation of a ternary complex.^31, 32^ This complex facilitates the E3 ligase’s ability to attach ubiquitin molecules to proximal lysine residues on the POI. Subsequently the ubiquitinated POI is recognized and degraded by the 26S proteasome.^33^ One of the significant advantages of this targeted degradation is its capacity to eliminate proteins of interest that are not typically attenuated by traditional small-molecule inhibitors.^34–36^ Expanding the range of druggable protein targets is possible through the design of potent and effective PROTAC degraders, as the ligand is not required to bind to a specific functional site in the POI.^37^ This unique characteristic opens up opportunities to target a wider variety of proteins for degradation including intrinsically disordered protein.

At this time, there is only one reported YAP PROTAC. Upon screening YAP-interacting small molecule compounds, Nakano et al. identified HK13, a platanic acid (**Figure 1**), as a novel compound that interacts with YAP which is validated by SPR.^38^ However, it was observed that HK13 exhibited non-specific binding to YAP at high concentrations, raising concerns about its specificity. Based on the YAP-binding capability of HK13, the researchers synthesized a series of YAP degraders using PROTAC technology. These degraders linked platanic acid with LCL-161, a compound known for its high affinity for cIAP (cellular inhibitor of apoptosis protein), an E3 ubiquitin ligase. Among these compounds, HK24 (**Figure 1**) showed the ability to inhibit the growth of NCI-H290 cells that overexpress YAP and to induce YAP degradation in a dose-dependent manner. However, it should be noted that HK24 exhibited weak YAP degradation, less than 50% degradation at 5 μM. To investigate the mechanism of YAP degradation induced by HK24, the researchers cultured NCI-H290 cells with HK24 in the presence of a proteasome inhibitor, confirming the involvement of the proteasome pathway in HK24-dependent YAP degradation. However, no *in vivo* studies were conducted as part of this research.

NSC682769 is one of the few small molecules whose reported binding to YAP validated by SPR with a K_D_ value less than 1 μM. Notably, NSC682769 has a lower molecular weight compared to VP. Furthermore, in preclinical models including tumor xenografts and genetically engineered mouse models, NSC682769 has demonstrated significant anti-tumor responses, increased overall survival, and notable penetration across the blood-brain barrier (BBB).^26^ Based on these promising characteristics, we designed and synthesized a series of novel YAP PROTACs based on NSC682769. These PROTACs were created by linking NSC682769 with either pomalidomide or a VHL ligand, using various linker lengths and types. The resulting compounds displayed rapid degradation of YAP in cells overexpressing YAP, functioning through a *bona fide* PROTAC mechanism. The observed acute and sustained degradation of YAP in multiple cancer cell lines by these PROTACs, specifically **YZ-6**, demonstrates their potential as valuable tools for studying YAP biology. Moreover, this research represents a significant advancement in the development of PROTAC-based therapeutics targeting the degradation of oncogenic YAP. Importantly, by successfully inducing the degradation of YAP, which is an intrinsically disordered protein often considered undruggable, this study provides a compelling proof of concept for the application of PROTACs in degrading such challenging protein targets.

## Results and Discussion

### Design and Biological Evaluation of NSC682769-Based YAP PROTAC Degraders

In view of its excellent *in vitro* and *in vivo* activities, as well as its synthetic tractability, we selected NSC682769 as a starting point to design YAP-targeting PROTACs.^26^ However, due to the lack of an experimental structure of NSC682769 in complex with YAP, the binding mode and solvent-exposed regions of NSC682769 are not known, making the design of YAP PROTACs more challenging. To overcome this, we attempted to identify suitable tethering sites for linker attachment based on the structure of NSC682769. We explored four potential tethering sites: site 1, the NH of the amide; site 2, the para substitution of the methoxy group; site 3, the para substitution of the 2-phenyl ring; and site 4, the meta substitution of the methoxy group (as shown in **Figure 2**). Using short linker moieties at these tethering sites, we synthesized compounds **7**, **12**, **13**, **14**, **19**, **20**, **21**, **25**, **26**, and **27**. We then employed a TEAD-dependent luciferase reporter assay to determine if these compounds retained YAP/TEAD transcriptional inhibition activity. The retention of activity would indicate that the respective tethering sites are available for linker attachment without affecting the binding to YAP. Excitingly, compounds **12**, **13** and **15**, which use tethering site 2, as well as compounds **25**, **26**, and **27**, which use tethering site 4, retained TEAD transcription inhibition activities (**Figure 2B**). This suggests that tethering site 2 and 4 may be solvent-exposed and good egress positions from the YAP protein for a linker design. With the ligand NSC682769 in hand and the linker tethering sites determined, we carried out a PROTAC design and synthesis campaign (**Figure 3**). We employed flexible alkyl or ether moieties to connect the NSC682769 ligand from tethering sites 2 and 4. Not knowing a priori at what distance these ligands would have to be positioned in the PROTAC to effectively associate their respective proteins, we undertook a systematic survey of connector length. Since the design of PROTAC YAP degraders also requires a small-molecule ligand for an E3-ligase, the von Hippel–Lindau protein 1 (VHL-1)/cullin 2 E3 ligase system has been employed for the design of PROTAC degraders of a number of proteins.^39–42^ We thus investigated whether the VHL-1/cullin 2 E3 ligase system can be employed here for the successful design of PROTAC YAP degraders. We used VHL ligand 2 which has been successfully used for the design of BCL-X_L_ PROTAC degrader DT2216, which is the only VHL-recruiting PROTAC in clinical trial.^43, 44^ In addition to the VHL-1/cullin 2 E3 ligase system, CRBN/cullin 4A E3 ligase has been employed extensively for the design of PROTAC degraders of various proteins.^45–47^ Pomalidomide is a potent, small-molecule ligand which binds to cereblon (CRBN), an adaptor protein in the cullin 4A E3 ligase degradation system. Pomalidomide has been successfully used in the design of PROTAC degraders for a number of proteins and it has more oral drug-like property compared with VHL ligand due to its smaller molecular weight.^48–50^ Therefore, we employed both VHL ligand 2 and pomalidomide as E3 ligase ligands in this study.

**Figure 2.**
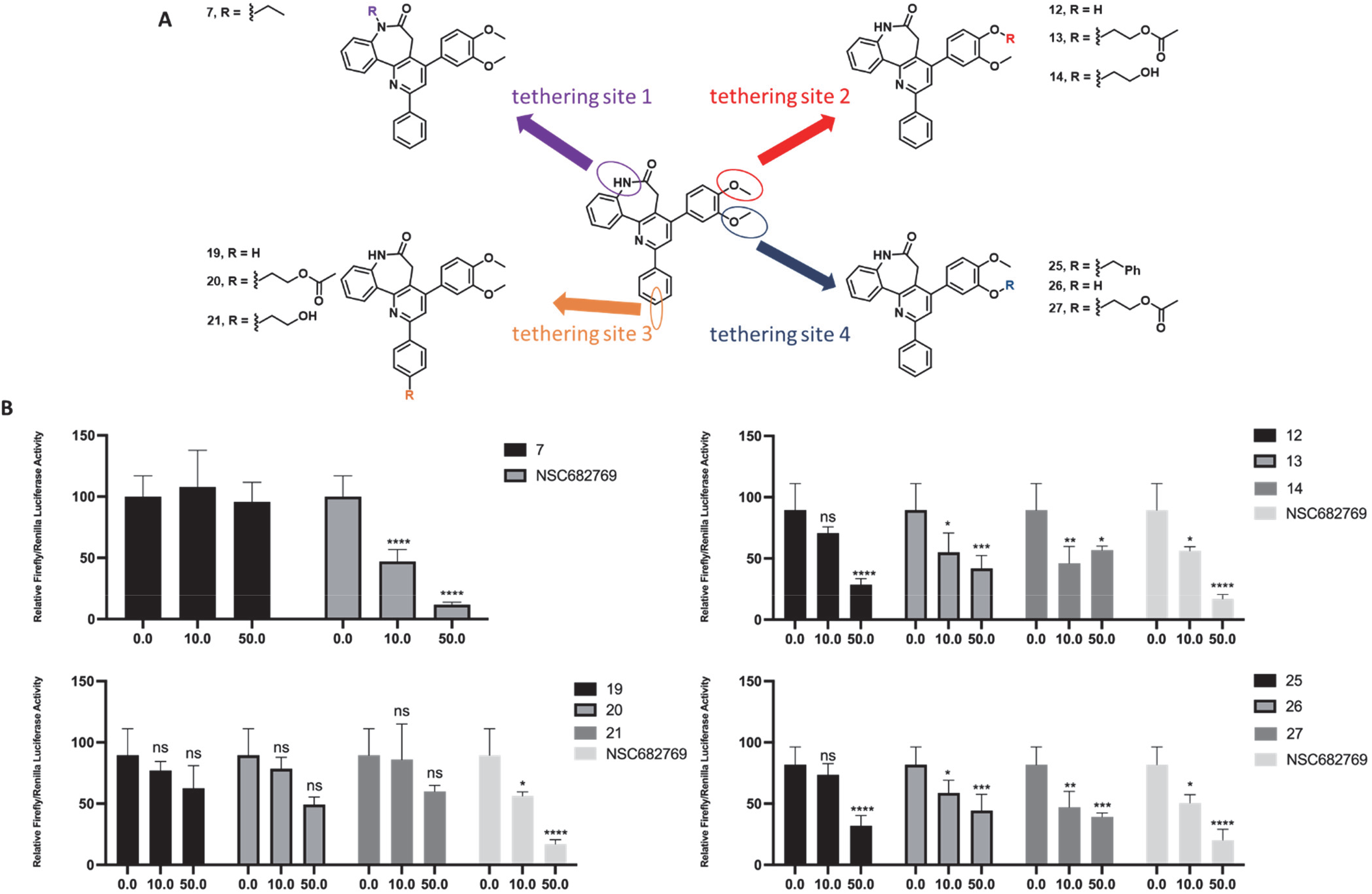
The strategy to find the suitable tethering site for the design of YAP PROTACs. (A) Structures of designed compounds built from four tethering sites with small linker attached. (B) TEAD dependent luciferase reporter assay. HEK 293T cells were treated with DMSO, and designed compounds or NSC68269 for 24h. Not Significant (N.S.); * p < 0.05; ** p < 0.01; *** p < 0.005; **** p < 0.001.

**Figure 3.**
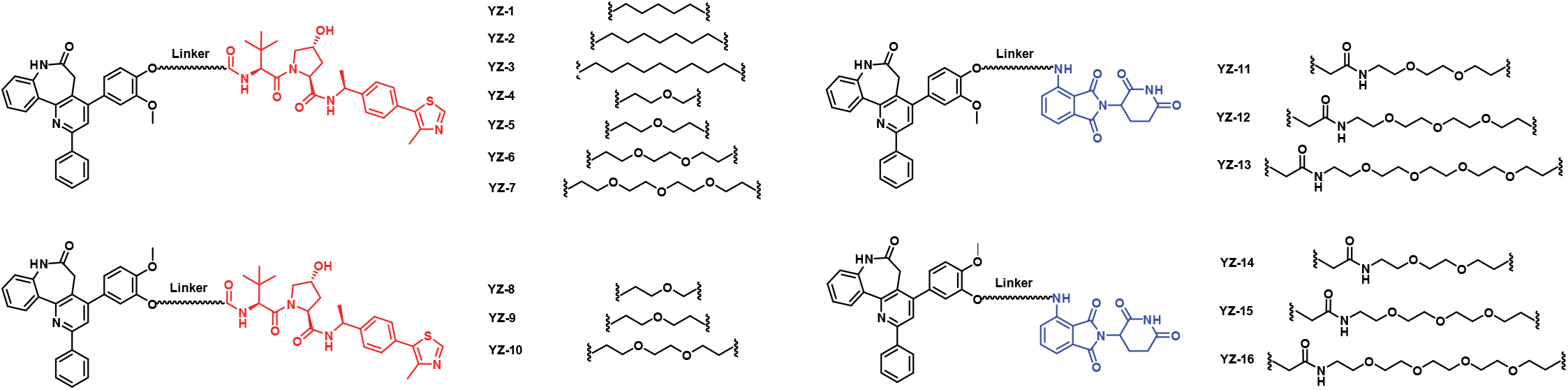
Chemical structure of designed NSC682769-based YAP degraders using tethering site 2 and 4.

In general, the NCI-H226 cell line is recognized as a well-characterized YAP-dependent mesothelioma cell line,^51, 52^ and among the three YAP-TEAD inhibitors currently undergoing clinical trials, all of them are intended for the treatment of mesothelioma (Clinical trial information: NCT05228015, NCT04857372, NCT04665206). In order to gain initial insights into the cellular effects of our designed YAP degraders, we evaluated the potency of each synthesized compound in terms of cell growth inhibition using this NCI-H226 cell line here. The resulting data are summarized in **Figure 4**.

**Figure 4.**
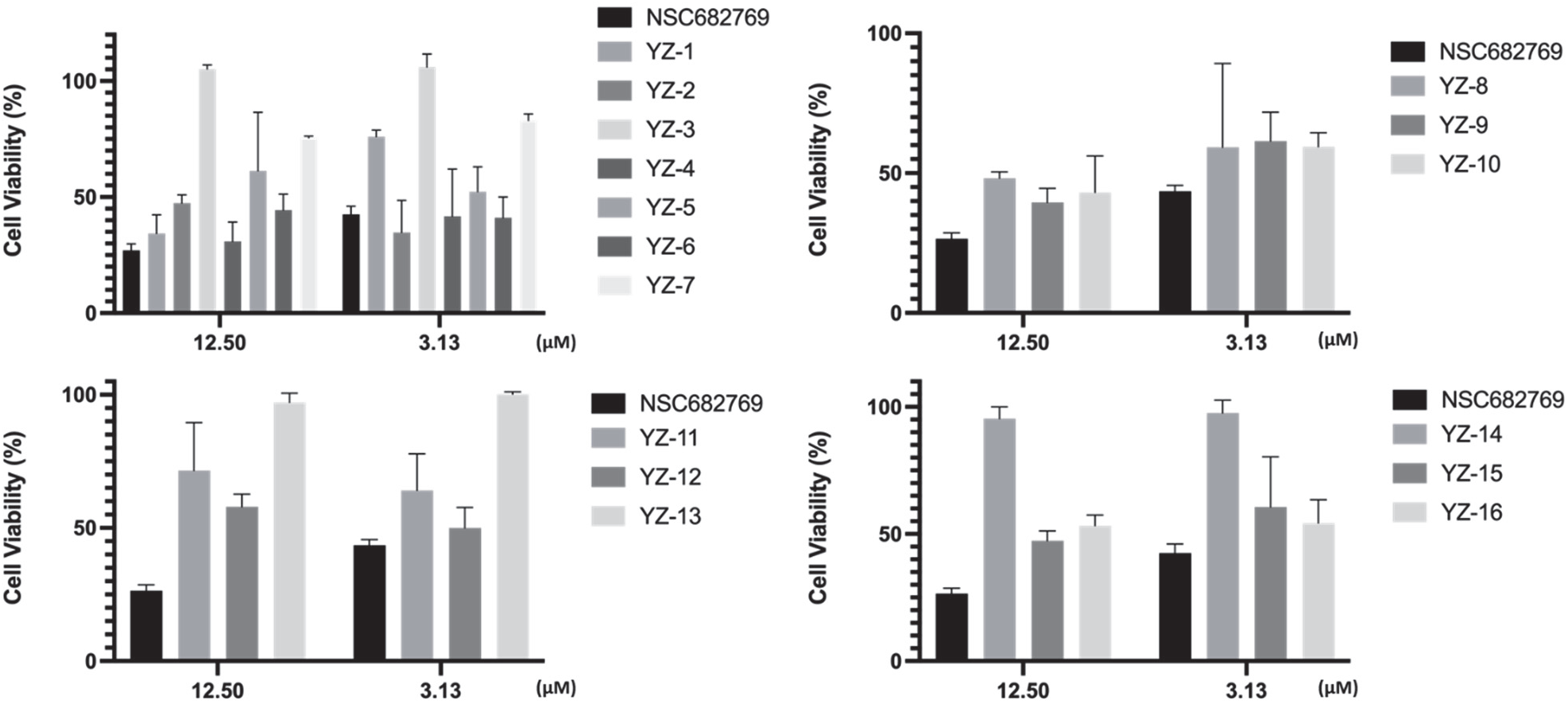
Cell viability assay. NCI-H226 cells were treated with indicated doses of compounds for 3 days. Cell viability was determined by CCK8 assay.

**YZ-3** and **YZ-13** demonstrated no cell killing activities on NCI-H226 cells, even at a concentration of 12.5 μM. **YZ-3** was synthesized by connecting NSC672869 to VHL ligand 2, while **YZ-13** utilized pomalidomide to recruit CRBN. Consequently, we did not assess the YAP expression level when treated with these two compounds. However, for the remaining 14 compounds that did display cell killing activity, we conducted a western blot assay to evaluate the YAP expression level at 10 μM using the NCI-H226 cell line (**Figure 5**). In the case of compounds **YZ-1** and **YZ-2**, our western blotting analysis showed that both compounds effectively induced YAP degradation at 10 μM in the NCI-H226 cell line. **YZ-2** was more potent than **YZ-1**, reducing the YAP level by more than 50%, indicating that the seven-carbon linker is more favorable than five-carbon linker. For compounds **YZ-4, 5, 6, 7**, which incorporated PEG linkers, western blotting analysis revealed that **YZ-4** and **YZ-5** exhibited similar YAP degradation activities to **YZ-1** and **YZ-2**. Excitingly, **YZ-6**, featuring two-PEG linker, effectively reduced the YAP level by over 80% at 10 μM in NCI-H226 cell line. However, when we further increased the linker length by one PEG group to produce **YZ-7**, the YAP degradation potency was sharply reduced. Therefore, these results validate the important role of the linker in the design of PROTACs. For compounds **YZ-11** and **YZ-12**, which use the CRBN ligand pomalidomide, both compounds moderately induced YAP degradation at 10 μM in NCI-H226 cell line. Unfortunately, when we used tethering site 4 to generate compounds **YZ-8**, **9**, **10** and **YZ-14**, **15**, **16**, none of the compounds were able to induce the degradation of YAP at 10 μM using NCI-H226 cell line. However, they did exhibit NCI-H226 cell growth inhibition activities, suggesting that these compounds function as YAP inhibitors rather than YAP degraders. These findings indicate that only tethering position 2 in NSC682769 can be successfully utilized for the design of potent and effective YAP degraders.

**Figure 5.**
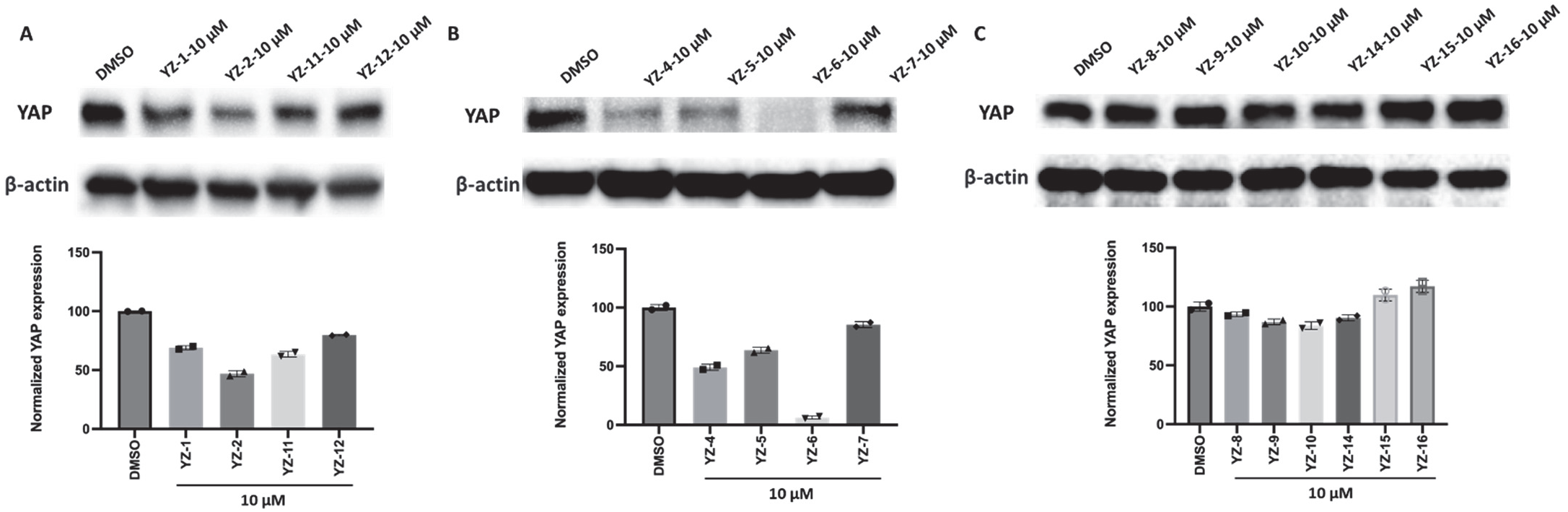
Degradation of YAP protein by synthesized compounds. (A) Western blotting analysis of YAP protein in NCI-H226 cells treated with compounds YZ-1, 2, YZ-11, 12. Quantitation is below. (B) Western blotting analysis of YAP protein in NCI-H226 cells treated with compounds YZ-4, 5, 6, 7. Quantitation is below. (C) Western blotting analysis of YAP protein in NCI-H226 cells treated with compounds YZ-8, 9, 10, YZ-14, 15, 16. Quantitation is below. Cells were treated with different compounds for 48 h, and whole cell lysates were then analyzed by Western blotting to examine the level of YAP protein. The membranes were stripped and reblotted for β-actin as the loading control. Quantified data represents mean ± SD from two independent biological replicates.

### Further evaluation of Two Potent YAP PROTAC Degraders, YZ-4, AND YZ-6

We selected the two most potent compound **YZ-4** and **YZ-6** to further investigate their degradation behaviors. The concentration-dependent YAP degradation by these two compounds is depicted in **Figure 6**. **YZ-4** exhibited a DC_50_ (50% degradation) value of 14.9 μM.^34^ On the other hand, **YZ-6** showed greater potency, with a DC_50_ value of 8.2 μM and a maximum degradation (Dmax) value of 97%.^34^ Additionally, both compounds effectively inhibited YAP-TEAD target gene CTGF expression in a dose dependent manner. Notably, **YZ-6** can firmly inhibit the CTGF expression at a low concentration of 5 μM in NCI-H226 cells, which is more potent than **NSC672869**. Subsequently, we conducted a cell proliferation assay specifically for **YZ-6** (**Figure 7**). Unfortunately, **YZ-6** exhibited an IC_50_ value of 15.3 μM, which is 3-fold less potent than the warhead **NSC682769**. The maximum cell inhibition achieved by **YZ-6** was 60%, whereas **NSC682769** was able to reach 95% cell growth inhibition. Due to solubility limitations, we were unable to increase the concentration further for both compounds.

**Figure 6.**
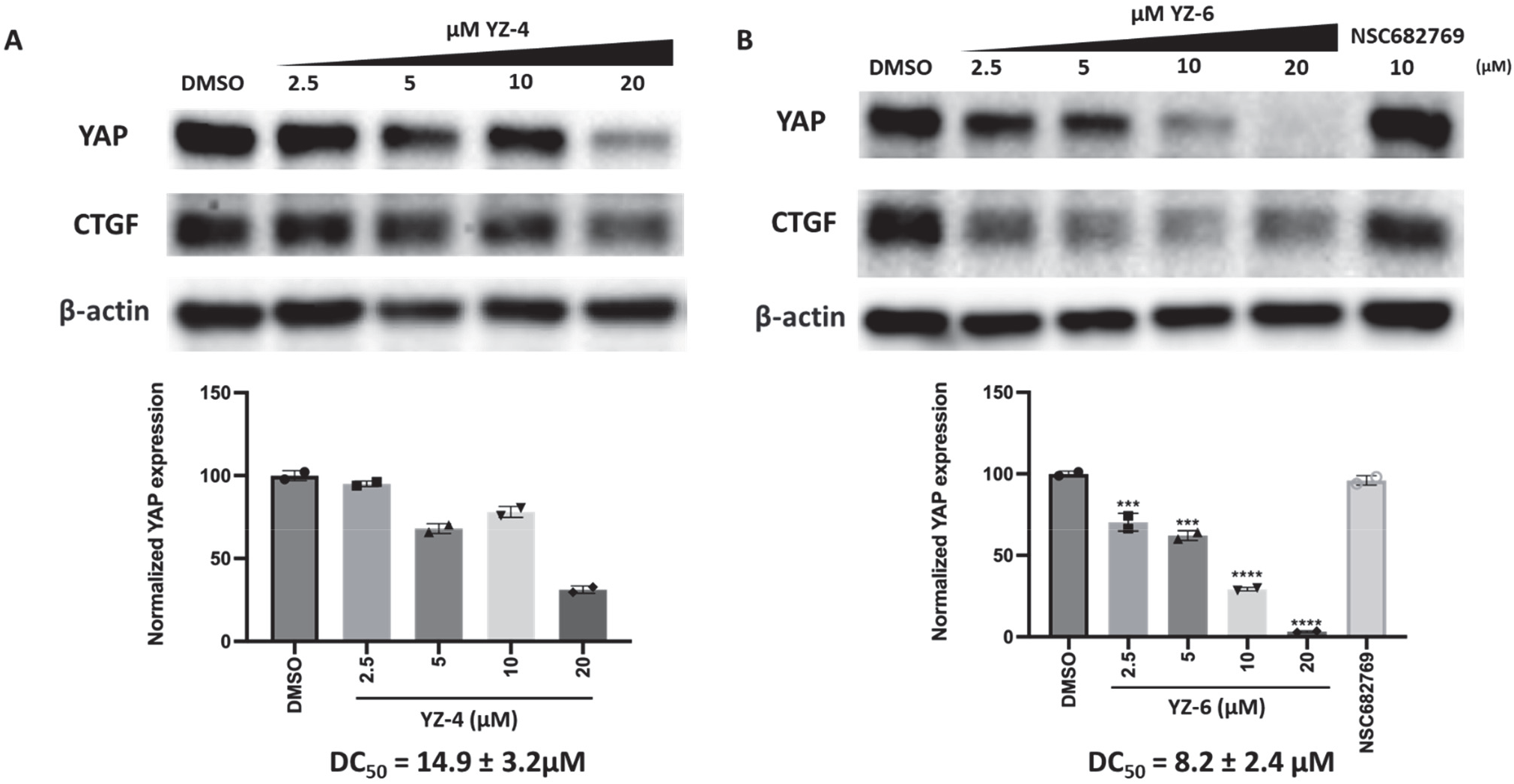
Compounds YZ-4 (A) and YZ-6 (B) concentration-dependently reduced YAP protein level and inhibit YAP-TEAD target gene CTGF expression. Quantitation is below. NCI-H226 cells were treated with DMSO, NSC682769 or serial dilutions of compounds for 48 h. Quantified data represents mean ± SD from two independent biological replicates. **\*\*\* p < 0.005; \*\*\*\*** p < 0.001.

**Figure 7.**
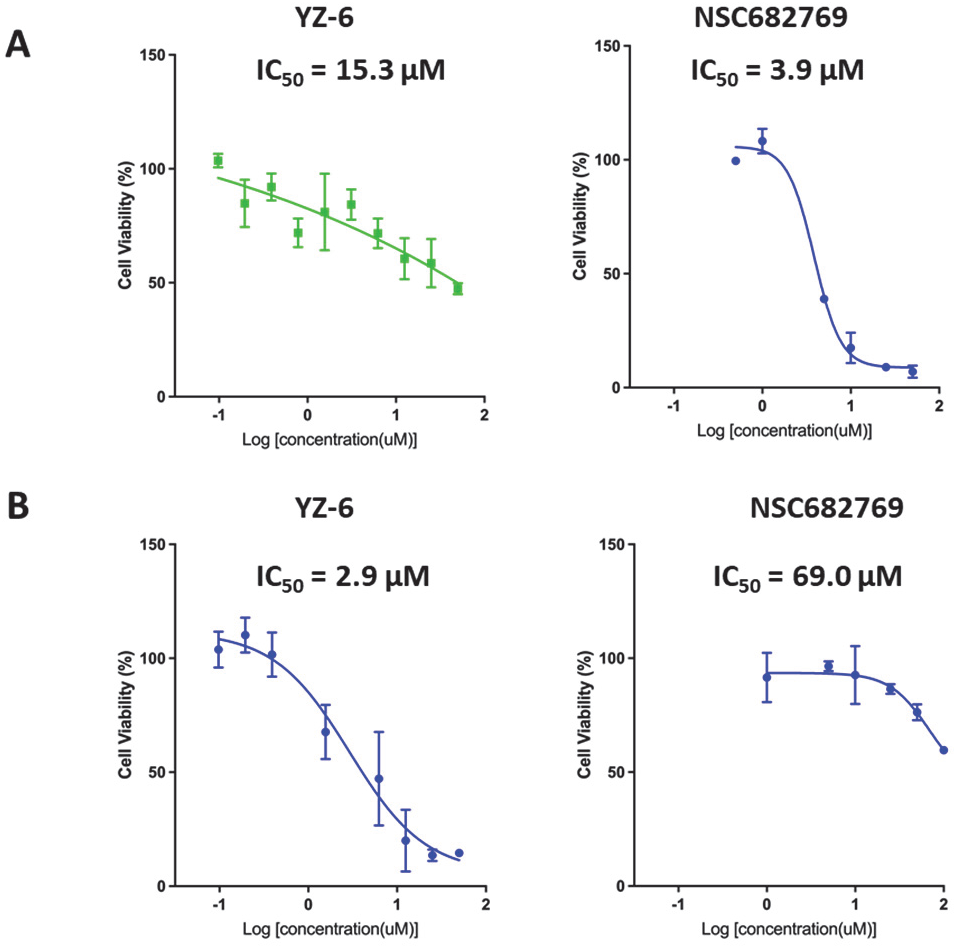
Cell viability assay of YZ-6 and NSC682769 in NCI-H226 and Huh7 cell lines. (A) NCI-H226 cells were treated with the indicated doses of compounds for 4 days. (B) Huh7 cells were treated with indicated doses of two compounds for 4 days. Cell viability was determined by CCK8 assay. IC_50_ values were calculated using the GraphPad Prism 9 software.

Interestingly, when we assessed the growth inhibition activities of **YZ-6** on Huh7 cells that overexpress YAP, it demonstrated significantly superior performance compared to the YAP ligand NSC682769 alone. **YZ-6** exhibited an IC_50_ value of 2.9 μM, making it twenty times more potent than the warhead NSC682769. More importantly, **YZ-6** was able to induce the degradation of YAP with a DC_50_ value of 4.3 μM and effectively inhibit CTGF expression dose dependently in the Huh7 cell line (**Figure 8A**).^53^ These findings indicate that Huh7 cells are more responsive to compound **YZ-6** compared to NCI-H226 cells.

**Figure 8.**
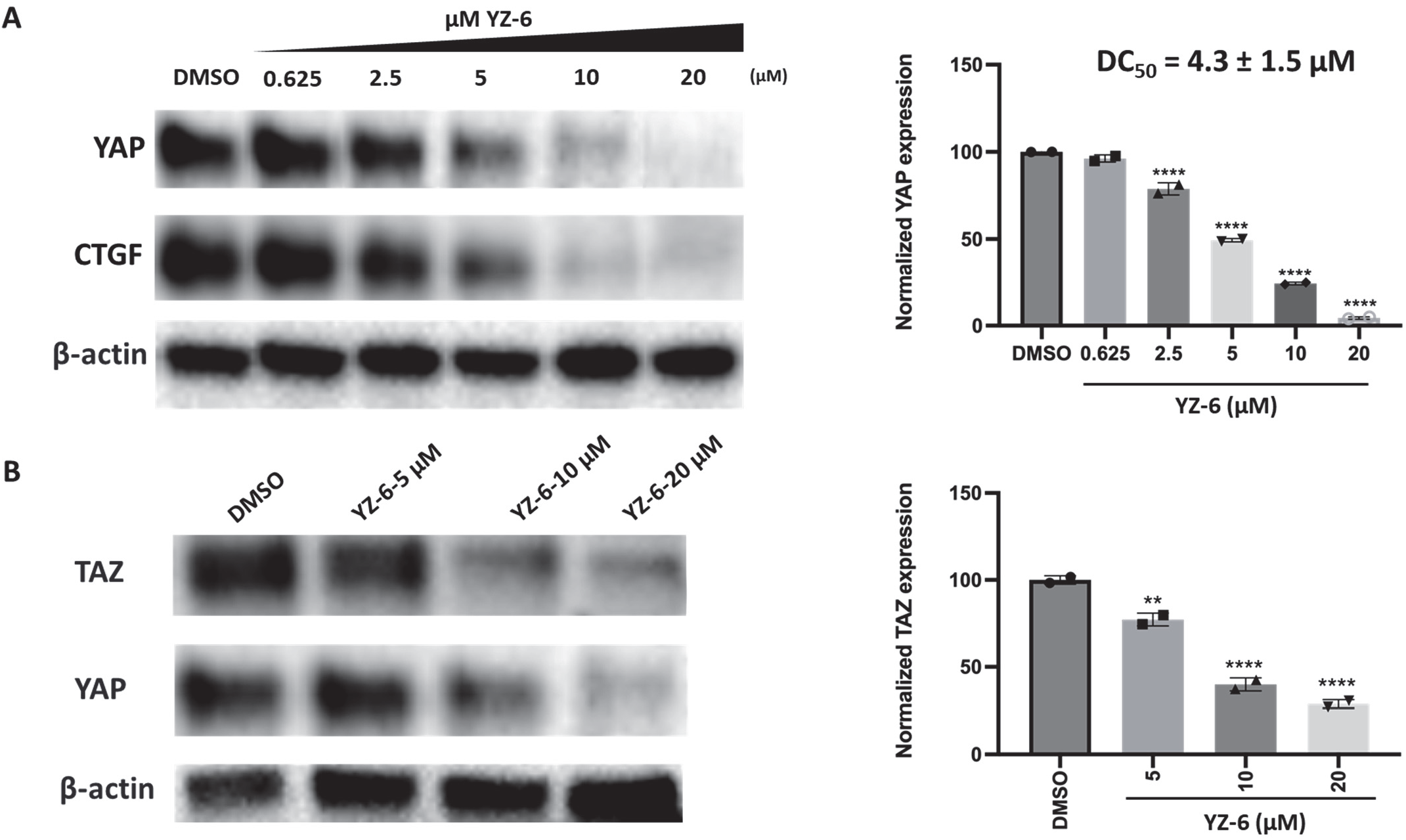
(A) Compound YZ-6 concentration-dependently reduced YAP protein level and inhibit YAP-TEAD target gene CTGF expression. Quantitation on the right. Huh7 cells were treated with DMSO or serial dilutions of compounds for 48 h. (B) Compound YZ-6 concentration-dependently reduced TAZ protein level. Quantitation on the right. Huh7 cells were treated with DMSO or serial dilutions of compounds for 24 h. Quantified data represents mean ± SD from two independent biological replicates. ** p < 0.01; **** p < 0.001.

Since TAZ is a homolog of YAP, we also investigated whether **YZ-6** can also degrade TAZ protein like YAP protein in the Huh7 cell line. The results demonstrated that the expression of TAZ protein was decreased in a dose-dependent manner upon treatment with **YZ-6**, as shown in **Figure 8B**. Thus, **YZ-6** might be able to target cancer cells expressing not only YAP but also TAZ at a high level.

Next, time-course studies were conducted in both NCI-H226 and Huh7 cells (**Figure 9**). YAP was significantly downregulated at 24 h after the treatment with compound **YZ-6** at a dose of 20 μM in both cells. Importantly, YAP could be almost completely degraded by compound **YZ-6** (20 μM) at 48 h in both NCI-H226 and Huh7 cells, demonstrating degradation levels of 97% and 98%, respectively.

**Figure 9.**
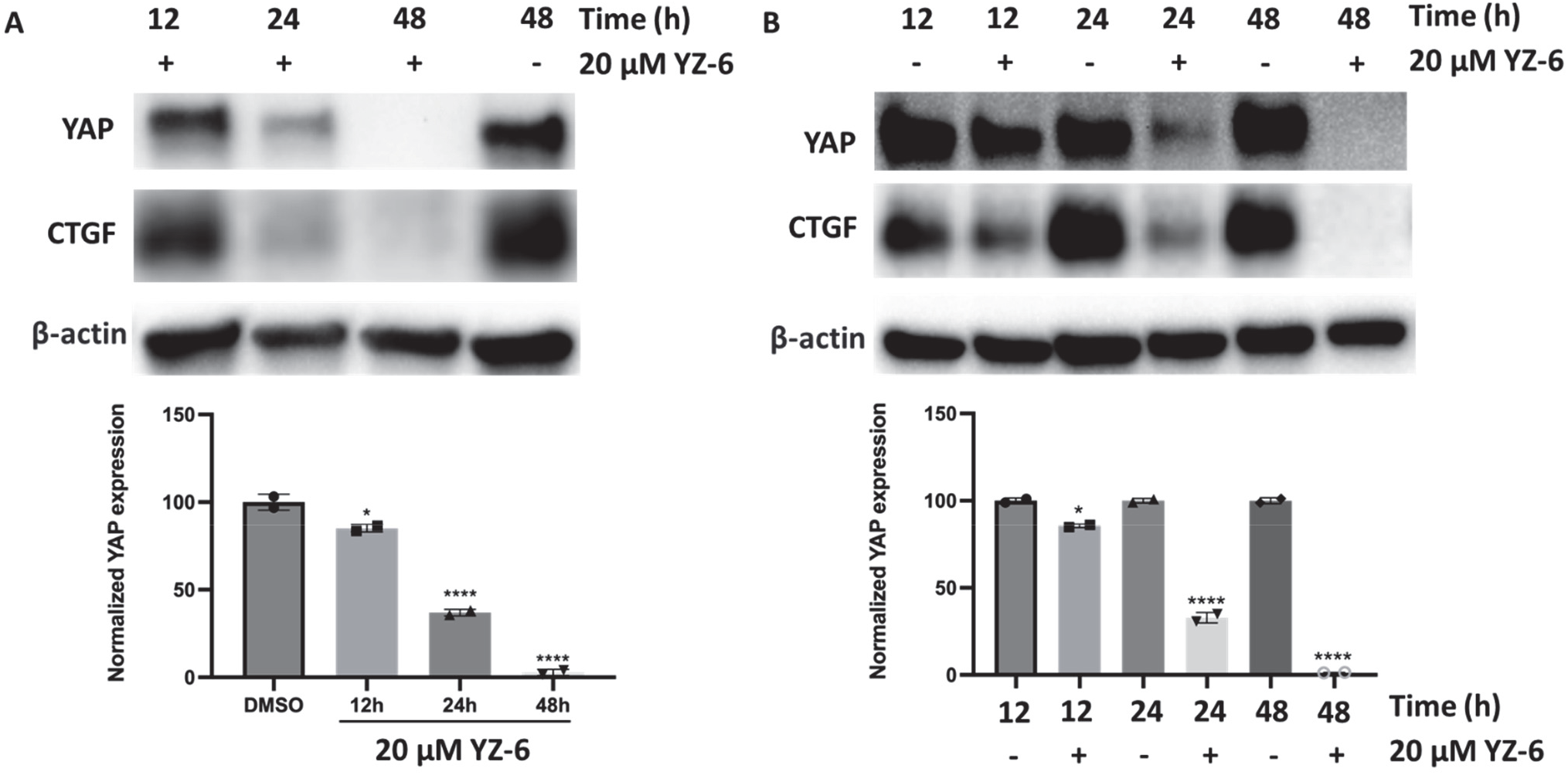
Compound YZ-6 reduces YAP protein levels in a time-dependent manner and inhibited YAP-TEAD target gene CTGF expression. (A) NCI-H226 and (B) Huh7 cells were treated with DMSO or YZ-6 for the indicated time. Quantitation is below. * p < 0.05; **** p < 0.001.

### YZ-6-Induced YAP Degradation Occurs via a *Bona Fide* PROTAC Mechanism

Since excess VHL ligand inhibits ternary complex formation, we performed competition experiments in Huh7 cells that were pretreated for 2 h with molar excess of VHL ligand 2 before being treated with 20 μM **YZ-6**. Competition of **YZ-6** with VHL ligand rescued YAP levels (**Figure 10A**) by preventing PROTAC engagement with VHL.

**Figure 10.**
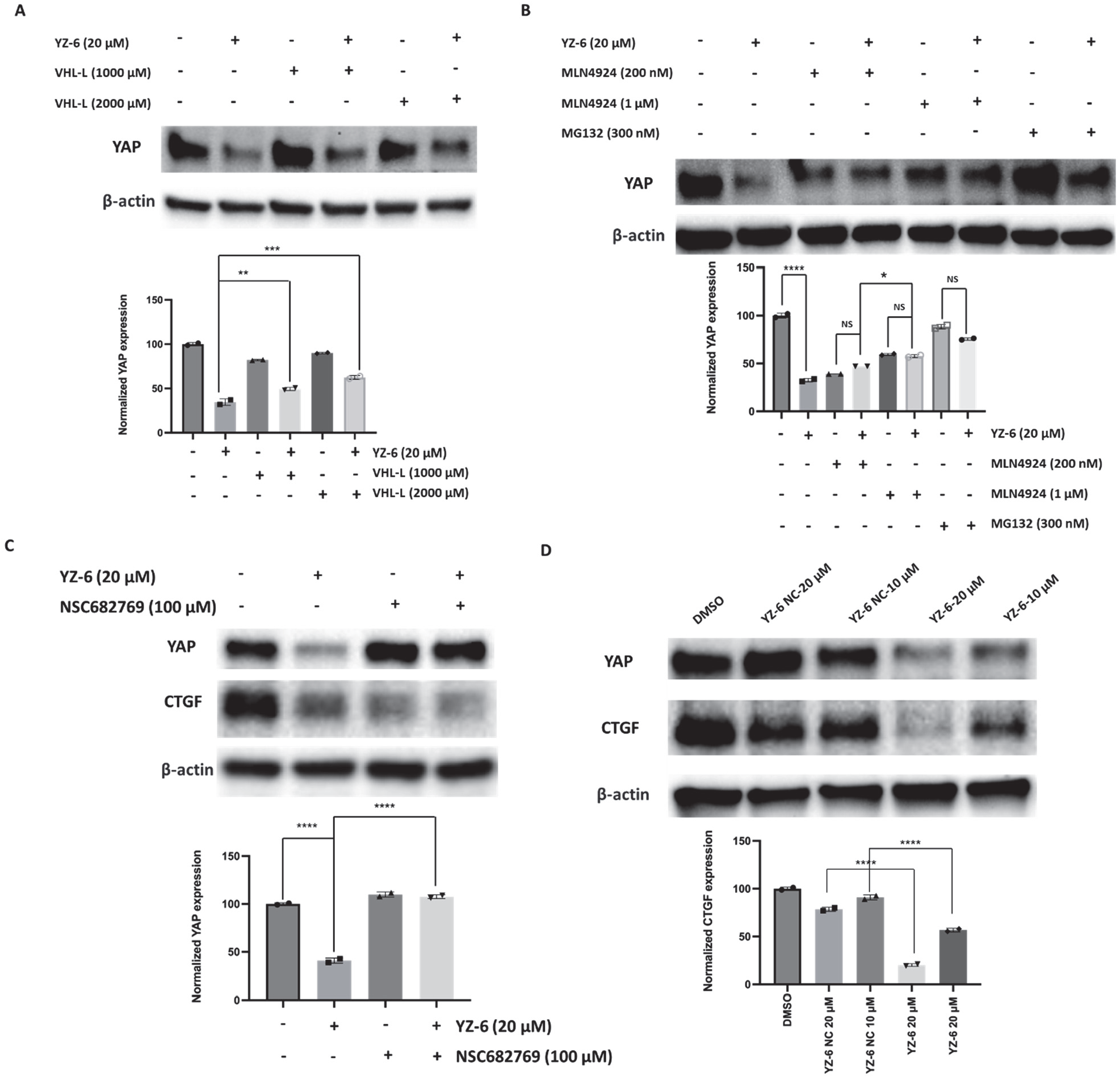
Degradation is dependent on VHL, YAP, and proteasome. (A) Huh7 cells were pretreated with VHL ligand 56 (1 mM) or 2 mM for 2 h, followed by treatment with DMSO or compound YZ-6 (20 μM) for 24 h. Quantitation is below. (B) Huh7 cells were pretreated with 200 nM or 1 μM neddylation inhibitors MLN4924, or the proteasome inhibitor MG132 (300 nM) for 2 h, followed by treatment with DMSO or compound YZ-6 (20 μM) for 24 h. Quantitation is below. (C) Huh7 cells were pretreated with 100 μM YAP inhibitor NSC682769 for 2 h, followed by treatment with DMSO or compound YZ-6 (20 μM) for 24 h. Quantitation is below. (D) Huh7 cells were treated with DMSO, or 10 μM or 20 μM YZ-6, or 10 μM or 20 μM YZ-6 NC for 24 h. Quantitation is below. * p < 0.05; ** p < 0.01; *** p < 0.005; **** p < 0.001.

To investigate whether compound **YZ-6** mediated the degradation of YAP through the corresponding E3 ligase and UPS, Huh7 cells were cotreated with compound **YZ-6** and MG-132 (a proteasome inhibitor) or MLN-4924 (an NEDD8-activating enzyme (NAE) inhibitor).^54, 55^ Both MG-132 and MLN-4924 rescued the YAP protein level in Huh7 cells (**Figure 10B**), suggesting that YAP degradation by **YZ-6** is both proteasome-and neddylation-dependent.

As *bona fide* PROTAC degrader molecule, depletion of YAP protein by **YZ-6** should require their binding to YAP through their **NSC682769** segment.^56^ Accordingly, we predicted that excessive amounts of **NSC682769** should effectively block the YAP degradation induced by **YZ-6**. Western blotting analysis showed that pretreatment with 100 μM **NSC682769** indeed effectively prevent YAP from degrading by **YZ-6** in Huh7 cell line (**Figure 10C**).

The hydroxy proline moiety of the VHL ligand confers binding to the E3 ligase, while inversion of the absolute stereochemistry of the 4-hydroxy proline moiety abrogates VHL binding.^57^ Therefore, we synthesized **YZ-6 NC** as a physicochemically matched negative control molecule that is unable to recruit VHL. When Huh7 cells were treated with 10 μM and 20 μM **YZ-6** NC for 24 h, no YAP degradation was observed, whereas 10 μM and 20 μM **YZ-6** induced significant degradation (∼50% and 78%, respectively; **Figure 10D**). The effect of **YZ-6** and **YZ-6 NC** on the YAP/TEAD downstream transcriptional marker CTGF was also assessed. The significantly more potent suppression of CTGF expression by **YZ-6** compared to **YZ-6 NC** is indicative of the additional degradation activity afforded by **YZ-6**, while **YZ-6 NC** only exerted an antagonist-induced effect.

Taken together, these data show that **YZ-6**-induced YAP degradation is dependent on ternary complex formation with VHL and a functioning ubiquitin proteasome system.

For tethering site 3 (**Figure 11**), we also synthesized 7 compounds, incorporating aliphatic or ether linkers connected to either VHL ligand or CRBN ligand pomalidomide. Since site 3 is not projected to be solvent exposed, those compounds will not bind to YAP, resulting in the loss of YAP degradation efficacy. As expected, none of the compounds displayed YAP degradation and NCI-H226 cell killing activity, proving our preliminary hypothesis that only attaching the linker at the solvent-exposed region could have the potential to induce YAP degradation.

**Figure 11.**
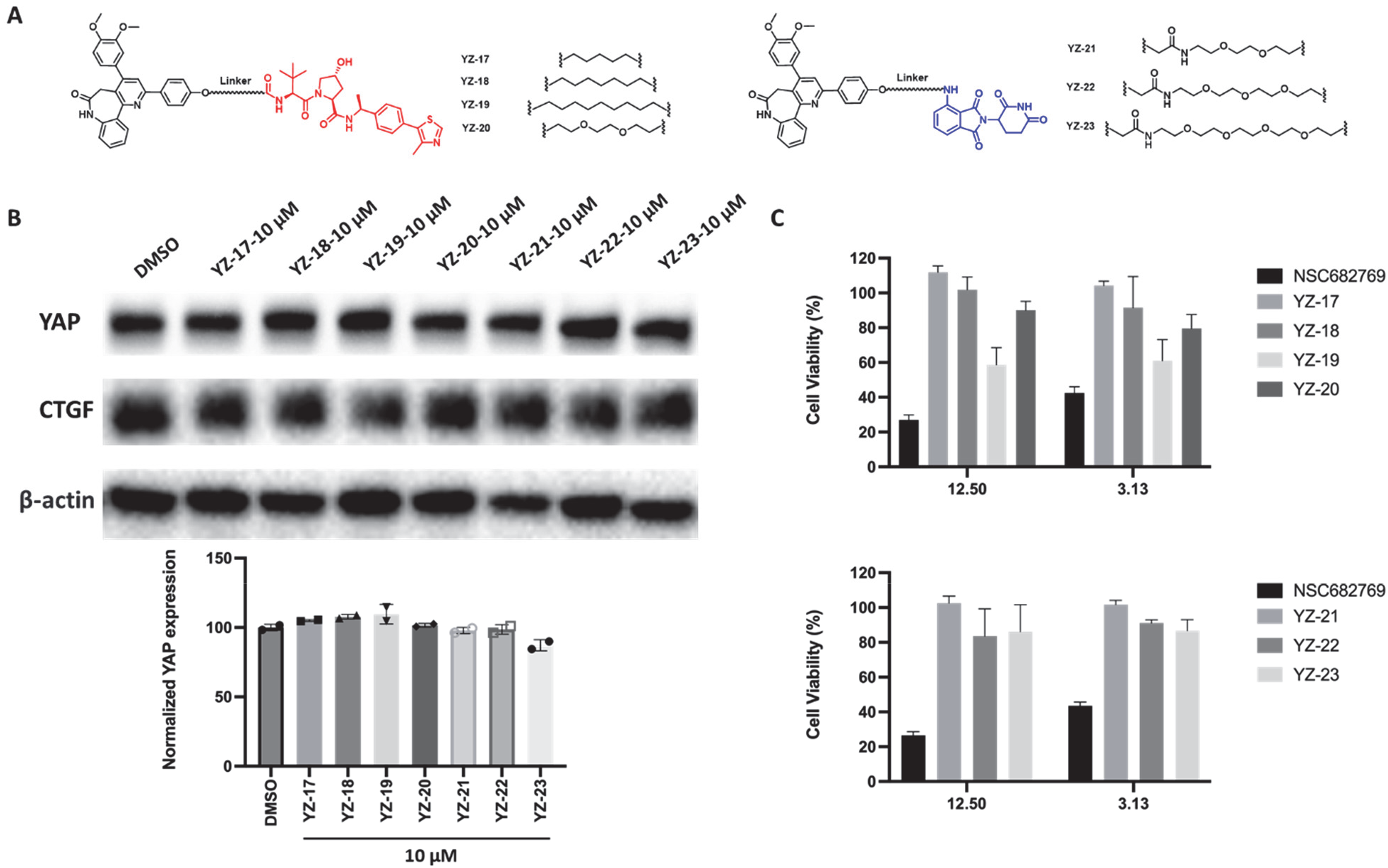
(A) Chemical structure of designed NSC682769-based YAP degraders using tethering site 3. (B) Western blotting analysis of YAP protein in NCI-H226 cells treated with compounds YZ-17-23. Quantitation is below. (C) Cell viability assay. NCI-H226 cells were treated with indicated doses of compounds for 4 days.

### Antitumor Activity of YZ-6 in Mice

Utilizing its promising in vitro potency in the Huh7 cell line as a starting point, we proceeded to assess the in *vivo* effectiveness of **YZ-6** in inhibiting tumor growth in a Huh7 xenograft model in mice. The mice received 35 mg/kg of **YZ-6** every three days for a total of 24 days through intraperitoneal injection (IP). As illustrated in **Figure 12A** and **12B**, **YZ-6** exhibited significant antitumor efficacy, achieving a tumor growth inhibition (TGI) rate of 82%. Remarkably, there was no notable change in the animals’ body weight, as depicted in **Figure 12C**, suggesting that **YZ-6** treatment did not induce apparent toxicity in the xenograft model. These findings demonstrate **YZ-6**’s capability to effectively suppress Huh7 xenograft tumor growth in *vivo*.

**Figure 12.**
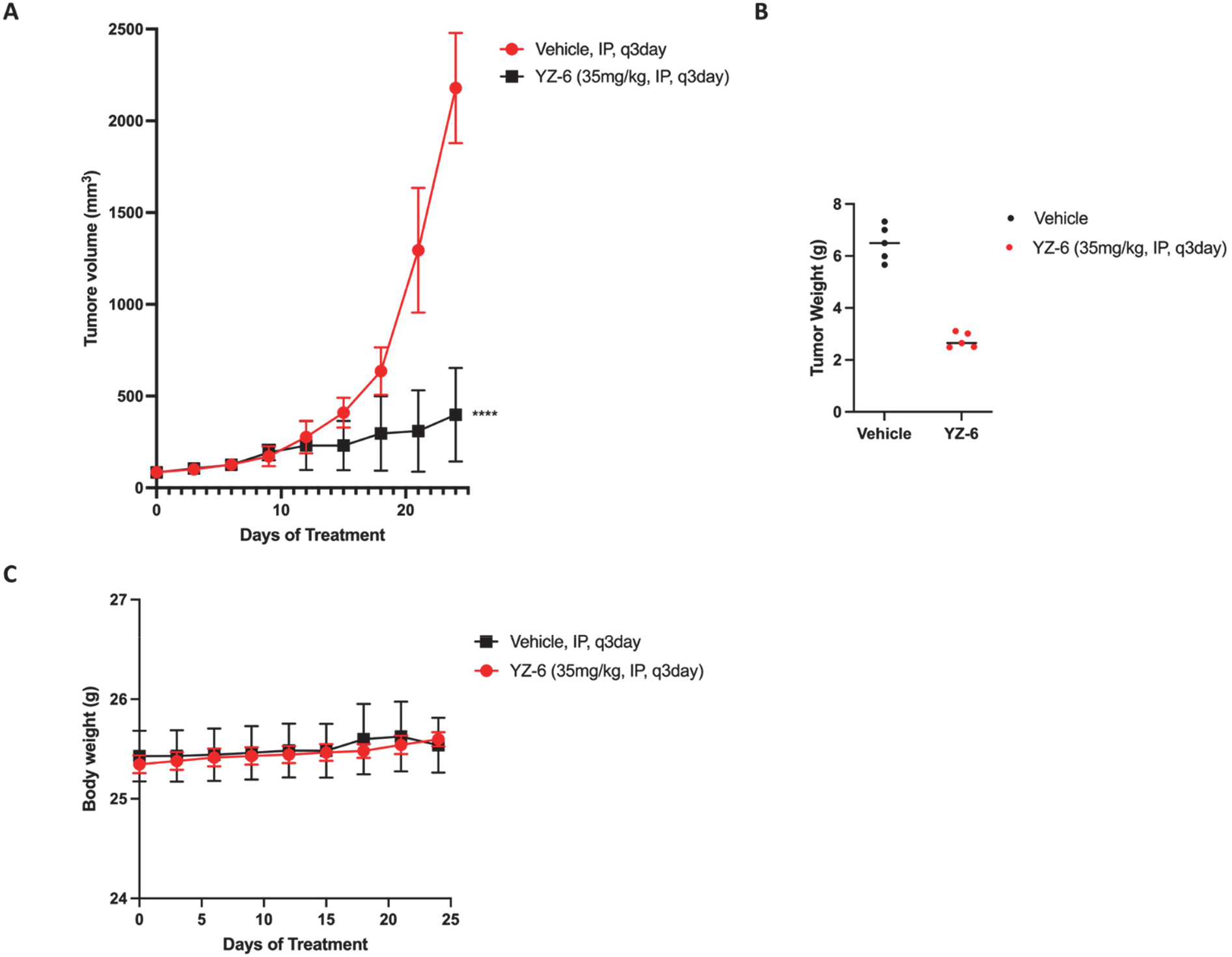
*In vivo* antitumor activity of compound YZ-6 in an Huh7 cell-derived xenograft. Tumor volume (A), tumor weight (B), and body weight (C) of the mice after treatment of compound YZ-6, n = 5 mice per group. ∗∗∗∗p < 0.0001.

### Chemistry

The preparation of compounds **6 (NSC682769)** and **7** is illustrated in Scheme 1.^58^ Compound **1** acetophenone underwent aldol condensation with dimethoxybenzaldehyde in the presence of sodium hydroxide to yield compound **3**. The synthesis of compound **5** was accomplished via a Michael addition with 3,4-dihydro-1H-benzo[b]azepine-2,5-dione catalyzed by 0.1 equivalent of KOH. Subsequent cyclization by means of ammonium ferric sulfate in glacial acetic acid yielded the desired pyrido[3,2- d][1]benzazepin-6-one compound **6**. Ethylation at the nitrogen of lactam 6 was carried out with sodium hydride and bromoethane to give compound **7**.

### Scheme 1

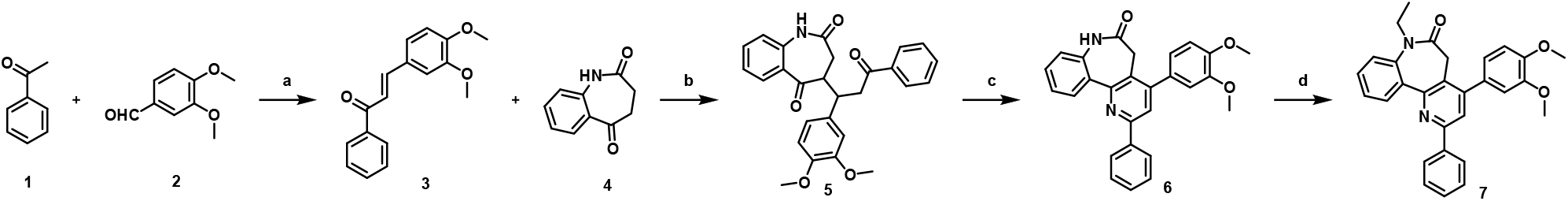

Scheme 1. Synthesis of Compounds **6** and **7**^a^

^a^Reagents and conditions: (a) NaOH, EtOH, H_2_O, rt, 18h, 83%; (b) KOH, EtOH, H_2_O, rt, 12h, 76%; (c) NH_4_Fe(SO_4_)_2_ ⋅ 12H_2_O, NH_4_OAc, AcOH, reflux, 4h, 85%; (d) bromoethane, NaH, KI, THF, rt, 24h, 55%.

The synthetic route for preparing compounds **12**, **13** and **14** is outlined in Scheme 2. Compound **1** acetophenone underwent aldol condensation with 4-(benzyloxy)-3-methoxybenzaldehyde in the presence of sodium hydroxide to give compound **9**. The synthesis of compound **10** was accomplished via a Michael addition with 3,4-dihydro-1H-benzo[b]azepine-2,5-dione catalyzed by 0.1 equivalent of KOH. Subsequent cyclization by means of ammonium ferric sulfate in glacial acetic acid yielded the desired pyrido[3,2-d][1]benzazepin-6-one compound **11**. Deprotection via palladium-catalyzed hydrogenolysis afforded the corresponding phenol **12**. The hydroxyl was then subjected to an S_N_2 reaction with 2-bromoethyl acetate to give the corresponding ether compound **13**, which was subsequently treated with potassium carbonate to produce the compound **14**.

### Scheme 2

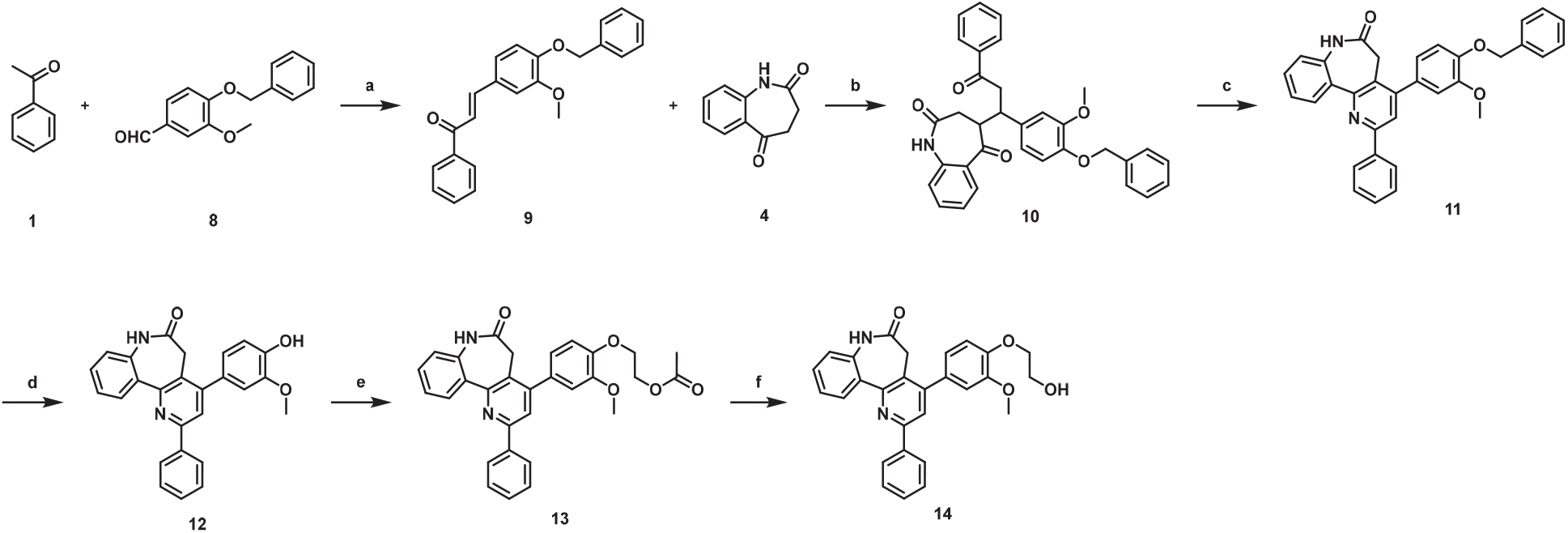

Scheme 2. Synthesis of Compounds **12**, **13** and **14**^a^

^a^Reagents and conditions: (a) NaOH, EtOH, H_2_O, rt, 18h, 81%; (b) KOH, EtOH, H_2_O, rt, 12h, 72%; (c) NH_4_Fe(SO_4_)_2_ ⋅ 12H_2_O, NH_4_OAc, AcOH, reflux, 4h, 82%; (d) Pd/C, H_2_, CH_3_OH, 60 °C, 12h, 90%; (e) 2-bromoethyl acetate, K_2_CO_3_, DMF, 50 °C, 10h, 65%; (f) K_2_CO_3_, CH_3_OH, rt, 6h, 95%.

The synthetic route for preparing compounds **19**, **20** and **21** is outlined in Scheme 3. Compound **15** 1-(4-(benzyloxy)phenyl)ethan-1-one underwent aldol condensation with 3,4-dimethoxybenzaldehyde in the presence of sodium hydroxide to give compound **16**. The synthesis of compound **17** was accomplished via a Michael addition with 3,4-dihydro-1H-benzo[b]azepine-2,5-dione catalyzed by 0.1 equivalent of KOH (crude for next step). Subsequent cyclization by means of ammonium ferric sulfate in glacial acetic acid yielded the desired pyrido[3,2-d][1]benzazepin-6-one compound **18** (crude for next step). Deprotection via palladium-catalyzed hydrogenolysis afforded the corresponding phenol **19**. The hydroxyl was then subjected to an S_N_2 reaction with 2- bromoethyl acetate to give the corresponding ether compound **20**, which was subsequently treated with potassium carbonate to produce compound **21**.

### Scheme 3

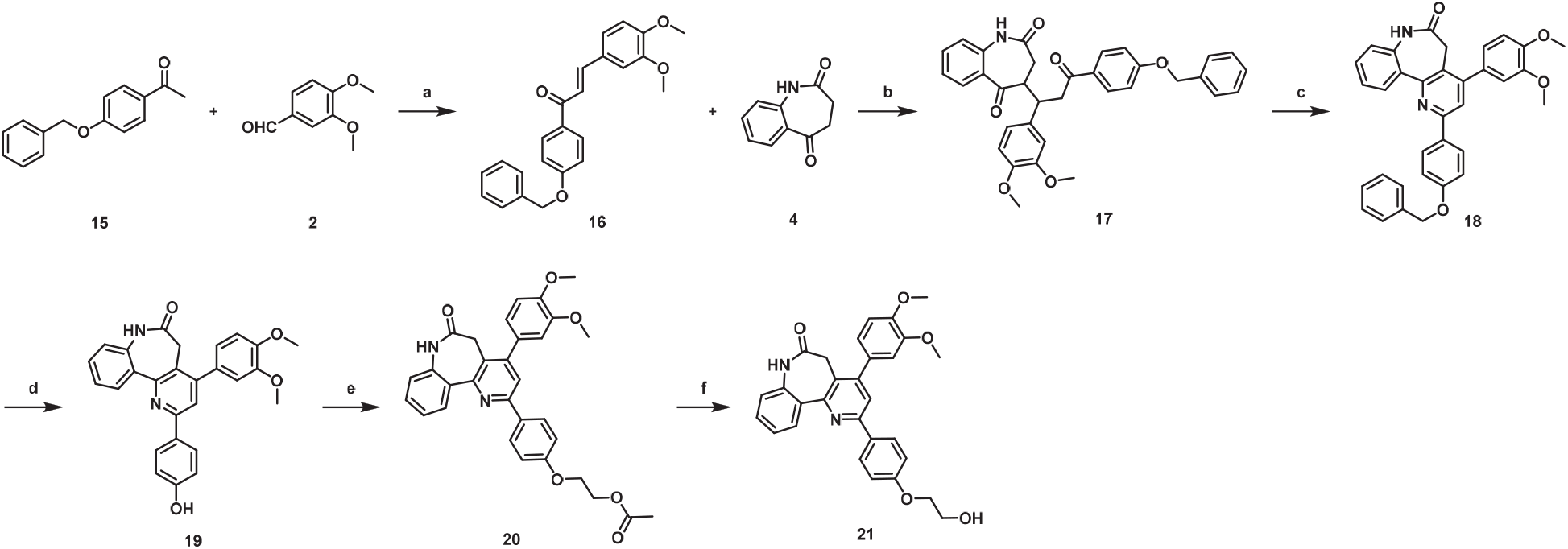

Scheme 3. Synthesis of Compounds **19**, **20** and **21**^a^

^a^Reagents and conditions: (a) NaOH, EtOH, H_2_O, rt, 18h, 87%; (b) KOH, EtOH, H_2_O, rt, 12h; (c) NH_4_Fe(SO_4_)_2_ ⋅ 12H_2_O, NH_4_OAc, AcOH, reflux, 4h; (d) Pd/C, H_2_, CH_3_OH, 60 °C, 12h, 58% (3 steps); (e) 2-bromoethyl acetate, K_2_CO_3_, DMF, 50 °C, 10h, 60%; (f) K_2_CO_3_, CH_3_OH, rt, 6h, 94%.

The synthetic route for preparing compounds **25**, **26** and **27** is outlined in Scheme 4. Compound **1** acetophenone underwent aldol condensation with 3-(benzyloxy)-4-methoxybenzaldehyde in the presence of sodium hydroxide to give compound **23**. The synthesis of compound **24** was accomplished via a Michael addition with 3,4-dihydro-1H-benzo[b]azepine-2,5-dione catalyzed by 0.1 equivalent of KOH. Subsequent cyclization by means of ammonium ferric sulfate in glacial acetic acid yielded the desired pyrido[3,2-d][1]benzazepin-6-one compound **25**. Deprotection via palladium-catalyzed hydrogenolysis afforded the corresponding phenol **26**. The hydroxyl was then subjected to an S_N_2 reaction with 2-bromoethyl acetate to give the corresponding ether compound **27.**

### Scheme 4

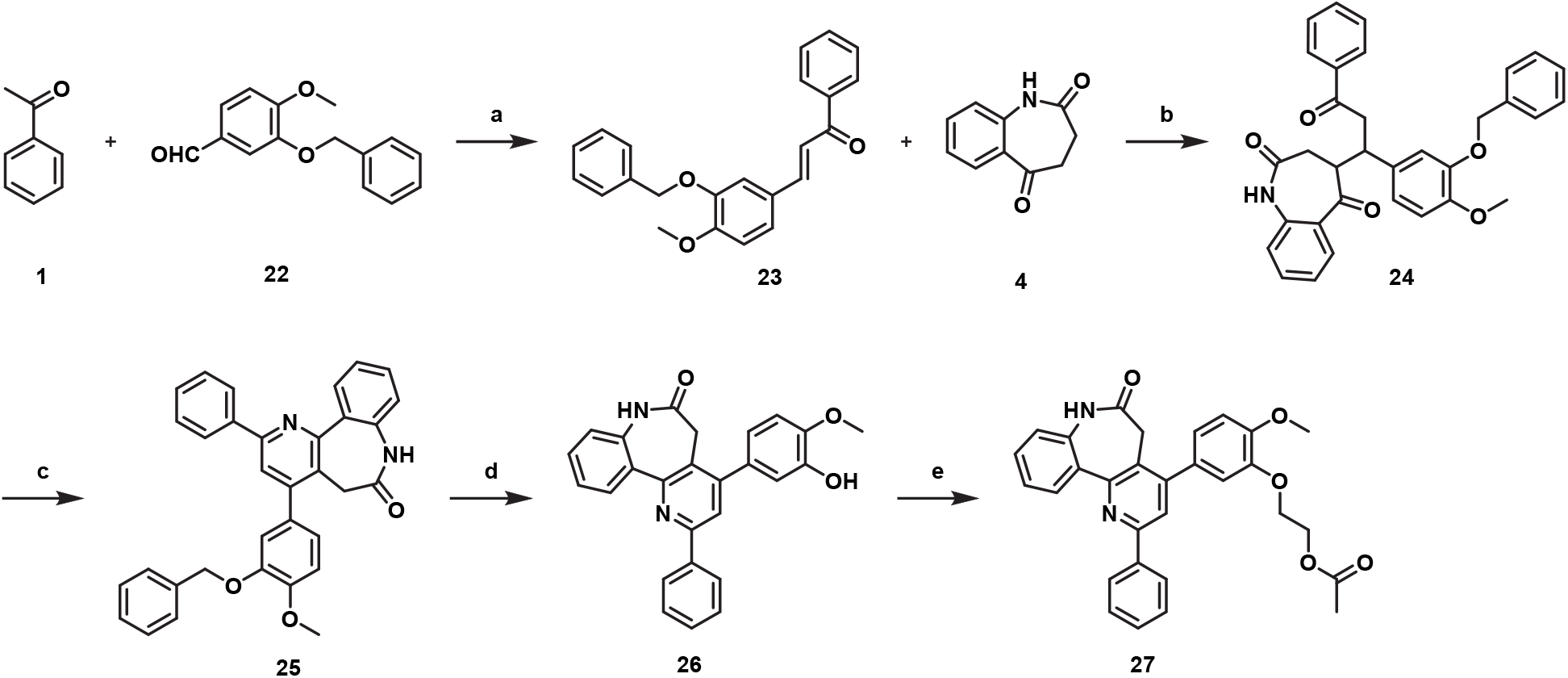

Scheme 4. Synthesis of Compounds **25**, **26** and **27**^a^

^a^Reagents and conditions: (a) NaOH, EtOH, H_2_O, rt, 18h, 78%; (b) KOH, EtOH, H_2_O, rt, 12h; (c) NH_4_Fe(SO_4_)_2_ ⋅ 12H_2_O, NH_4_OAc, AcOH, reflux, 4h, 56% (2 steps); (d) Pd/C, H_2_, CH_3_OH, 60 °C, 12h, 83%; (e) 2-bromoethyl acetate, K_2_CO_3_, DMF, 50 °C, 10h, 61%.

The synthetic route for preparing compound **32** is outlined in Scheme 5. The compound **28** was reacted with *p*-toluenesulfonyl chloride to give a sulfonate ester **29**. The *t*-butyl group of **29** was cleaved via TFA to afford compound **30**. Then, coupling with commercially available compound **31** in the presence of HATU and DIPEA provided the final compound **32**.

### Scheme 5

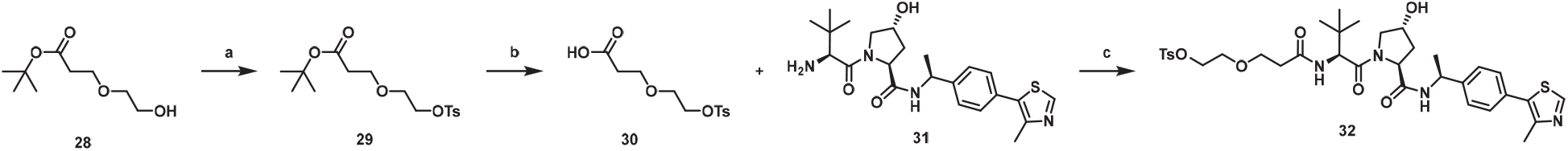

Scheme 5. Synthesis of Compound **32**^a^

^a^Reagents and conditions: (a) p-toluenesulfonyl chloride, NEt_3_, DMAP, DCM, rt, 8h, 82%; (b) TFA, DCM, rt, 2h, 95%; (c) HATU, DIPEA, DMF, rt, 8h, 74%.

The synthetic route for preparing compound **35** is outlined in Scheme 6. The *t*-butyl group of **33** was cleaved via TFA to afford compound **34**. Then, coupling with commercially available compound **31** in the presence of HATU and DIPEA provided the final compound **35**.

### Scheme 6

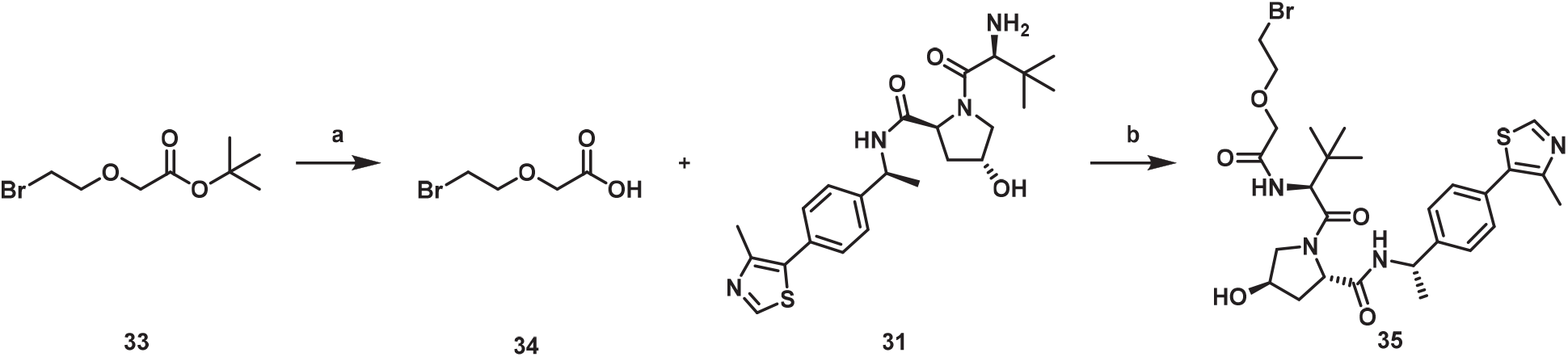

Scheme 6. Synthesis of Compound **35**^a^

^a^Reagents and conditions: (a) TFA, DCM, rt, 2h, 95%; (c) HATU, DIPEA, DMF, rt, 8h, 70%.

The synthetic route for preparing compounds **40** and **41** is outlined in Scheme 7. The *t*-butyl groups of **36** and **37** were cleaved via TFA to afford corresponding compounds **38** and **39**. Then, coupling with commercially available compound **31** in the presence of HATU and DIPEA provided the final compounds **40** and **41**.

### Scheme 7

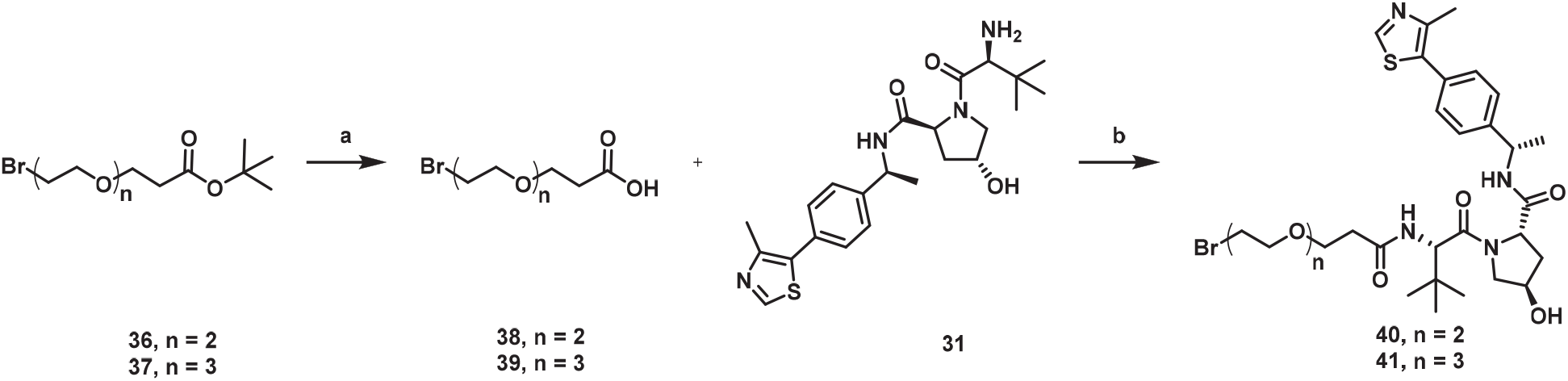

Scheme 7. Synthesis of Compounds **40** and **41**^a^

^a^Reagents and conditions: (a) TFA, DCM, rt, 2h, 91%-93%; (c) HATU, DIPEA, DMF, rt, 8h, 55%-62%.

The synthetic route for preparing compounds **YZ-1, YZ-2** and **YZ-3** is outlined in Scheme 8. The hydroxyl of compound **12** was subjected to an S_N_2 reaction with compounds **42**, **43** and **44** to give corresponding ether compounds **YZ-1**, **YZ-2** and **YZ-3**.

### Scheme 8

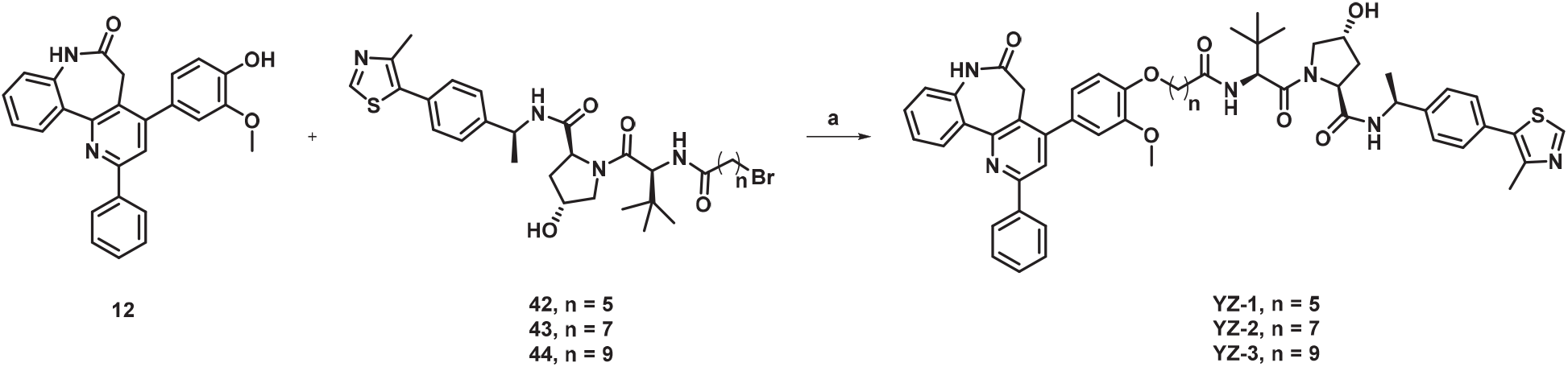

Scheme 8. Synthesis of Compounds **YZ-1**, **YZ-2** and **YZ-3**^a^

^a^Reagents and conditions: (a) K_2_CO_3_, KI, DMF, rt, 12h, 41% - 55%.

The synthetic route for preparing compound **YZ-4** is outlined in Scheme 9. The hydroxyl of compound **12** was subjected to an S_N_2 reaction with compound **35** to give the corresponding ether compound **YZ-4**.

### Scheme 9

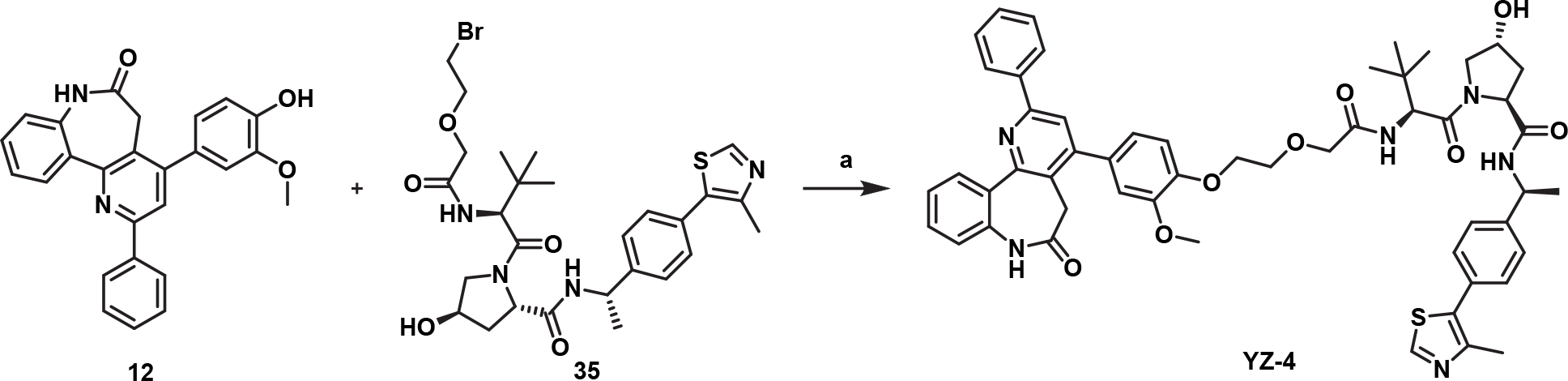

Scheme 9. Synthesis of Compound **YZ-4**^a^

^a^Reagents and conditions: (a) K_2_CO_3_, KI, DMF, rt, 12h, 45%.

The synthetic route for preparing compound **YZ-5** is outlined in Scheme 10. The hydroxyl of compound **12** was subjected to an S_N_2 reaction with compound **32** to give the corresponding ether compound **YZ-5**.

### Scheme 10

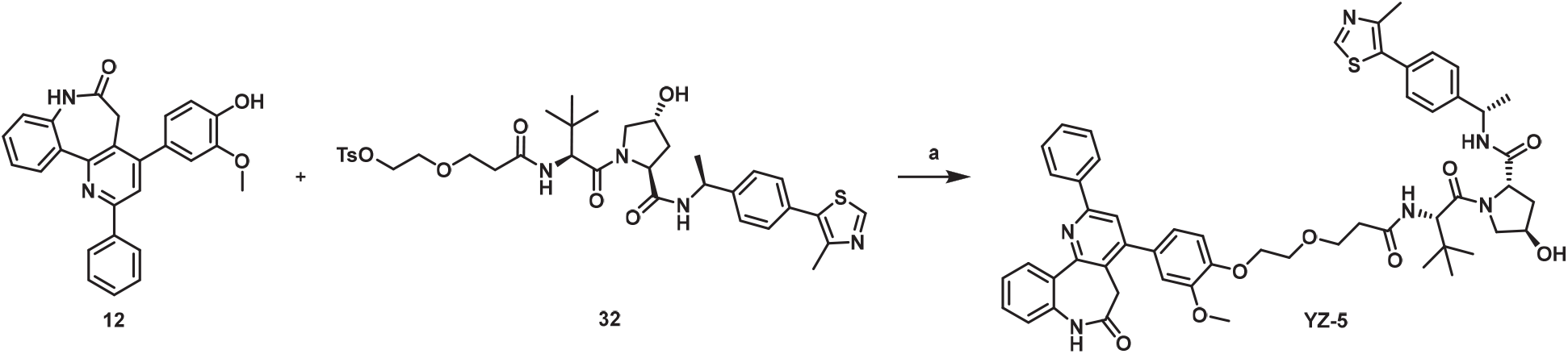

Scheme 10. Synthesis of Compound **YZ-5**^a^

^a^Reagents and conditions: (a) K_2_CO_3_, KI, DMF, rt, 12h, 49%.

The synthetic route for preparing compounds **YZ-6** and **YZ-7** is outlined in Scheme 11. The hydroxyl of compound **12** was subjected to an S_N_2 reaction with compounds **40** and **41** to give corresponding ether compounds **YZ-6** and **YZ-7.**

### Scheme 11

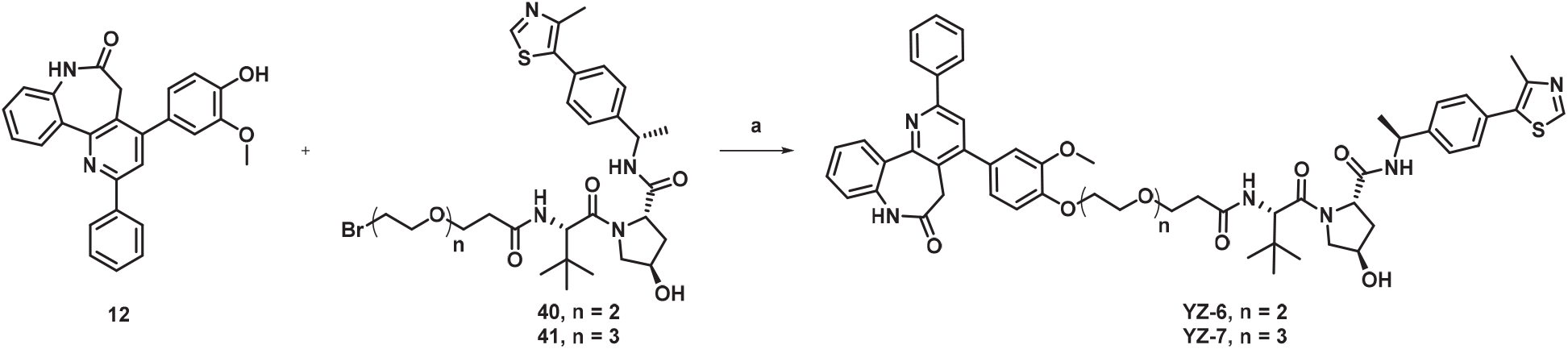

Scheme 11. Synthesis of Compounds **YZ-6** and **YZ-7**^a^

^a^Reagents and conditions: (a) K_2_CO_3_, KI, DMF, rt, 12h, 53%-58%.

The synthetic route for preparing compound **YZ-8** is outlined in Scheme 12. The hydroxyl of compound **26** was subjected to an S_N_2 reaction with compound **35** to give corresponding ether compound **YZ-8**.

### Scheme 12

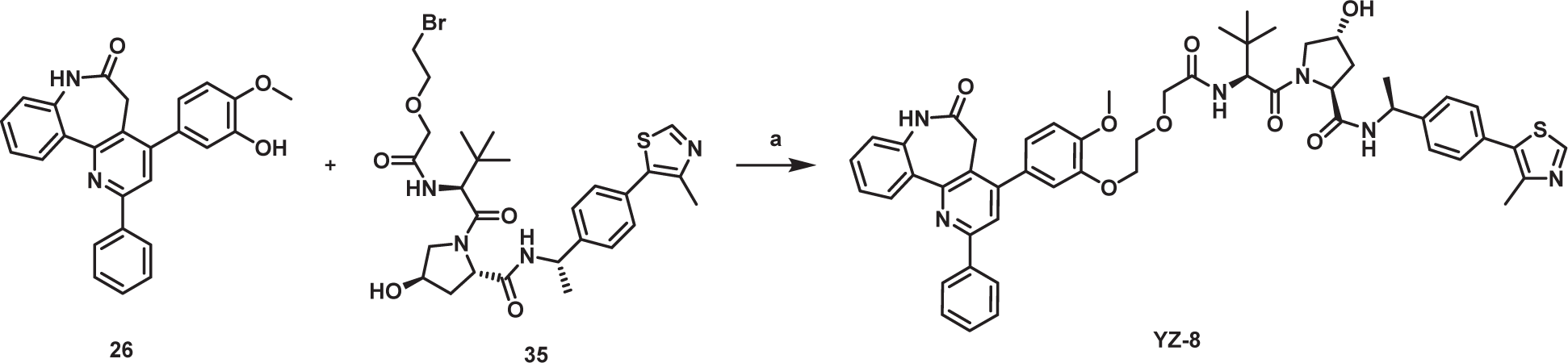

Scheme 12. Synthesis of Compound **YZ-8**^a^

^a^Reagents and conditions: (a) K_2_CO_3_, KI, DMF, rt, 12h, 41%.

The synthetic route for preparing compound **YZ-9** is outlined in Scheme 13. The hydroxyl of compound **26** was subjected to an S_N_2 reaction with compound **32** to give corresponding ether compound **YZ-9**.

### Scheme 13

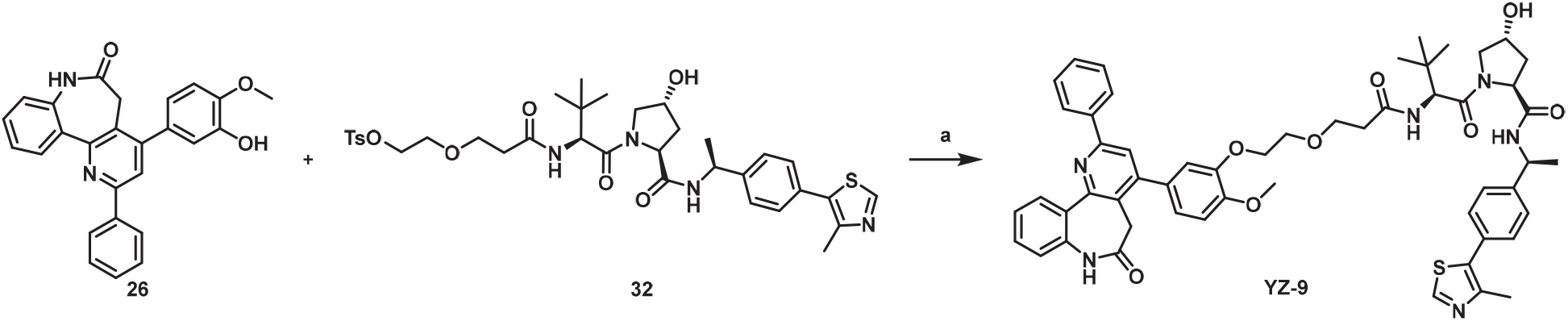

Scheme 13. Synthesis of Compound **YZ-9**^a^

^a^Reagents and conditions: (a) K_2_CO_3_, KI, DMF, rt, 12h, 48%.

The synthetic route for preparing compound **YZ-10** is outlined in Scheme 14. The hydroxyl of compound **26** was subjected to an S_N_2 reaction with compound **40** to give corresponding ether compound **YZ-10**.

### Scheme 14

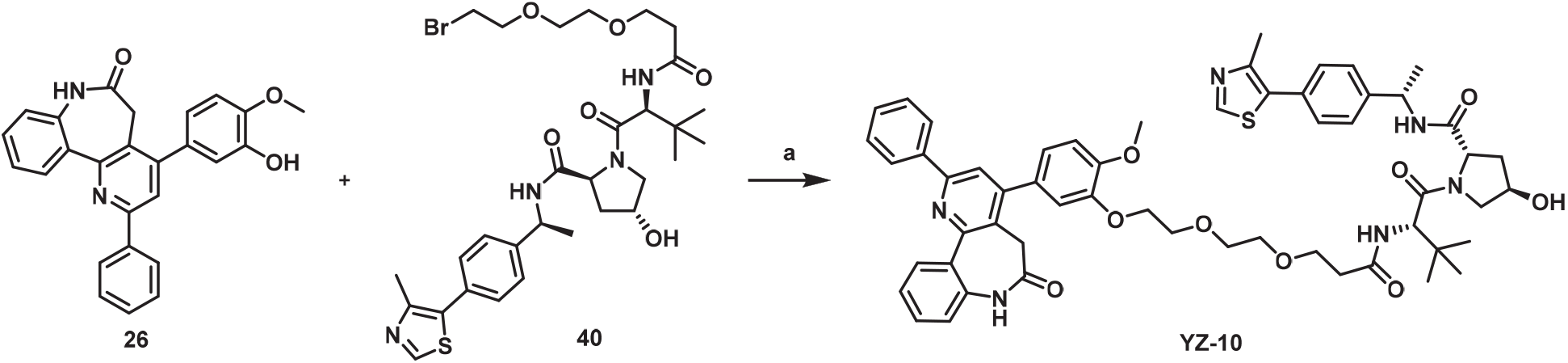

Scheme 14. Synthesis of Compound **YZ-10**^a^

^a^Reagents and conditions: (a) K_2_CO_3_, KI, DMF, rt, 12h, 55%.

The synthetic route for preparing compound **46** is outlined in Scheme 15. The hydroxyl of compound **12** was subjected to an S_N_2 reaction with *tert*-butyl 2-bromoacetate to give corresponding ether compound **45** (crude for next step). The *t*-butyl group of **45** was cleaved via TFA to afford compound **46**.

### Scheme 15

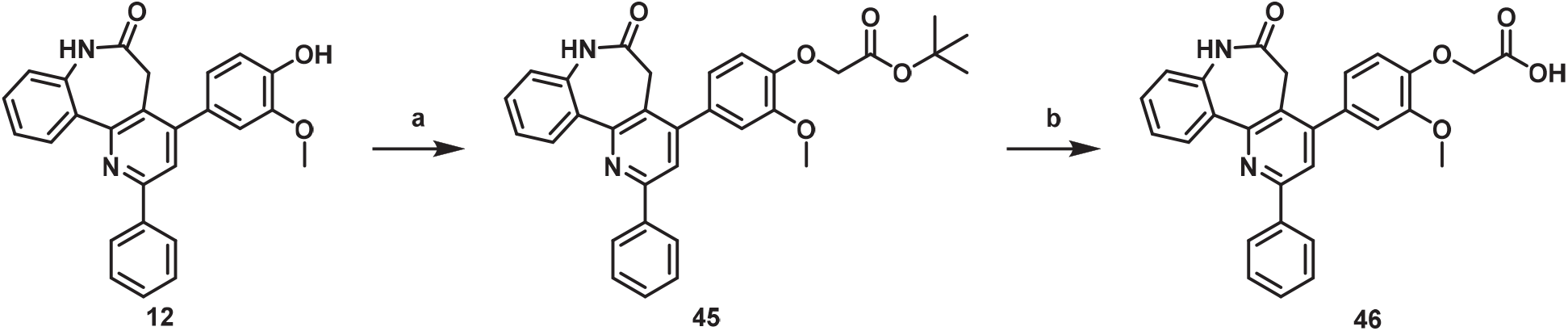

Scheme 15. Synthesis of Compound **46**^a^

^a^Reagents and conditions: (a) *tert*-butyl 2-bromoacetate, K_2_CO_3_, DMF, rt, 12h. (b) TFA, DCM, rt, 2h, 56% (2 steps).

The synthetic route for preparing compound **48** is outlined in Scheme 16. The hydroxyl of compound **26** was subjected to an S_N_2 reaction with *tert*-butyl 2-bromoacetate to give corresponding ether compound **47**. The *t*-butyl group of **47** was cleaved via TFA to afford compound **48**.

### Scheme 16

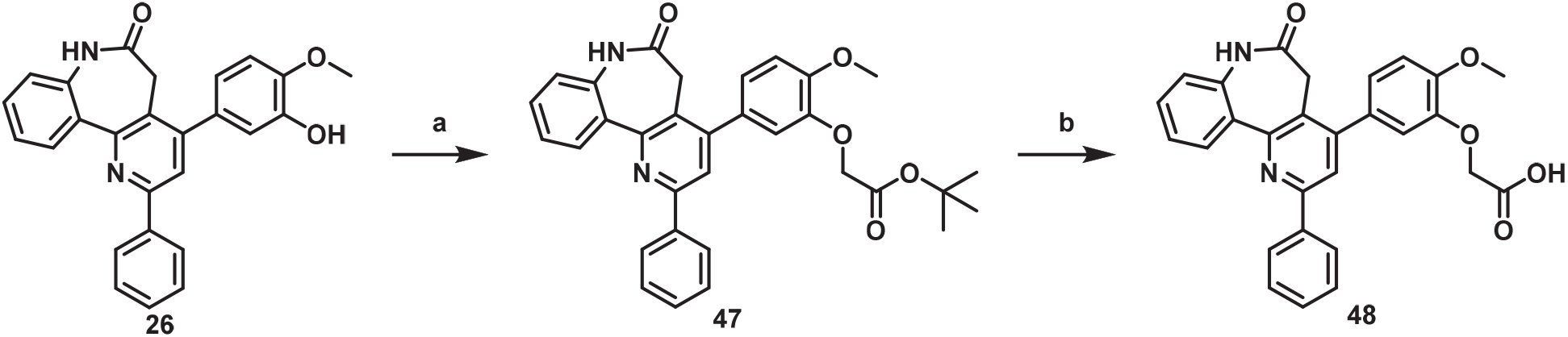

Scheme 16. Synthesis of Compound **48**^a^

^a^Reagents and conditions: (a) *tert*-butyl 2-bromoacetate, K_2_CO_3_, DMF, rt, 12h, 60%. (b) TFA, DCM, rt, 2h, 93%.

The synthetic route for preparing compounds **YZ-11**, **YZ-12** and **YZ-13** is outlined in Scheme 17. Compound **46** was coupled with commercially available compounds **49**, **50** and **51** in the presence of HATU and DIPEA provided the corresponding amide compounds **YZ-11**, **YZ-12** and **YZ-13**.

### Scheme 17

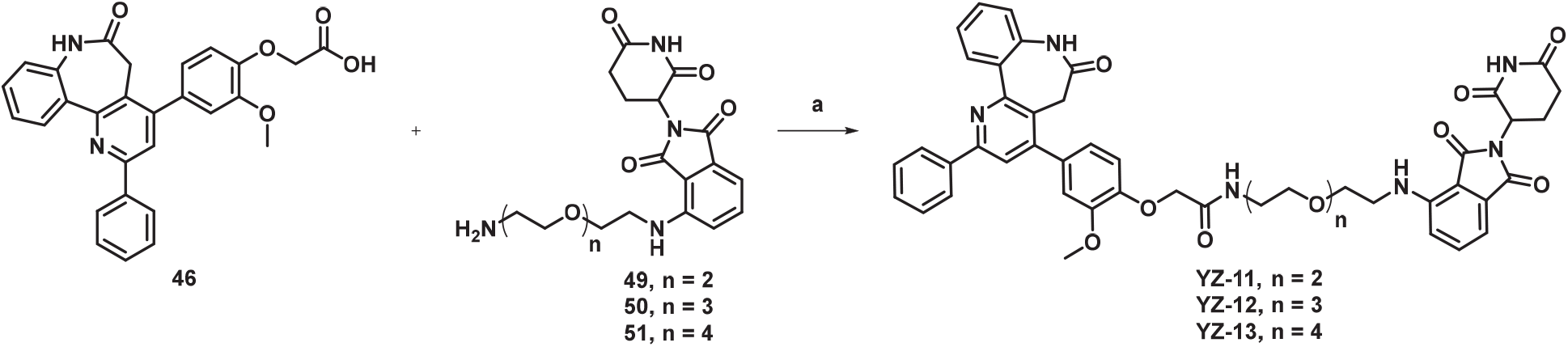

Scheme 17. Synthesis of Compounds **YZ-11**, **YZ-12** and **YZ-13**^a^

^a^Reagents and conditions: (a) HATU, DIPEA, DMF, rt, 16h, 50%-65%.

The synthetic route for preparing compounds **YZ-14**, **YZ-15** and **YZ-16** is outlined in Scheme 18. Compound **48** was coupled with commercially available compounds **49**, **50** and **51** in the presence of HATU and DIPEA provided the corresponding amide compounds **YZ-14**, **YZ-15** and **YZ-16**.

### Scheme 18

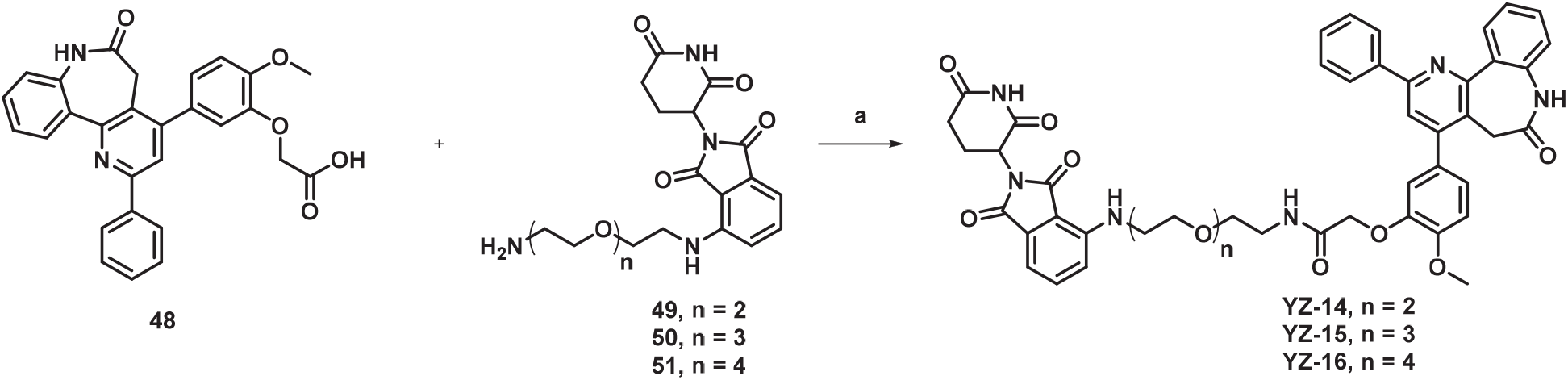

Scheme 18. Synthesis of Compounds **YZ-14**, **YZ-15** and **YZ-16**^a^

^a^Reagents and conditions: (a) HATU, DIPEA, DMF, rt, 16h, 62%-73%.

The synthetic route for preparing compound **53** is outlined in Scheme 19. Compound **38** was coupled with commercially available compound **52** in the presence of HATU and DIPEA provided the final compound **53**.

### Scheme 19

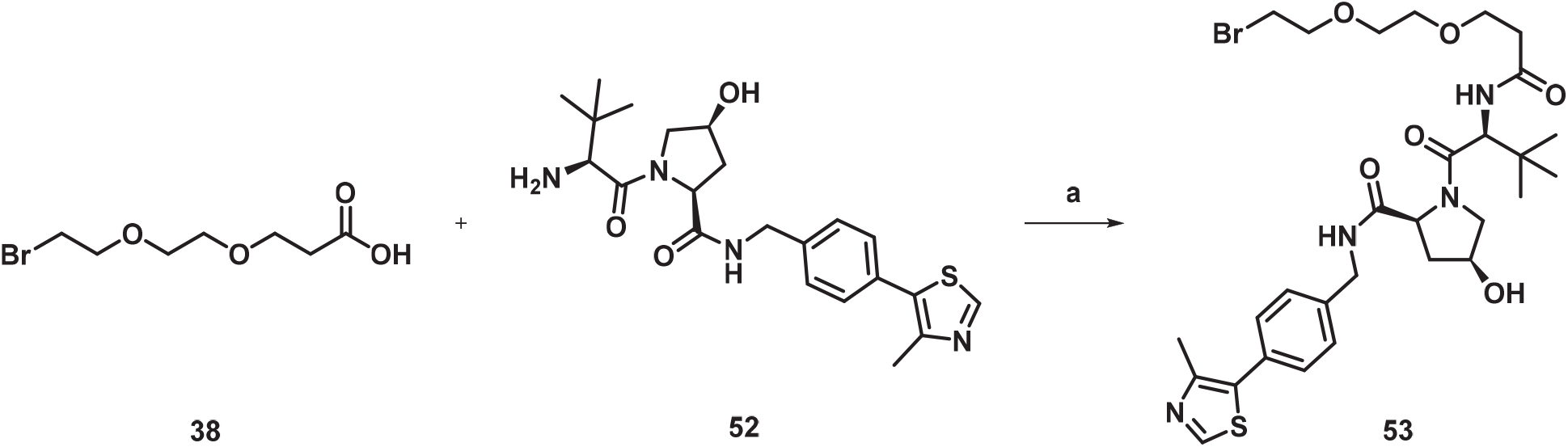

Scheme 19. Synthesis of Compound **53**^a^

^a^Reagents and conditions: (a) HATU, DIPEA, DMF, rt, 16h, 61%.

The synthetic route for preparing compound **YZ-6 NC** is outlined in Scheme 20. The hydroxyl of compound **12** was subjected to an S_N_2 reaction with compound **53** to give corresponding ether compound **YZ-6 NC**.

### Scheme 20

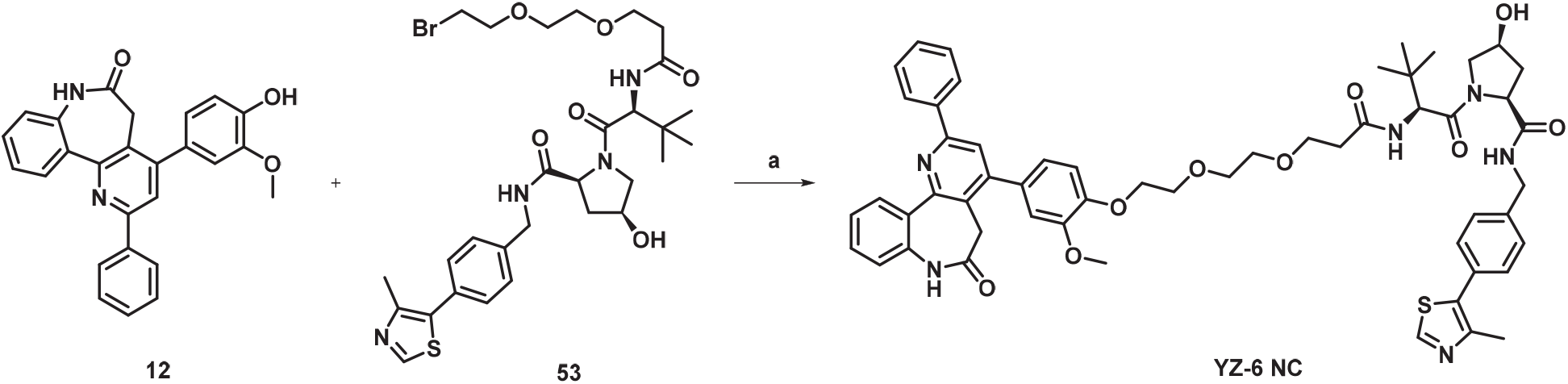

Scheme 20. Synthesis of Compound **YZ-6 NC**^a^

^a^Reagents and conditions: (a) K_2_CO_3_, KI, DMF, rt, 24h, 64%.

The synthetic route for preparing compounds **YZ-17, YZ-18** and **YZ-19** is outlined in Scheme 21. The hydroxyl of compound **19** was subjected to an S_N_2 reaction with compounds **42**, **43** and **44** to give corresponding ether compounds **YZ-17**, **YZ-18** and **YZ- 19**.

### Scheme 21

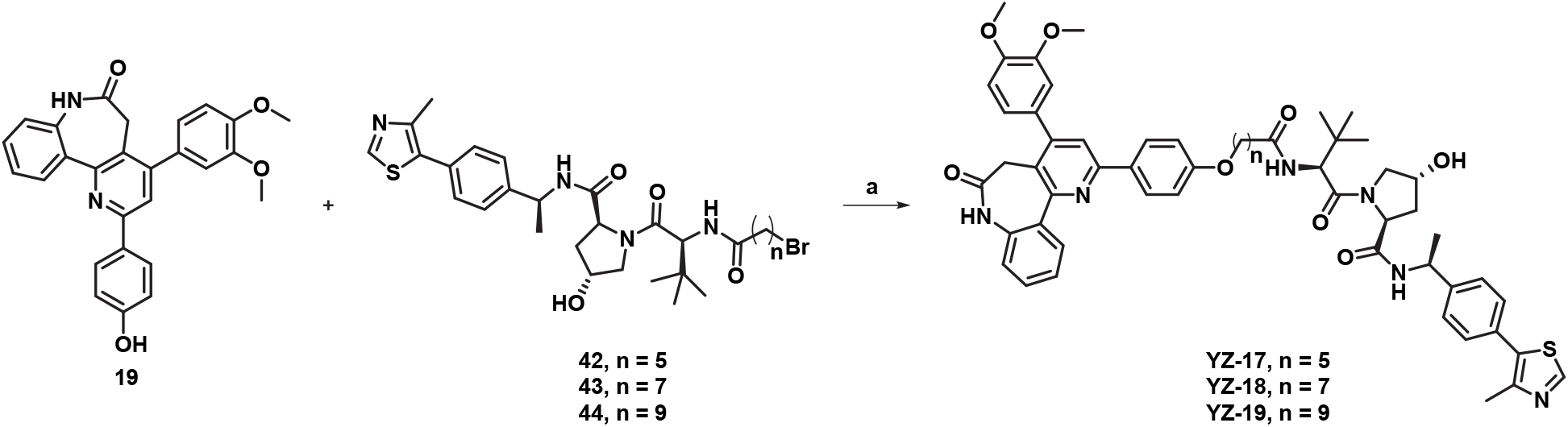

Scheme 21. Synthesis of Compounds **YZ-17**, **YZ-18** and **YZ-19**^a^

^a^Reagents and conditions: (a) K_2_CO_3_, KI, DMF, rt, 12h, 38% - 51%.

The synthetic route for preparing compound **YZ-20** is outlined in Scheme 22. The hydroxyl of compound **19** was subjected to an S_N_2 reaction with compound **40** to give corresponding ether compounds **YZ-20**.

### Scheme 22

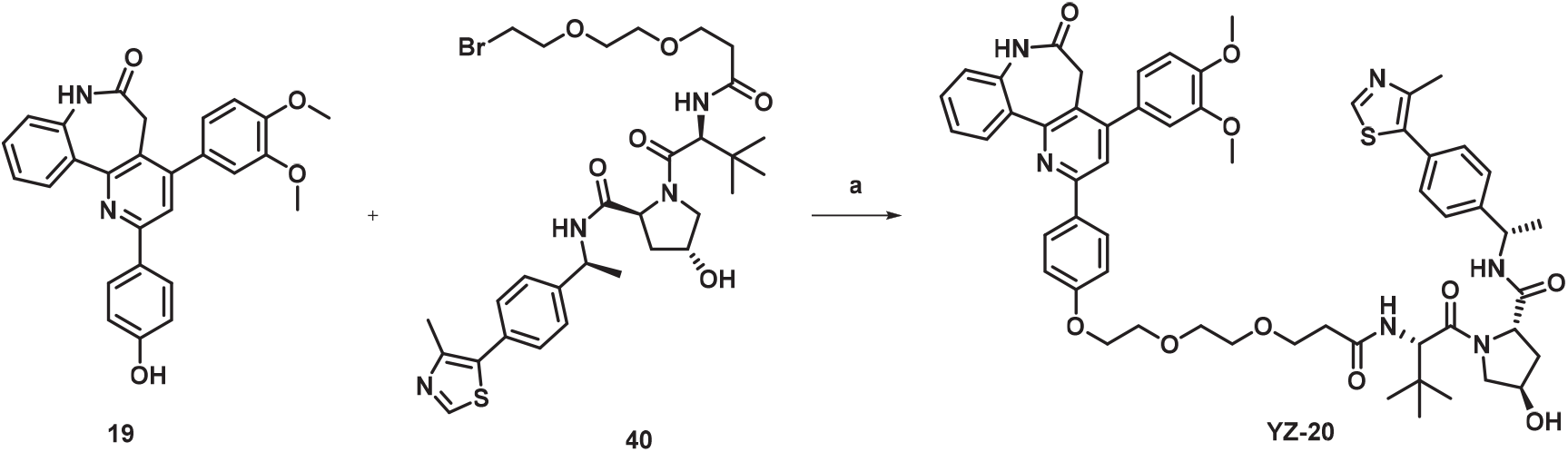

Scheme 22. Synthesis of Compound **YZ-20**^a^

^a^Reagents and conditions: (a) K_2_CO_3_, KI, DMF, rt, 24h, 51%.

The synthetic route for preparing compound **55** is outlined in Scheme 23. The hydroxyl of compound **19** was subjected to an S_N_2 reaction with *tert*-butyl 2-bromoacetate to give corresponding ether compound **54** (crude for next step). The *t*-butyl group of **54** was cleaved via TFA to afford compound **55**.

### Scheme 23

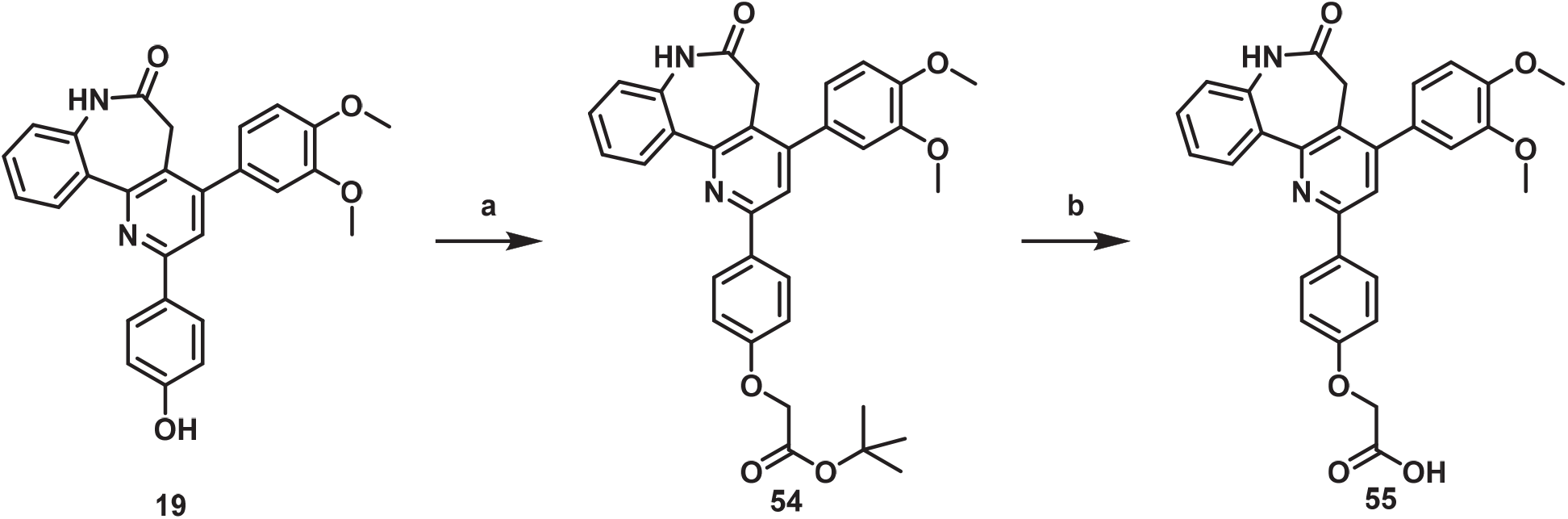

Scheme 23. Synthesis of Compound **55**^a^

^a^Reagents and conditions: (a) K_2_CO_3_, DMF, rt, 12h. (b) TFA, DCM, rt, 2h, 60% (2 steps).

The synthetic route for preparing compounds **YZ-21**, **YZ-22** and **YZ-23** is outlined in Scheme 24. Compound **55** was coupled with commercially available compounds **49**, **50** and **51** in the presence of HATU and DIPEA provided the corresponding amide compounds **YZ-21**, **YZ-22** and **YZ-23**.

### Scheme 24

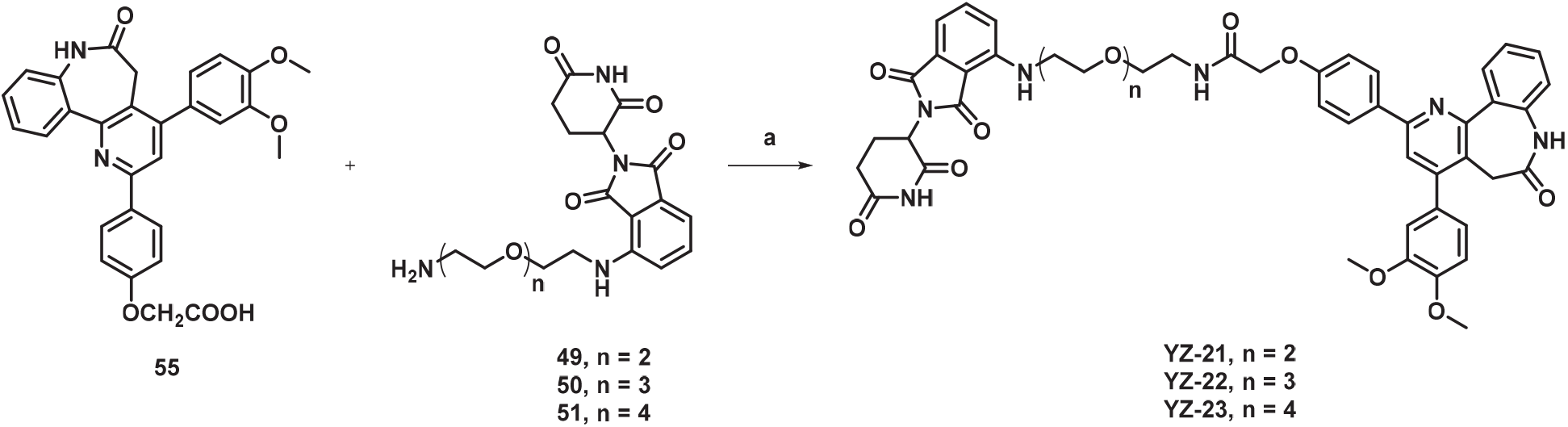

Scheme 24. Synthesis of Compounds **YZ-21**, **YZ-22** and **YZ-23**^a^

^a^Reagents and conditions: (a) HATU, DIPEA, DMF, rt, 16h, 45%-57%.

The synthetic route for preparing compound **56** is outlined in Scheme 25. Compound **31** was reacted with acetic anhydride in the presence of NEt_3_ provided the compound **56**.

### Scheme 25

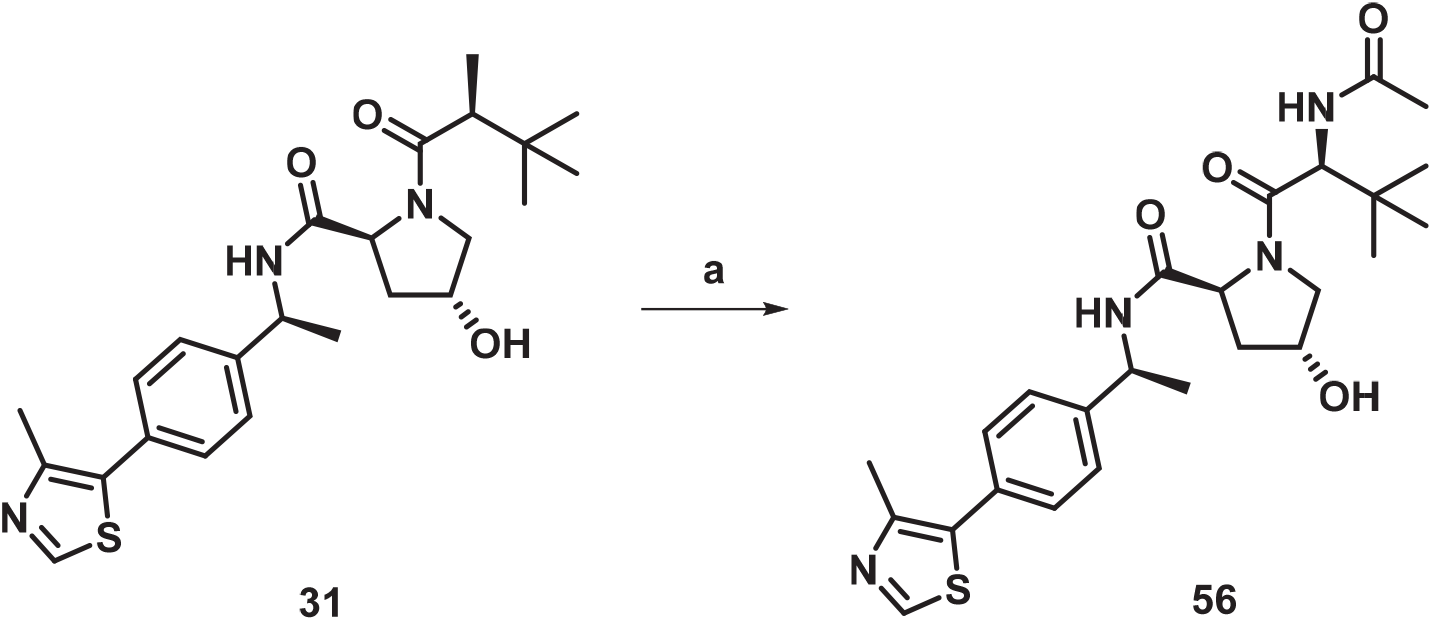

Scheme 25. Synthesis of Compound **56**^a^

^a^Reagents and conditions: (a) acetic anhydride, NEt_3_, DCM, rt, 2h, 78%

## Conclusion

This study represents the second report of PROTAC-induced YAP degradation in cancer cells. Our lead PROTAC molecule, **YZ- 6**, combines the YAP inhibitor NSC682769 with the VHL ligand 2. By recruiting VHL to YAP, **YZ-6** promotes YAP ubiquitination and subsequent degradation in YAP-dependent NCI-H226 and Huh7 cells with DC_50_ values of 4.3 and 8.2 μM, respectively. **YZ-6** is potent in inhibition of liver cancer cell Huh7 growth and is more than 20 times more potent than the YAP inhibitor NSC682769. This study demonstrates that **YZ-6** exhibits rapid engagement with YAP and sustains YAP degradation over time. Additionally, it leads to attenuated Hippo signaling in both NCI-H226 and Huh7 cell lines. More importantly, **YZ-6** exhibites potent anticancer efficacy in the Huh7 xenograft mouse model with a TGI rate of 82%. This tool compound, **YZ-6**, will serve as a valuable resource for further investigations into the impact of YAP degradation on downstream signaling pathways and the viability of cancer cells that overexpress YAP. Importantly, the use of PROTACs allows for more precise temporal control over YAP degradation compared to traditional knockdown methods, enabling researchers to explore the dynamic effects of YAP modulation on cellular processes.

The ability to target YAP with small molecules was itself a milestone in drug discovery. It showed that YAP, an “undruggable” intrinsically disordered protein, could be directly inhibited by a small molecule. Similarly, the results presented here demonstrate that YAP can be degraded as long as a suitable ligand is identified. While ligand development for YAP continues, **YZ-6** can serve as a tool compound to investigate biology in the context of rapid YAP degradation. Despite its limitations such as poor permeability and oral bioavailability, the discovery of **YZ-6** opens new opportunities for targeting YAP in cancer therapy.

### Experimental Section

### Chemical Reagents and General Method

All commercially available starting materials and solvents are reagent grade and used without further purification. Purification by silica gel column chromatography was carried out using the Biotage Isolera One purification system with prepacked cartridges. Analytical Thin Layer Chromatography was performed on silica gel plates and visualized under ultraviolet light (254 nm) for monitoring reactions. ^1^H and ^13^C NMR spectra were obtained on a Bruker Avance NEO-600 spectrometer or Bruker Avance III HD 600 (^1^H NMR: 600 MHz; ^13^C NMR: 151 MHz) using DMSO-*d*_6_, or CDCl_3_ as solvents.^59^ Chemical shifts (δ values) are expressed in ppm using tetramethyl silane as an internal standard (TMS ^1^H NMR: 0.00 ppm, ^13^C NMR: 0.00 ppm), and the coupling constants (*J* values) are indicated in hertz (Hz). High-resolution mass spectra were obtained from the Mass Spectrometry Facility within the University of Florida Department of Medicinal Chemistry Thermo Scientific™ Orbitrap Focus mass spectrometer with Dionex™ Ultimate™ 3000 UHPLC system.

### (*E*)-3-(3,4-dimethoxyphenyl)-1-phenylprop-2-en-1-one (3)

Acetophenone (600.8 mg, 5.0 mmol) was solubilized in ethanol (10.0 mL) and cooled in an ice bath for 15min and was added a solution of sodium hydroxide (600.0 mg, 15.0 mmol) in water (5.0 mL). Then 3,4-dimethoxybenzaldehyde (830.9 mg, 5.0 mmol) was added. The reaction mixture was remained under stirring during 18 h and monitored by TLC (thin layer chromatography) (ethyl acetate:hexane 1:5 v/v as eluent; UV lamp was used to visualize the plates). After, the reaction mixture was neutralized with HCl 5% until pH ≈ 7 and the solids formed were vacuum filtered and purified by recrystallization in ethanol to afford compound **3** (1113.5 mg, 83%) as a yellow solid. ^1^H NMR (600 MHz, Chloroform-*d*) δ 8.04 – 7.99 (m, 2H), 7.77 (d, *J* = 15.6 Hz, 1H), 7.61 – 7.55 (m, 1H), 7.54 – 7.48 (m, 2H), 7.39 (d, *J* = 15.6 Hz, 1H), 7.24 (dd, *J* = 8.3, 2.0 Hz, 1H), 7.17 (d, *J* = 2.0 Hz, 1H), 6.91 (d, *J* = 8.3 Hz, 1H), 3.96 (s, 3H), 3.94 (s, 3H). ^13^C NMR (151 MHz, Chloroform-*d*) δ 190.7, 151.5, 149.3, 145.1, 138.5, 132.6, 128.6, 128.5, 127.9, 123.2, 120.1, 111.2, 110.1, 56.0, 56.0.

### (*E*)-3-(4-(benzyloxy)-3-methoxyphenyl)-1-phenylprop-2-en-1-one (9)

Compound **9** was synthesized in a similar manner as described for compound **3**. Yellow solid, 81% yield. ^1^H NMR (600 MHz, Chloroform-*d*) δ 8.03 – 7.98 (m, 2H), 7.74 (d, *J* = 15.6 Hz, 1H), 7.60 – 7.55 (m, 1H), 7.53 – 7.47 (m, 2H), 7.46 – 7.41 (m, 2H), 7.41 – 7.35 (m, 3H), 7.35 – 7.29 (m, 1H), 7.20 – 7.14 (m, 2H), 6.91 (d, *J* = 8.1 Hz, 1H), 5.21 (s, 2H), 3.96 (s, 3H). ^13^C NMR (151 MHz, Chloroform-*d*) δ 190.7, 150.6, 149.8, 145.0, 138.5, 136.5, 132.6, 128.7, 128.6, 128.5, 128.2, 128.1, 127.3, 122.9, 120.2, 113.5, 110.8, 70.9, 56.1.

### (*E*)-1-(4-(benzyloxy)phenyl)-3-(3,4-dimethoxyphenyl)prop-2-en-1-one (16)

Compound **16** was synthesized in a similar manner as described for compound **3**. Yellow solid, 87% yield. ^1^H NMR (600 MHz, Chloroform-*d*) δ 8.06 – 8.01 (m, 2H), 7.75 (d, *J* = 15.5 Hz, 1H), 7.46 – 7.37 (m, 5H), 7.37 – 7.32 (m, 1H), 7.23 (dd, *J* = 8.4, 2.0 Hz, 1H), 7.16 (d, *J* = 2.0 Hz, 1H), 7.08 – 7.03 (m, 2H), 6.89 (d, *J* = 8.3 Hz, 1H), 5.15 (s, 2H), 3.95 (s, 3H), 3.93 (s, 3H). ^13^C NMR (151 MHz, Chloroform-*d*) δ 188.8, 162.5, 151.3, 149.3, 144.2, 136.3, 131.6, 130.8, 128.7, 128.3, 128.1, 127.5, 123.0, 119.8, 114.7, 111.2, 110.1, 77.3, 77.1, 76.9, 70.2, 56.0, 56.0.

### (*E*)-3-(3-(benzyloxy)-4-methoxyphenyl)-1-phenylprop-2-en-1-one (23)

Compound **23** was synthesized in a similar manner as described for compound **3**. Yellow solid, 78% yield. ^1^H NMR (600 MHz, DMSO-*d*_6_) δ 8.17 – 8.13 (m, 2H), 7.82 (d, *J* = 15.5 Hz, 1H), 7.73 – 7.64 (m, 3H), 7.59 (dd, *J* = 8.3, 7.1 Hz, 2H), 7.55 – 7.48 (m, 2H), 7.45 – 7.40 (m, 3H), 7.38 – 7.34 (m, 1H), 7.06 (d, *J* = 8.3 Hz, 1H), 5.19 (s, 2H), 3.83 (s, 3H). ^13^C NMR (151 MHz, DMSO-*d*_6_) δ 188.9, 151.5, 147.9, 144.4, 137.7, 136.8, 132.9, 128.6, 128.4, 128.3, 128.0, 127.9, 127.3, 124.2, 119.5, 112.2, 111.8, 70.0, 55.6.

### 4-(1-(3,4-dimethoxyphenyl)-3-oxo-3-phenylpropyl)-3,4-dihydro-1*H*-benzo[*b*]azepine-2,5-dione (5)

To a solution of **3** (512.9 mg, 1.9 mmol) and 3,4-dihydro-1H-benzo[b]azepine-2,5-dione (334.9 mg, 1.9 mmol) in 15 mL of ethanol was added 1.91 ml 0.1 M KOH. The resultant reaction mixture was stirred at 25 °C for 12 h. Upon completion of the reaction, the reaction mixture was neutralized with HCl 5% until pH ≈ 6 and the solids formed were vacuum filtered and purified by silica gel column chromatography (EtOAc:hexane = 1:1) to obtain compound **5** (640.4 mg, 76%) as a white solid. ^1^H NMR (600 MHz, DMSO-*d*_6_) δ 10.19 (s, 1H), 7.99 – 7.94 (m, 2H), 7.73 (dd, *J* = 7.8, 1.7 Hz, 1H), 7.63 – 7.58 (m, 1H), 7.50 (td, *J* = 7.7, 7.3, 3.1 Hz, 3H), 7.17 – 7.12 (m, 2H), 6.81 (d, *J* = 2.1 Hz, 1H), 6.76 (d, *J* = 8.3 Hz, 1H), 6.71 (dd, *J* = 8.3, 2.1 Hz, 1H), 4.06 (dt, *J* = 9.9, 4.7 Hz, 1H), 3.77 – 3.72 (m, 1H), 3.69 (d, *J* = 13.3 Hz, 6H), 3.46 (qd, *J* = 7.0, 5.0 Hz, 1H), 3.26 – 3.16 (m, 2H), 2.93 (dd, *J* = 14.6, 11.2 Hz, 1H), 2.66 (dt, *J* = 14.6, 1.7 Hz, 1H). ^13^C NMR (151 MHz, DMSO-*d*_6_) δ 200.5, 198.5, 172.9, 148.8, 147.8, 138.9, 137.2, 134.2, 134.2, 133.5, 131.0, 129.0, 128.4, 127.9, 123.8, 121.7, 120.1, 112.6, 111.8, 56.6, 55.8, 55.8, 53.5, 41.3, 38.9.

### 4-(1-(4-(benzyloxy)-3-methoxyphenyl)-3-oxo-3-phenylpropyl)-3,4-dihydro-1*H*-benzo[*b*]azepine-2,5-dione (10)

Compound **10** was synthesized in a similar manner as described for compound **5**. White solid, 72% yield. ^1^H NMR (600 MHz, DMSO-*d*_6_) δ 10.21 (s, 1H), 7.99 – 7.96 (m, 2H), 7.74 (d, *J* = 8.4 Hz, 1H), 7.63 (t, *J* = 7.4 Hz, 1H), 7.55 – 7.49 (m, 3H), 7.42 – 7.37 (m, 4H), 7.32 (t, *J* = 7.2 Hz, 1H), 7.16 (t, *J* = 7.6 Hz, 2H), 6.89 – 6.84 (m, 2H), 6.69 (dd, *J* = 8.3, 1.8 Hz, 1H), 4.99 (s, 2H), 4.06 (dt, *J* = 9.9, 4.6 Hz, 1H), 3.78 (dd, *J* = 17.4, 10.5 Hz, 1H), 3.67 (s, 3H), 3.30 (br s, 2H), 3.25 – 3.13 (m, 3H), 2.95 (dd, *J* = 14.7, 11.3 Hz, 1H), 2.68 – 2.60 (m, 1H).

### 4-(3,4-dimethoxyphenyl)-2-phenyl-5,7-dihydro-6*H*-benzo[*b*]pyrido[2,3-*d*]azepin-6-one (6, NSC682769)

To accomplish the ring closure reaction, a mixture of the Michael adduct **5** (309.9 mg, 0.70 mmol), ammonium ferric sulfate dodecahydrate (697 mg, 1.45 mmol), and ammonium acetate (844 mg, 10.95 mmol) was refluxed in glacial acetic acid (10.0 mL) under nitrogen for 5 h. After cooling, the mixture was poured on crushed ice (30 g) and stirred until melting of the ice. The precipitate was collected by filtration, washed with water, and recrystallized from ethanol/toluene to afford compound **6** (251.4 mg, 85%) as a white solid. ^1^H NMR (600 MHz, DMSO-*d*_6_) δ 10.41 (s, 1H), 8.23 (ddd, *J* = 11.0, 8.3, 1.6 Hz, 3H), 7.97 (s, 1H), 7.56 – 7.50 (m, 3H), 7.50 – 7.44 (m, 1H), 7.37 (td, *J* = 7.5, 1.2 Hz, 1H), 7.29 (d, *J* = 2.1 Hz, 1H), 7.25 (ddd, *J* = 16.7, 8.2, 1.7 Hz, 2H), 7.16 (d, *J* = 8.3 Hz, 1H), 3.91 (s, 3H), 3.86 (s, 3H), 3.73 (s, 1H), 3,45 (br s, 2H). ^13^C NMR (151 MHz, DMSO-*d*_6_) δ 172.0, 155.0, 154.7, 149.8, 149.6, 149.0, 138.9, 138.0, 131.6, 131.6, 130.6, 130.4, 129.6, 129.2, 127.3, 126.0, 124.4, 122.6, 122.0, 120.8, 114.1, 112.3, 56.1, 56.1, 36.0.

### 4-(4-(benzyloxy)-3-methoxyphenyl)-2-phenyl-5,7-dihydro-6*H*-benzo[*b*]pyrido[2,3-*d*]azepin-6-one (11)

Compound **11** was synthesized in a similar manner as described for compound **6**. White solid, 82% yield. ^1^H NMR (600 MHz, DMSO-*d*_6_) δ 10.41 (s, 1H), 8.24 (ddd, *J* = 12.4, 8.4, 1.6 Hz, 3H), 7.97 (s, 1H), 7.56 – 7.50 (m, 5H), 7.50 – 7.46 (m, 1H), 7.44 (t, *J* = 7.6 Hz, 2H), 7.40 – 7.34 (m, 2H), 7.32 (d, *J* = 2.0 Hz, 1H), 7.29 – 7.23 (m, 2H), 7.21 (dd, *J* = 8.2, 2.0 Hz, 1H), 5.21 (s, 2H), 3.94 (s, 3H), 3,45 (br s, 2H).

### 4-(3-(benzyloxy)-4-methoxyphenyl)-2-phenyl-5,7-dihydro-6*H*-benzo[*b*]pyrido[2,3-*d*]azepin-6-one (25)

Compound **25** was synthesized using crude compound **24** in a similar manner as described for compound **6**. White solid, 56% yield (2 steps). ^1^H NMR (600 MHz, DMSO-*d*_6_) δ 10.45 (s, 1H), 8.22 (dq, *J* = 8.3, 1.7 Hz, 3H), 7.91 (s, 1H), 7.57 – 7.51 (m, 3H), 7.51 – 7.44 (m, 3H), 7.43 – 7.31 (m, 5H), 7.26 (td, *J* = 8.2, 1.6 Hz, 2H), 7.20 (d, *J* = 8.3 Hz, 1H), 3.88 (s, 3H), 3.40 (br s, 2H). ^13^C NMR (151 MHz, DMSO-*d*_6_) δ 171.5, 154.4, 154.1, 149.3, 149.0, 147.4, 138.3, 137.4, 137.0, 131.0, 131.0, 129.9, 129.8, 129.1, 128.7, 128.3, 127.8, 127.7, 126.7, 125.4, 123.8, 122.4, 121.4, 120.2, 115.1, 112.0, 69.8, 55.6, 35.4.

### 4-(3,4-dimethoxyphenyl)-7-ethyl-2-phenyl-5,7-dihydro-6*H*-benzo[*b*]pyrido[2,3-*d*]azepin-6-one (7)

Sodium hydride (2.17 mg, 0.09 mmol) was added to a solution of **6** (25.5 mg, 0.06 mmol) in THF (2 mL). After the mixture was stirred for 5 min, a solution of bromomethane (13.2 mg, 0.12 mmol) and KI (20 mg, 0.12 mmol) in THF (3 mL) was added and was stirred for 24 h. Upon completion of the reaction, the reaction mixture was quenched with water and extracted with ethyl acetate three times. The combined organic layers were washed with brine, dried over anhydrous Na_2_SO_4,_ and concentrated under reduced pressure. Purify the residue further by silica gel column chromatography (EtOAc:hexane = 1:2) to obtain compound 7 (14.9 mg, 55%) as a white solid. ^1^H NMR (600 MHz, Chloroform-*d*) δ 8.22 (dd, *J* = 7.8, 1.7 Hz, 1H), 8.17 – 8.12 (m, 2H), 7.79 (s, 1H), 7.56 – 7.47 (m, 3H), 7.47 – 7.38 (m, 4H), 7.19 (d, *J* = 8.2 Hz, 1H), 7.04 (d, *J* = 8.3 Hz, 1H), 4.15 (ddt, *J* = 27.7, 14.2, 7.1 Hz, 2H), 4.07 (s, 3H), 3.97 (s, 3H), 3.94 (d, *J* = 12.8 Hz, 1H), 3.80 (dq, *J* = 14.1, 7.1 Hz, 1H), 3.16 (d, *J* = 12.8 Hz, 1H), 1.15 (t, *J* = 7.1 Hz, 3H). ^13^C NMR (151 MHz, Chloroform-*d*) δ 170.3, 155.8, 154.4, 149.4, 149.2, 148.8, 141.1, 139.1, 134.7, 131.6, 130.9, 129.7, 129.1, 128.8, 127.2, 127.1, 125.5, 122.4, 121.1, 113.5, 111.2, 99.8, 56.4, 56.0, 43.5, 35.9, 13.7.

### 4-(4-hydroxy-3-methoxyphenyl)-2-phenyl-5,7-dihydro-6*H*-benzo[*b*]pyrido[2,3-*d*]azepin-6-one (12)

Stir a mixture of **11** (99.7 mg, 0.20 mmol) and Pd/C (10 mg, 10 wt. % Pd) in methanol (5 mL) in a reaction vial equipped with a H_2_ balloon at 60 °C for 12 h. Monitor the reaction progress by thin layer chromatography (TLC). Filter the mixture upon completion. Remove the methanol under vacuum. Purify the residue further by silica gel column chromatography (EtOAc:hexane = 1:1) to obtain compound 12 (73.5 mg, 76.1%) as a white solid. ^1^H NMR (600 MHz, DMSO-*d*_6_) δ 10.40 (s, 1H), 9.36 (s, 1H), 8.23 (ddd, *J* = 9.6, 8.2, 1.5 Hz, 3H), 7.94 (s, 1H), 7.56 – 7.50 (m, 3H), 7.50 – 7.44 (m, 1H), 7.37 (td, *J* = 7.5, 1.2 Hz, 1H), 7.30 – 7.23 (m, 2H), 7.11 (dd, *J* = 8.0, 2.1 Hz, 1H), 6.97 (d, *J* = 8.1 Hz, 1H), 3.93 (s, 3H), 3,45 (br s, 2H). ^13^C NMR (151 MHz, DMSO-*d*_6_) δ 172.1, 154.9, 154.6, 150.1, 147.9, 147.5, 139.0, 138.0, 131.6, 130.3, 129.6, 129.2, 129.1, 127.3, 126.0, 124.4, 122.9, 122.0, 121.6, 120.7, 116.0, 114.6, 56.2, 35.9.

### 4-(3,4-dimethoxyphenyl)-2-(4-hydroxyphenyl)-5,7-dihydro-6*H*-benzo[*b*]pyrido[2,3-*d*]azepin-6-one (19)

Compound **19** was synthesized from crude compound **18** in a similar manner as described for compound **12**. White solid, 58% yield (3 steps). ^1^H NMR (600 MHz, DMSO-*d*_6_) δ 10.37 (s, 1H), 9.76 (s, 1H), 8.20 (dd, *J* = 7.8, 1.6 Hz, 1H), 8.12 – 8.06 (m, 2H), 7.84 (s, 1H), 7.53 (ddd, *J* = 8.2, 7.3, 1.7 Hz, 1H), 7.36 (ddd, *J* = 8.1, 7.3, 1.3 Hz, 1H), 7.29 – 7.23 (m, 2H), 7.21 (dd, *J* = 8.2, 2.1 Hz, 1H), 7.16 (d, *J* = 8.3 Hz, 1H), 6.92 – 6.87 (m, 2H), 3.91 (s, 3H), 3.86 (s, 3H), 3.45 (br s, 2H).

### 4-(3-hydroxy-4-methoxyphenyl)-2-phenyl-5,7-dihydro-6*H*-benzo[*b*]pyrido[2,3-*d*]azepin-6-one (26)

Compound **26** was synthesized in a similar manner as described for compound **12**. White solid, 83% yield. ^1^H NMR (600 MHz, DMSO-*d*_6_) δ 10.37 (s, 1H), 9.23 (s, 1H), 8.25 – 8.17 (m, 3H), 7.89 (s, 1H), 7.52 (ddd, *J* = 9.9, 5.4, 2.9 Hz, 3H), 7.49 – 7.43 (m, 1H), 7.36 (ddd, *J* = 8.2, 7.3, 1.2 Hz, 1H), 7.24 (dd, *J* = 8.1, 1.2 Hz, 1H), 7.13 (d, *J* = 8.1 Hz, 1H), 7.10 – 7.04 (m, 2H), 3.87 (s, 3H), 3.40 (br s, 2H).

### 2-(2-methoxy-4-(6-oxo-2-phenyl-6,7-dihydro-5*H*-benzo[*b*]pyrido[2,3-*d*]azepin-4-yl)phenoxy)ethyl acetate (13)

To a solution of compound **12** (61.2 mg, 0.15 mmol) in N, N-dimethylformamide (4 ml) was added 2-bromoethyl acetate (50.1 mg, 0.3 mmol), potassium carbonate (62.2 mg, 0.45 mmol). The resultant reaction mixture was stirred at 25 °C for 10 h. Upon completion of the reaction, the reaction mixture was quenched with 100 ml water and extracted with ethyl acetate three times. The combined organic layers were washed with brine, dried over anhydrous Na_2_SO_4_ and concentrated under reduced pressure. Purify the residue further by silica gel column chromatography (EtOAc:hexane = 1:1) to obtain compound **13** (48.2 mg, 65%) as a white solid. ^1^H NMR (600 MHz, DMSO-*d*_6_) δ 10.42 (s, 1H), 8.24 (ddd, *J* = 11.1, 8.3, 1.6 Hz, 3H), 7.97 (s, 1H), 7.57 – 7.51 (m, 3H), 7.51 – 7.45 (m, 1H), 7.38 (ddd, *J* = 8.2, 7.3, 1.3 Hz, 1H), 7.32 (d, *J* = 2.0 Hz, 1H), 7.26 (dd, *J* = 8.1, 1.3 Hz, 1H), 7.25 – 7.17 (m, 2H), 4.41 – 4.36 (m, 2H), 4.29 (dd, *J* = 5.6, 3.5 Hz, 2H), 3.93 (s, 3H), 3,45 (br s, 2H), 2.08 (s, 3H). ^13^C NMR (151 MHz, DMSO-*d*_6_) δ 172.0, 170.9, 155.0, 154.7, 149.7, 149.3, 148.4, 138.9, 137.9, 131.6, 131.6, 131.2, 130.4, 129.7, 129.2, 127.3, 126.0, 124.4, 122.5, 122.0, 120.7, 114.5, 114.0, 67.2, 63.1, 56.1, 36.0, 21.2.

### 2-(4-(4-(3,4-dimethoxyphenyl)-6-oxo-6,7-dihydro-5*H*-benzo[*b*]pyrido[2,3-*d*]azepin-2-yl)phenoxy)ethyl acetate (20)

Compound **20** was synthesized in a similar manner as described for compound **13**. White solid, 60% yield. ^1^H NMR (600 MHz, DMSO-*d*_6_) δ 10.39 (s, 1H), 8.25 – 8.18 (m, 3H), 7.91 (s, 1H), 7.53 (ddd, J = 8.3, 7.3, 1.6 Hz, 1H), 7.37 (td, J = 7.6, 1.3 Hz, 1H), 7.32 – 7.20 (m, 4H), 7.16 (d, J = 8.3 Hz, 1H), 7.13 – 7.07 (m, 2H), 4.37 (dd, J = 5.6, 3.4 Hz, 2H), 4.29 – 4.25 (m, 2H), 3.91 (s, 3H), 3.86 (s, 3H), 3,45 (br s, 2H), 2.07 (s, 3H). ^13^C NMR (151 MHz, DMSO-*d*_6_) δ 171.6, 170.4, 159.2, 154.1, 154.0, 149.2, 149.0, 148.5, 137.4, 131.2, 131.19, 131.18, 130.2, 129.8, 128.2, 124.8, 123.9, 122.0, 121.5, 119.5, 114.7, 113.6, 111.8, 65.9, 62.5, 55.6, 55.6, 35.4, 20.7.

### 2-(2-methoxy-5-(6-oxo-2-phenyl-6,7-dihydro-5*H*-benzo[*b*]pyrido[2,3-*d*]azepin-4-yl)phenoxy)ethyl acetate (27)

Compound **27** was synthesized in a similar manner as described for compound **13**. White solid, 61% yield. ^1^H NMR (600 MHz, DMSO-*d*_6_) δ 10.42 (s, 1H), 8.26 – 8.20 (m, 3H), 7.97 (s, 1H), 7.56 – 7.50 (m, 3H), 7.50 – 7.44 (m, 1H), 7.37 (ddd, *J* = 8.3, 7.2, 1.2 Hz, 1H), 7.32 (d, *J* = 2.1 Hz, 1H), 7.27 (ddd, *J* = 13.1, 8.2, 1.7 Hz, 2H), 7.19 (d, *J* = 8.2 Hz, 1H), 4.38 (m, 4H), 3.87 (s, 3H), 3.40 (br s, 2H), 2.05 (s, 3H). ^13^C NMR (151 MHz, DMSO-*d*_6_) δ 172.0, 170.8, 155.0, 154.7, 149.8, 149.6, 147.9, 138.9, 137.9, 131.6, 131.6, 130.6, 130.4, 129.7, 129.2, 127.3, 126.0, 124.4, 123.2, 122.0, 120.8, 115.6, 112.7, 67.1, 63.1, 56.2, 35.9, 21.2.

### 4-(4-(2-hydroxyethoxy)-3-methoxyphenyl)-2-phenyl-5,7-dihydro-6*H*-benzo[*b*]pyrido[2,3-*d*]azepin-6-one (14)

To a solution of compound **13** (29.7 mg, 0.06 mmol) in methanol (4 ml) was added potassium carbonate (41.5 mg, 0.3 mmol). The resultant reaction mixture was stirred at 25 °C for 6 h. Upon completion of the reaction, the reaction mixture was quenched with 10 ml water. Remove the methanol under vacuum and extract with ethyl acetate three times. The combined organic layers were washed with brine, dried over anhydrous Na_2_SO_4_ and concentrated under reduced pressure. Purify the residue further by silica gel column chromatography (EtOAc:hexane = 1:1) to obtain compound **14** (25.8 mg, 95%) as a white solid. ^1^H NMR (600 MHz, DMSO-*d*_6_) δ 10.41 (s, 1H), 8.24 (ddd, *J* = 11.4, 8.3, 1.6 Hz, 3H), 7.97 (s, 1H), 7.57 – 7.51 (m, 3H), 7.51 – 7.45 (m, 1H), 7.38 (ddd, *J* = 8.2, 7.4, 1.3 Hz, 1H), 7.30 (d, *J* = 2.0 Hz, 1H), 7.27 (dd, *J* = 8.1, 1.2 Hz, 1H), 7.22 (dd, *J* = 8.3, 2.1 Hz, 1H), 7.18 (d, *J* = 8.3 Hz, 1H), 4.92 (t, *J* = 5.5 Hz, 1H), 4.09 (t, *J* = 5.1 Hz, 2H), 3.92 (s, 3H), 3.78 (q, *J* = 5.3 Hz, 2H), 3,45 (br s, 2H). ^13^C NMR (151 MHz, DMSO-*d*_6_) δ 172.0, 155.0, 154.7, 149.8, 149.2, 148.9, 138.9, 137.9, 131.6, 130.7, 130.4, 129.7, 129.2, 127.3, 126.0, 124.4, 122.5, 122.0, 120.7, 114.3, 113.5, 70.8, 60.1, 56.0, 36.0.

### 4-(3,4-dimethoxyphenyl)-2-(4-(2-hydroxyethoxy)phenyl)-5,7-dihydro-6*H*-benzo[*b*]pyrido[2,3-*d*]azepin-6-one (21)

Compound **21** was synthesized in a similar manner as described for compound **14**. White solid, 96% yield. ^1^H NMR (600 MHz, DMSO-*d*_6_) δ ^1^H NMR (600 MHz, DMSO-*d*_6_) δ 10.38 (s, 1H), 8.19 (td, *J* = 7.2, 1.9 Hz, 3H), 7.89 (s, 1H), 7.52 (td, *J* = 7.6, 1.6 Hz, 1H), 7.36 (t, *J* = 7.5 Hz, 1H), 7.29 – 7.23 (m, 2H), 7.21 (dd, *J* = 8.3, 2.1 Hz, 1H), 7.15 (d, *J* = 8.3 Hz, 1H), 7.10 – 7.04 (m, 2H), 4.06 (t, *J* = 5.0 Hz, 2H), 3.85 (s, 3H), 3.75 (t, *J* = 5.0 Hz, 2H), 3.24 (br s, 2H).

### *tert*-butyl 3-(2-(tosyloxy)ethoxy)propanoate (29)

To a solution of compound **28** (190.2 mg, 1 mmol) in DCM (6 ml) was added p-Toluenesulfonyl chloride (381.3 mg, 2 mmol), NEt_3_ (303.6 mg, 3 mmol), DMAP (12.3 mg, 0.1 mmol). The resultant reaction mixture was stirred at 25 °C for 8 h. Upon completion of the reaction, the reaction mixture was quenched with 20 ml water and extracted with ethyl acetate three times. The combined organic layers were washed with brine, dried over anhydrous Na_2_SO_4_ and concentrated under reduced pressure. Purify the residue further by silica gel column chromatography (EtOAc:hexane = 1:3) to obtain compound **29** (282.4 mg, 82%) as a white solid. ^1^H NMR (600 MHz, Chloroform-*d*) δ 7.82 – 7.77 (m, 2H), 7.37 – 7.32 (m, 2H), 4.17 – 4.12 (m, 2H), 3.66 – 3.61 (m, 4H), 2.45 (s, 3H), 2.42 (t, *J* = 6.4 Hz, 2H), 1.44 (s, 9H).

### 3-(2-(tosyloxy)ethoxy)propanoic acid (30)

To a solution of compound **29** (275.5 mg, 0.8 mmol) in DCM (3 ml) was added 1 ml TFA. The resultant reaction mixture was stirred at 25 °C for 2 h. Remove the DCM and TFA under vacuum. The material (219.1 mg, 95%) is used for subsequent reactions without further purification. ^1^H NMR (600 MHz, Chloroform-*d*) δ 7.82 – 7.77 (m, 2H), 7.37 – 7.32 (m, 2H), 4.18 – 4.13 (m, 2H), 3.73 – 3.64 (m, 4H), 2.58 (t, *J* = 6.2 Hz, 2H), 2.45 (s, 3H).

### 2-(3-(((*S*)-1-((2*S*,4*R*)-4-hydroxy-2-(((*S*)-1-(4-(4-methylthiazol-5-yl)phenyl)ethyl)carbamoyl)pyrrolidin-1-yl)-3,3-dimethyl-1-oxobutan-2-yl)amino)-3-oxopropoxy)ethyl 4-methylbenzenesulfonate (32)

Compound **30** (200 mg, 0.416 mmol,), **31** (210 mg, 0.437 mmol), HATU (206 mg, 0.54 mmol), and diisopropylethylamine (268.8 mg, 2.08 mmol) were dissolved in N, N-dimethylformamide (6 ml) and stirred at 25 °C for 8 h. After the reaction was complete (monitored by TLC), water (150 mL) was added. The organic layer was extracted with ethyl acetate and washed with brine, dried over Na_2_SO_4_, filtered and then concentrated. The residue was purified further by silica gel column chromatography (DCM:methanol = 15:1) to obtain compound **32** (220 mg, 74%) as a white solid. ^1^H NMR (600 MHz, Chloroform-*d*) δ 8.67 (s, 1H), 7.83 – 7.78 (m, 2H), 7.49 (d, *J* = 7.9 Hz, 1H), 7.42 – 7.32 (m, 6H), 6.77 (d, *J* = 8.2 Hz, 1H), 5.08 (p, *J* = 7.1 Hz, 1H), 4.75 (t, *J* = 7.9 Hz, 1H), 4.51 (d, *J* = 8.2 Hz, 2H), 4.17 (td, *J* = 4.2, 1.5 Hz, 2H), 4.14 – 4.10 (m, 1H), 3.71 – 3.64 (m, 4H), 3.58 (dd, *J* = 11.4, 3.7 Hz, 1H), 3.09 (d, *J* = 4.2 Hz, 1H), 2.58 – 2.53 (m, 1H), 2.52 (s, 3H), 2.44 (d, *J* = 6.5 Hz, 5H), 2.10 – 2.04 (m, 2H), 1.47 (d, *J* = 6.9 Hz, 3H), 1.25 (dt, *J* = 10.3, 7.1 Hz, 1H), 1.04 (s, 9H).

### (2*S*,4*R*)-1-((*S*)-2-(2-(2-bromoethoxy)acetamido)-3,3-dimethylbutanoyl)-4-hydroxy-*N*-((*S*)-1-(4-(4-methylthiazol-5-yl)phenyl)ethyl)pyrrolidine-2-carboxamide (35)

Compound **35** was synthesized in a similar manner as described for compound **32**. White solid, 70% yield. ^1^H NMR (600 MHz, Chloroform-*d*) δ 8.67 (s, 1H), 7.48 (d, *J* = 7.8 Hz, 1H), 7.43 – 7.38 (m, 2H), 7.38 – 7.35 (m, 2H), 7.30 (d, *J* = 8.2 Hz, 1H), 5.08 (p, *J* = 7.1 Hz, 1H), 4.77 (t, *J* = 7.8 Hz, 1H), 4.54 – 4.48 (m, 2H), 4.18 – 4.10 (m, 1H), 4.04 (d, *J* = 2.9 Hz, 2H), 3.90 (dt, *J* = 10.9, 5.4 Hz, 1H), 3.83 (dt, *J* = 10.8, 5.8 Hz, 1H), 3.60 (dd, *J* = 11.4, 3.7 Hz, 1H), 3.53 (t, *J* = 5.6 Hz, 2H), 2.89 (s, 1H), 2.59 (ddd, *J* = 13.5, 7.4, 4.7 Hz, 1H), 2.53 (s, 3H), 2.10 – 2.03 (m, 2H), 1.48 (d, *J* = 7.0 Hz, 3H), 1.09 (s, 9H).

### (2*S*,4*R*)-1-((*S*)-2-(3-(2-(2-bromoethoxy)ethoxy)propanamido)-3,3-dimethylbutanoyl)-4-hydroxy-*N*-((*S*)-1-(4-(4-methylthiazol-5-yl)phenyl)ethyl)pyrrolidine-2-carboxamide (40)

Compound **40** was synthesized in a similar manner as described for compound **32**. White solid, 55% yield. ^1^H NMR (600 MHz, Chloroform-*d*) δ 8.67 (s, 1H), 7.51 (d, *J* = 7.8 Hz, 1H), 7.43 – 7.39 (m, 2H), 7.37 (d, *J* = 8.3 Hz, 2H), 7.02 (d, *J* = 8.1 Hz, 1H), 5.08 (p, *J* = 7.1 Hz, 1H), 4.76 (t, *J* = 7.9 Hz, 1H), 4.49 (t, *J* = 7.1 Hz, 2H), 4.17 (dt, *J* = 11.5, 1.9 Hz, 1H), 3.80 (t, *J* = 6.3 Hz, 2H), 3.77 – 3.72 (m, 2H), 3.72 – 3.61 (m, 4H), 3.57 (dd, *J* = 11.5, 3.6 Hz, 1H), 3.48 (t, *J* = 6.3 Hz, 2H), 3.10 (s, 1H), 2.58 (ddd, *J* = 13.6, 7.6, 4.7 Hz, 1H), 2.53 (s, 3H), 2.53 – 2.48 (m, 2H), 2.06 (ddt, *J* = 13.6, 8.3, 2.0 Hz, 1H), 1.47 (d, *J* = 6.9 Hz, 3H), 1.06 (s, 9H).

### (2*S*,4*R*)-1-((*S*)-1-bromo-14-(*tert*-butyl)-12-oxo-3,6,9-trioxa-13-azapentadecan-15-oyl)-4-hydroxy-*N*-((*S*)-1-(4-(4-methylthiazol-5-yl)phenyl)ethyl)pyrrolidine-2-carboxamide (41)

Compound **41** was synthesized in a similar manner as described for compound **32**. White solid, 62% yield. ^1^H NMR (600 MHz, DMSO-*d*_6_) δ 8.98 (s, 1H), 8.37 (d, *J* = 7.8 Hz, 1H), 7.88 – 7.84 (m, 1H), 7.43 (d, *J* = 8.2 Hz, 2H), 7.38 (d, *J* = 8.2 Hz, 2H), 5.10 (d, *J* = 3.6 Hz, 1H), 4.92 (t, *J* = 7.3 Hz, 1H), 4.53 (d, *J* = 9.4 Hz, 1H), 4.42 (t, *J* = 8.0 Hz, 1H), 4.28 (s, 1H), 3.73 (t, *J* = 5.8 Hz, 2H), 3.63 – 3.55 (m, 8H), 3.53 – 3.46 (m, 6H), 2.56 – 2.52 (m, 1H), 2.45 (s, 3H), 2.40 – 2.32 (m, 1H), 2.04 – 1.98 (m, 1H), 1.79 (ddd, *J* = 12.9, 8.5, 4.7 Hz, 1H), 1.38 (d, *J* = 7.0 Hz, 3H), 0.93 (s, 9H).

### (2S,4R)-4-hydroxy-1-((S)-2-(3-(2-(2-(2-methoxy-4-(6-oxo-2-phenyl-6,7-dihydro-5H-benzo[b]pyrido[2,3-d]azepin-4-yl)phenoxy)ethoxy)ethoxy)propanamido)-3,3-dimethylbutanoyl)-N-((S)-1-(4-(4-methylthiazol-5- yl)phenyl)ethyl)pyrrolidine-2-carboxamide (YZ-6)

To a solution of compound **12** (91.8 mg, 0.225 mmol) and **40** (150 mg, 0.225 mmol) in DMF (6 ml) was added potassium carbonate (93.1 mg, 0.674 mmol), potassium iodide (372.9 mg, 2.25 mmol). The resultant reaction mixture was stirred at 25 °C for 12 h. Upon completion of the reaction, the reaction mixture was quenched with 200 ml water and extracted with ethyl acetate three times. The combined organic layers were washed with brine, dried over anhydrous Na_2_SO_4_ and concentrated under reduced pressure. Purify the residue further by silica gel column chromatography (DCM:methanol = 10:1) to obtain compound **YZ-6** (118.7 mg, 53%) as a white solid. ^1^H NMR (600 MHz, DMSO-*d*_6_) δ 10.41 (s, 1H), 8.98 (s, 1H), 8.37 (d, J = 7.8 Hz, 1H), 8.26 – 8.20 (m, 3H), 7.96 (s, 1H), 7.88 (d, J = 9.3 Hz, 1H), 7.56 – 7.50 (m, 3H), 7.50 – 7.45 (m, 1H), 7.45 – 7.40 (m, 2H), 7.40 – 7.32 (m, 3H), 7.30 (d, J = 2.1 Hz, 1H), 7.26 (dd, J = 8.1, 1.2 Hz, 1H), 7.22 (dd, J = 8.2, 2.0 Hz, 1H), 7.18 (d, J = 8.2 Hz, 1H), 5.11 (d, J = 3.6 Hz, 1H), 4.91 (p, J = 7.3 Hz, 1H), 4.54 (d, J = 9.3 Hz, 1H), 4.43 (t, J = 8.1 Hz, 1H), 4.28 (s, 1H), 4.21 – 4.16 (m, 2H), 3.92 (s, 3H), 3.82 – 3.76 (m, 2H), 3.66 – 3.59 (m, 6H), 3.55 (dt, J = 10.4, 5.0 Hz, 2H), 3.31 (br s, 2H), 2.56 (dt, J = 14.0, 6.8 Hz, 1H), 2.45 (s, 3H), 2.38 (dt, J = 14.6, 6.2 Hz, 1H), 2.05 – 1.98 (m, 1H), 1.79 (ddd, J = 12.9, 8.5, 4.7 Hz, 1H), 1.36 (d, J = 6.9 Hz, 3H), 0.94 (s, 9H). ^13^C NMR (151 MHz, DMSO-*d*_6_) δ 171.5, 170.5, 169.8, 169.3, 154.4, 154.1, 151.4, 149.2, 148.6, 148.2, 147.6, 144.6, 138.3, 137.4, 131.04, 131.00, 130.3, 130.2, 129.8, 129.6, 129.1, 128.7, 128.7, 126.8, 126.3, 125.4, 123.8, 122.0, 121.5, 120.2, 113.8, 113.0, 69.8, 69.4, 68.9, 68.7, 67.9, 66.9, 58.4, 56.3, 56.2, 55.5, 47.6, 37.6, 35.6, 35.4, 35.2, 26.3, 22.3, 15.9. HR-MS (ESI) m/z: calc. for C_56_H_63_N_6_O_9_S [M+H]^+^: 995.4377, found: 995.4353.

### (2*S*,4*R*)-4-hydroxy-1-((*S*)-2-(6-(2-methoxy-4-(6-oxo-2-phenyl-6,7-dihydro-5*H*-benzo[*b*]pyrido[2,3-*d*]azepin-4-yl)phenoxy)hexanamido)-3,3-dimethylbutanoyl)-*N*-((*S*)-1-(4-(4-methylthiazol-5-yl)phenyl)ethyl)pyrrolidine-2- carboxamide (YZ-1)

Compound **YZ-1** was synthesized in a similar manner as described for compound **YZ-6**. White solid, 55% yield. ^1^H NMR (600 MHz, DMSO-*d*_6_) δ 10.40 (s, 1H), 8.98 (s, 1H), 8.37 (d, *J* = 7.8 Hz, 1H), 8.26 – 8.19 (m, 3H), 7.96 (s, 1H), 7.83 (d, *J* = 9.3 Hz, 1H), 7.56 – 7.49 (m, 3H), 7.47 (t, *J* = 7.3 Hz, 1H), 7.43 (d, *J* = 8.2 Hz, 2H), 7.41 – 7.34 (m, 3H), 7.29 (d, *J* = 1.7 Hz, 1H), 7.26 (d, *J* = 7.9 Hz, 1H), 7.21 (dd, *J* = 8.2, 1.9 Hz, 1H), 7.15 (d, *J* = 8.3 Hz, 1H), 5.11 (d, *J* = 3.5 Hz, 1H), 4.92 (h, *J* = 6.8 Hz, 1H), 4.54 (d, *J* = 9.4 Hz, 1H), 4.43 (t, *J* = 8.0 Hz, 1H), 4.31 – 4.26 (m, 1H), 4.04 (t, *J* = 6.6 Hz, 2H), 3.91 (s, 3H), 3.62 (s, 2H), 3.31 (br s, 2H), 2.45 (d, *J* = 2.2 Hz, 3H), 2.31 (dt, *J* = 14.8, 7.7 Hz, 1H), 2.18 (ddd, *J* = 14.1, 7.8, 6.0 Hz, 1H), 2.05 – 1.98 (m, 1H), 1.79 (ddd, *J* = 22.0, 11.2, 6.0 Hz, 3H), 1.59 (tt, *J* = 13.9, 6.5 Hz, 2H), 1.49 – 1.42 (m, 2H), 1.37 (d, *J* = 7.0 Hz, 3H), 0.94 (d, *J* = 8.7 Hz, 9H). ^13^C NMR (151 MHz, DMSO-*d*_6_) δ 172.0, 171.5, 170.6, 169.6, 154.5, 154.2, 151.5, 149.3, 148.7, 148.5, 147.7, 144.7, 138.4, 137.5, 131.1, 131.1, 130.0, 129.9, 129.7, 129.2, 128.8, 128.8, 126.9, 126.4, 125.5, 123.9, 122.1, 121.6, 120.3, 114.5, 113.8, 112.9, 68.8, 68.2, 58.6, 56.4, 56.3, 55.6, 47.7, 37.7, 35.5, 35.2, 34.9, 29.0, 28.5, 26.5, 25.2, 22.4, 16.0. HR-MS (ESI) m/z: calc. for C_55_H_61_N_6_O_7_S [M+H]^+^: 949.4322, found: 949.4307.

### (2*S*,4*R*)-4-hydroxy-1-((*S*)-2-(8-(2-methoxy-4-(6-oxo-2-phenyl-6,7-dihydro-5*H*-benzo[*b*]pyrido[2,3-*d*]azepin-4-yl)phenoxy)octanamido)-3,3-dimethylbutanoyl)-*N*-((*S*)-1-(4-(4-methylthiazol-5-yl)phenyl)ethyl)pyrrolidine-2- carboxamide (YZ-2)

Compound **YZ-2** was synthesized in a similar manner as described for compound **YZ-6**. White solid, 41% yield. ^1^H NMR (600 MHz, DMSO-*d*_6_) δ 10.40 (s, 1H), 8.98 (s, 1H), 8.36 (d, *J* = 7.8 Hz, 1H), 8.26 – 8.20 (m, 3H), 7.96 (s, 1H), 7.80 (d, *J* = 9.3 Hz, 1H), 7.56 – 7.50 (m, 3H), 7.50 – 7.45 (m, 1H), 7.45 – 7.40 (m, 2H), 7.37 (dd, *J* = 7.9, 6.3 Hz, 3H), 7.29 (d, *J* = 2.1 Hz, 1H), 7.26 (dd, *J* = 8.1, 1.3 Hz, 1H), 7.20 (dd, *J* = 8.2, 2.1 Hz, 1H), 7.15 (d, *J* = 8.3 Hz, 1H), 5.10 (d, *J* = 3.5 Hz, 1H), 4.95 – 4.87 (m, 1H), 4.53 (d, *J* = 9.3 Hz, 1H), 4.42 (t, *J* = 8.1 Hz, 1H), 4.28 (p, *J* = 3.4 Hz, 1H), 4.08 – 4.00 (m, 2H), 3.91 (s, 3H), 3.64 – 3.57 (m, 2H),), 3.31 (br s, 2H), 2.45 (s, 3H), 2.31 – 2.24 (m, 1H), 2.13 (ddd, *J* = 14.3, 8.1, 6.2 Hz, 1H), 2.04 – 1.97 (m, 1H), 1.82 – 1.73 (m, 3H), 1.57 – 1.42 (m, 6H), 1.37 (d, *J* = 7.0 Hz, 3H), 1.31 – 1.27 (m, 2H), 0.94 (s, 9H). ^13^C NMR (151 MHz, DMSO-*d*_6_) δ 172.3, 171.7, 170.7, 169.7, 154.6, 154.3, 151.6, 149.4, 148.8, 148.5, 147.8, 144.7, 138.5, 137.5, 131.22, 131.20, 131.19, 130.1, 130.0, 129.7, 129.3, 128.90, 128.86, 126.9, 126.5, 125.6, 124.0, 122.2, 121.6, 120.3, 113.9, 113.0, 69.9, 68.8, 68.3, 58.6, 56.5, 55.7, 47.8, 37.8, 35.5, 35.3, 35.0, 28.8, 28.7, 28.6, 26.5, 25.55, 25.49, 22.5, 16.0. HR-MS (ESI) m/z: calc. for C_57_H_65_N_6_O_7_S [M+H]^+^: 977.4635, found: 977.4615.

### (2*S*,4*R*)-4-hydroxy-1-((*S*)-2-(10-(2-methoxy-4-(6-oxo-2-phenyl-6,7-dihydro-5*H*-benzo[*b*]pyrido[2,3-*d*]azepin-4-yl)phenoxy)decanamido)-3,3-dimethylbutanoyl)-*N*-((*S*)-1-(4-(4-methylthiazol-5-yl)phenyl)ethyl)pyrrolidine-2- carboxamide (YZ-3)

Compound **YZ-3** was synthesized in a similar manner as described for compound **YZ-6**. White solid, 50% yield. ^1^H NMR (600 MHz, DMSO-*d*_6_) δ 10.41 (s, 1H), 8.98 (s, 1H), 8.37 (d, *J* = 7.8 Hz, 1H), 8.23 (ddd, *J* = 9.6, 8.2, 1.6 Hz, 3H), 7.96 (s, 1H), 7.79 (d, *J* = 9.3 Hz, 1H), 7.56 – 7.50 (m, 3H), 7.50 – 7.41 (m, 3H), 7.40 – 7.34 (m, 3H), 7.29 (d, *J* = 2.1 Hz, 1H), 7.26 (dd, *J* = 8.1, 1.3 Hz, 1H), 7.20 (dd, *J* = 8.2, 2.1 Hz, 1H), 7.14 (d, *J* = 8.3 Hz, 1H), 5.12 – 5.08 (m, 1H), 4.91 (p, *J* = 7.1 Hz, 1H), 4.53 (d, *J* = 9.3 Hz, 1H), 4.43 (t, *J* = 8.0 Hz, 1H), 4.28 (s, 1H), 4.04 (q, *J* = 6.3 Hz, 2H), 3.91 (s, 3H), 3.65 – 3.57 (m, 2H), 3,37 (br s, 2H), 2.45 (s, 3H), 2.30 – 2.22 (m, 1H), 2.12 (ddd, *J* = 14.2, 8.1, 6.2 Hz, 1H), 2.01 (ddd, *J* = 11.3, 7.3, 2.7 Hz, 1H), 1.83 – 1.72 (m, 3H), 1.48 (ddt, *J* = 36.3, 15.0, 7.2 Hz, 5H), 1.37 (d, *J* = 7.0 Hz, 3H), 1.35 – 1.25 (m, 7H), 0.94 (s, 9H). ^13^C NMR (151 MHz, DMSO-*d*_6_) δ 172.5, 172.0, 171.1, 170.1, 155.0, 154.7, 151.9, 149.8, 149.2, 148.9, 148.2, 145.1, 138.9, 138.1, 137.9, 131.6, 130.5, 130.3, 130.2, 129.6, 129.3, 129.2, 127.3, 126.8, 126.0, 124.4, 122.6, 122.0, 120.7, 114.3, 113.4, 110.7, 69.2, 68.8, 59.0, 56.8, 56.7, 56.1, 48.2, 38.2, 36.0, 35.7, 35.4, 29.4, 29.2, 29.2, 29.2, 29.1, 26.9, 26.0, 25.9, 22.9, 16.5. HR-MS (ESI) m/z: calc. for C_59_H_69_N_6_O_7_S [M+H]^+^: 1005.4948, found: 1005.4931.

### (2S,4R)-4-hydroxy-1-((S)-2-(2-(2-(2-methoxy-4-(6-oxo-2-phenyl-6,7-dihydro-5H-benzo[b]pyrido[2,3-d]azepin-4-yl)phenoxy)ethoxy)acetamido)-3,3-dimethylbutanoyl)-N-((S)-1-(4-(4-methylthiazol-5-yl)phenyl)ethyl)pyrrolidine-2- carboxamide (YZ-4)

Compound **YZ-4** was synthesized in a similar manner as described for compound **YZ-6**. White solid, 45% yield. ^1^H NMR (600 MHz, DMSO-*d*_6_) δ 10.42 (s, 1H), 8.98 (s, 1H), 8.44 (d, J = 7.7 Hz, 1H), 8.27 – 8.20 (m, 3H), 7.97 (s, 1H), 7.56 – 7.50 (m, 3H), 7.50 – 7.46 (m, 1H), 7.46 – 7.40 (m, 3H), 7.40 – 7.33 (m, 3H), 7.31 (s, 1H), 7.26 (dd, J = 8.1, 1.2 Hz, 1H), 7.24 – 7.21 (m, 2H), 5.15 (d, J = 3.6 Hz, 1H), 4.90 (h, J = 7.6 Hz, 1H), 4.58 (d, J = 9.5 Hz, 1H), 4.46 (t, J = 8.1 Hz, 1H), 4.32 – 4.21 (m, 3H), 4.14 – 4.04 (m, 2H), 3.93 (s, 3H), 3.91 (t, J = 4.6 Hz, 2H), 3.65 – 3.57 (m, 2H), 3.31 (br s, 2H), 2.45 (s, 3H), 2.09 – 2.02 (m, 1H), 1.78 (ddd, J = 13.1, 8.8, 4.6 Hz, 1H), 1.35 (d, J = 6.9 Hz, 3H), 0.95 (s, 9H). ^13^C NMR (151 MHz, DMSO-*d*_6_) δ 171.4, 170.4, 168.9, 168.3, 154.4, 154.1, 151.4, 149.2, 148.7, 148.1, 147.6, 144.6, 138.3, 137.4, 131.04, 131.00, 130.5, 129.8, 129.6, 129.1, 128.8, 128.78, 128.71, 126.8, 126.2, 125.4, 123.8, 122.0, 121.5, 120.2, 113.8, 113.3, 69.6, 69.4, 68.7, 67.8, 58.5, 56.4, 55.6, 55.5, 47.6, 37.6, 35.6, 35.4, 26.1, 22.3, 15.9. HR-MS (ESI) m/z: calc. for C_53_H_57_N_6_O_8_S [M+H]^+^: 937.3959, found: 937.3940.

### (2*S*,4*R*)-4-hydroxy-1-((*S*)-2-(3-(2-(2-methoxy-4-(6-oxo-2-phenyl-6,7-dihydro-5*H*-benzo[*b*]pyrido[2,3-*d*]azepin-4-yl)phenoxy)ethoxy)propanamido)-3,3-dimethylbutanoyl)-*N*-((*S*)-1-(4-(4-methylthiazol-5-yl)phenyl)ethyl)pyrrolidine-2- carboxamide (YZ-5)

Compound **YZ-5** was synthesized in a similar manner as described for compound **YZ-6**. White solid, 49% yield. ^1^H NMR (600 MHz, DMSO-*d*_6_) δ 10.41 (s, 1H), 8.98 (s, 1H), 8.37 (d, *J* = 7.9 Hz, 1H), 8.26 – 8.20 (m, 3H), 7.96 (s, 1H), 7.92 (d, *J* = 9.3 Hz, 1H), 7.53 (tdd, *J* = 7.7, 5.6, 3.0 Hz, 3H), 7.50 – 7.44 (m, 1H), 7.42 (d, *J* = 8.3 Hz, 2H), 7.40 – 7.32 (m, 3H), 7.30 (d, *J* = 2.0 Hz, 1H), 7.26 (dd, *J* = 8.2, 1.2 Hz, 1H), 7.21 (dd, *J* = 8.2, 2.1 Hz, 1H), 7.16 (d, *J* = 8.3 Hz, 1H), 5.11 (d, *J* = 3.6 Hz, 1H), 4.97 – 4.87 (m, 1H), 4.55 (d, *J* = 9.3 Hz, 1H), 4.43 (t, *J* = 8.1 Hz, 1H), 4.28 (s, 1H), 4.17 (t, *J* = 4.9 Hz, 2H), 3.92 (s, 3H), 3.81 – 3.74 (m, 2H), 3.74 – 3.67 (m, 2H), 3.65 – 3.57 (m, 2H), 3.30 (br s, 2H), 2.62 – 2.55 (m, 1H), 2.45 (s, 4H), 2.43 (s, 1H), 2.01 (ddt, *J* = 15.9, 10.6, 5.2 Hz, 1H), 1.79 (ddd, *J* = 12.9, 8.5, 4.7 Hz, 1H), 1.37 (d, *J* = 7.0 Hz, 4H), 0.94 (s, 9H). ^13^C NMR (151 MHz, DMSO-*d*_6_) δ 171.4, 170.5, 169.8, 169.3, 154.4, 154.1, 151.4, 149.2, 148.6, 148.1, 147.6, 144.5, 138.3, 137.3, 131.0, 131.0, 130.2, 129.8, 129.6, 129.1, 128.7, 128.7, 128.6, 126.7, 126.2, 125.4, 123.8, 122.0, 121.4, 120.1, 113.8, 113.0, 69.7, 68.6, 68.5, 67.8, 67.0, 59.1, 58.4, 56.3, 56.2, 55.5, 47.6, 37.6, 35.2, 26.3, 22.3, 15.9. HR-MS (ESI) m/z: calc. for C_54_H_59_N_6_O_8_S [M+H]^+^: 951.4115, found: 951.4100.

### (2*S*,4*R*)-1-((*S*)-14-(*tert*-butyl)-1-(2-methoxy-4-(6-oxo-2-phenyl-6,7-dihydro-5*H*-benzo[*b*]pyrido[2,3-*d*]azepin-4-yl)phenoxy)-12-oxo-3,6,9-trioxa-13-azapentadecan-15-oyl)-4-hydroxy-*N*-((*S*)-1-(4-(4-methylthiazol-5- yl)phenyl)ethyl)pyrrolidine-2-carboxamide (YZ-7)

Compound **YZ-7** was synthesized in a similar manner as described for compound **YZ-6**. White solid, 58% yield. ^1^H NMR (600 MHz, DMSO-*d*_6_) δ 10.42 (s, 1H), 8.98 (s, 1H), 8.38 (d, *J* = 7.8 Hz, 1H), 8.27 – 8.20 (m, 3H), 7.96 (s, 1H), 7.87 (d, *J* = 9.3 Hz, 1H), 7.55 – 7.51 (m, 3H), 7.47 (t, *J* = 7.3 Hz, 1H), 7.43 (d, *J* = 8.2 Hz, 2H), 7.39 – 7.36 (m, 3H), 7.30 (d, *J* = 1.7 Hz, 1H), 7.26 (d, *J* = 7.8 Hz, 1H), 7.22 (dd, *J* = 8.2, 1.8 Hz, 1H), 7.18 (d, *J* = 8.3 Hz, 1H), 5.11 (d, *J* = 3.6 Hz, 1H), 4.94 – 4.87 (m, 1H), 4.53 (d, *J* = 9.4 Hz, 1H), 4.43 (t, *J* = 8.0 Hz, 1H), 4.28 (s, 1H), 4.22 – 4.16 (m, 2H), 3.92 (s, 3H), 3.82 – 3.79 (m, 2H), 3.64 – 3.62 (m, 3H), 3.59 (dd, *J* = 7.5, 3.8 Hz, 2H), 3.58 – 3.56 (m, 3H), 3.53 (d, *J* = 6.9 Hz, 3H), 3.50 – 3.48 (m, 1H), 3.33 (br s, 2H), 2.55 (dd, *J* = 14.5, 7.0 Hz, 1H), 2.45 (s, 3H), 2.37 (dd, *J* = 13.5, 7.2 Hz, 1H), 2.01 (t, *J* = 10.1 Hz, 1H), 1.79 (ddd, *J* = 12.9, 8.5, 4.6 Hz, 1H), 1.37 (d, *J* = 7.0 Hz, 3H), 0.93 (s, 10H). ^13^C NMR (151 MHz, DMSO-*d*_6_) δ 172.0, 171.1, 170.4, 169.9, 155.0, 154.6, 151.9, 149.7, 149.2, 148.7, 148.2, 145.1, 138.9, 137.9, 131.6, 130.8, 130.3, 130.1, 129.6, 129.3, 129.2, 127.3, 126.8, 126.0, 124.4, 122.5, 122.0, 120.7, 114.3, 113.5, 70.4, 70.3, 70.3, 70.2, 69.4, 69.2, 68.5, 67.4, 59.0, 56.8, 56.7, 56.1, 48.2, 38.2, 36.2, 35.9, 35.8, 26.8, 22.9, 16.4. HR-MS (ESI) m/z: calc. for C_58_H_67_N_6_O_10_S [M+H]^+^: 1039.4639, found: 1039.4622.

### (2*S*,4*R*)-4-hydroxy-1-((*S*)-2-(2-(2-(2-methoxy-5-(6-oxo-2-phenyl-6,7-dihydro-5*H*-benzo[*b*]pyrido[2,3-*d*]azepin-4-yl)phenoxy)ethoxy)acetamido)-3,3-dimethylbutanoyl)-*N*-((*S*)-1-(4-(4-methylthiazol-5-yl)phenyl)ethyl)pyrrolidine-2- carboxamide (YZ-8)

Compound **YZ-8** was synthesized in a similar manner as described for compound **YZ-6**. White solid, 41% yield. ^1^H NMR (600 MHz, DMSO-*d*_6_) δ 10.43 (s, 1H), 8.98 (s, 1H), 8.42 (d, *J* = 7.7 Hz, 1H), 8.26 – 8.20 (m, 3H), 7.98 (s, 1H), 7.53 (q, *J* = 7.6, 6.8 Hz, 3H), 7.45 (dd, *J* = 28.0, 7.8 Hz, 4H), 7.40 – 7.30 (m, 4H), 7.29 – 7.23 (m, 2H), 7.18 (d, *J* = 8.4 Hz, 1H), 5.13 (d, *J* = 3.6 Hz, 1H), 4.88 (q, *J* = 6.9 Hz, 1H), 4.56 (d, *J* = 9.6 Hz, 1H), 4.44 (t, *J* = 8.1 Hz, 1H), 4.31 (m, 3H), 4.08 (s, 2H), 3.91 (t, *J* = 4.5 Hz, 2H), 3.87 (s, 3H), 3.65 – 3.54 (m, 2H), 3.31 (br s, 2H), 2.45 (s, 3H), 2.09 – 2.00 (m, 1H), 1.81 – 1.73 (m, 1H), 1.35 (d, *J* = 7.0 Hz, 3H), 0.93 (s, 9H). ^13^C NMR (151 MHz, DMSO-*d*_6_) δ 171.5, 170.3, 168.9, 168.3, 154.4, 154.1, 151.4, 149.2, 149.1, 147.6, 147.6, 144.6, 138.3, 137.3, 131.0, 131.0, 131.0, 129.9, 129.8, 129.6, 129.1, 128.7, 128.6, 126.8, 126.2, 125.4, 123.8, 122.4, 121.4, 120.2, 114.8, 112.0, 69.6, 69.4, 68.7, 67.6, 58.5, 56.4, 55.6, 47.6, 37.6, 35.6, 35.4, 26.1, 22.3, 15.9, 13.4. HR-MS (ESI) m/z: calc. for C_52_H_57_N_6_O_8_S [M+H]^+^: 937.3959, found: 937.3939.

### (2*S*,4*R*)-4-hydroxy-1-((*S*)-2-(3-(2-(2-methoxy-5-(6-oxo-2-phenyl-6,7-dihydro-5*H*-benzo[*b*]pyrido[2,3-*d*]azepin-4-yl)phenoxy)ethoxy)propanamido)-3,3-dimethylbutanoyl)-*N*-((*S*)-1-(4-(4-methylthiazol-5-yl)phenyl)ethyl)pyrrolidine-2- carboxamide (YZ-9)

Compound **YZ-9** was synthesized in a similar manner as described for compound **YZ-6**. White solid, 48% yield. ^1^H NMR (600 MHz, DMSO-*d*_6_) δ 10.43 (s, 1H), 8.98 (s, 1H), 8.36 (d, *J* = 7.8 Hz, 1H), 8.26 – 8.19 (m, 3H), 7.96 (s, 1H), 7.92 (d, *J* = 9.3 Hz, 1H), 7.53 (q, *J* = 7.6 Hz, 3H), 7.46 (t, *J* = 7.2 Hz, 1H), 7.42 (d, *J* = 8.2 Hz, 2H), 7.36 (d, *J* = 8.1 Hz, 3H), 7.30 – 7.22 (m, 3H), 7.17 (d, *J* = 8.3 Hz, 1H), 5.12 – 5.08 (m, 1H), 4.90 (p, *J* = 6.8 Hz, 1H), 4.53 (d, *J* = 9.4 Hz, 1H), 4.42 (t, *J* = 8.0 Hz, 1H), 4.24 (m, 3H), 3.86 (s, 3H), 3.81 – 3.74 (m, 2H), 3.72 – 3.66 (m, 2H), 3.63 – 3.55 (m, 2H), 3.36 (br s, 2H), 2.59 (dt, *J* = 14.2, 7.0 Hz, 1H), 2.45 (s, 3H), 2.42 – 2.36 (m, 1H), 2.04 – 1.96 (m, 1H), 1.78 (ddd, *J* = 12.9, 8.5, 4.7 Hz, 1H), 1.35 (d, *J* = 7.0 Hz, 3H), 0.91 (s, 9H). ^13^C NMR (151 MHz, DMSO-*d*_6_) δ 172.1, 171.1, 170.3, 169.9, 155.0, 154.6, 151.9, 149.7, 148.2, 148.2, 145.1, 138.9, 137.9, 131.6, 131.6, 130.5, 130.4, 130.1, 129.6, 129.3, 129.2, 127.3, 126.8, 126.0, 124.4, 122.8, 122.0, 120.8, 115.2, 112.5, 69.2, 69.2, 68.3, 67.6, 59.0, 56.9, 56.8, 56.1, 48.2, 38.2, 36.2, 35.9, 35.8, 26.8, 22.9, 16.4. HR-MS (ESI) m/z: calc. for C_54_H_59_N_6_O_8_S [M+H]^+^: 951.4115, found: 951.4096.

### (2*S*,4*R*)-4-hydroxy-1-((*S*)-2-(3-(2-(2-(2-methoxy-5-(6-oxo-2-phenyl-6,7-dihydro-5*H*-benzo[*b*]pyrido[2,3-*d*]azepin-4-yl)phenoxy)ethoxy)ethoxy)propanamido)-3,3-dimethylbutanoyl)-*N*-((*S*)-1-(4-(4-methylthiazol-5- yl)phenyl)ethyl)pyrrolidine-2-carboxamide (YZ-10)

Compound **YZ-10** was synthesized in a similar manner as described for compound **YZ-6**. White solid, 55% yield. ^1^H NMR (600 MHz, DMSO-*d*_6_) δ 10.42 (s, 1H), 8.98 (s, 1H), 8.37 (d, *J* = 7.8 Hz, 1H), 8.26 – 8.20 (m, 3H), 7.97 (s, 1H), 7.87 (d, *J* = 9.3 Hz, 1H), 7.53 (q, *J* = 7.7 Hz, 3H), 7.45 (dd, *J* = 27.4, 7.7 Hz, 3H), 7.36 (dd, *J* = 7.6, 3.4 Hz, 3H), 7.30 – 7.22 (m, 3H), 7.17 (d, *J* = 8.4 Hz, 1H), 5.10 (d, *J* = 3.6 Hz, 1H), 4.93 – 4.88 (m, 1H), 4.52 (d, *J* = 9.4 Hz, 1H), 4.43 (t, *J* = 8.0 Hz, 1H), 4.25 (m, 3H), 3.87 (s, 3H), 3.80 (t, *J* = 4.6 Hz, 2H), 3.56 (ddt, *J* = 44.7, 10.2, 5.1 Hz, 8H), 3.31 (br s, 2H), 2.57 – 2.51 (m, 1H), 2.45 (s, 3H), 2.40 – 2.32 (m, 1H), 2.01 (dd, *J* = 14.2, 7.6 Hz, 1H), 1.78 (ddd, *J* = 12.9, 8.5, 4.6 Hz, 1H), 1.36 (d, *J* = 7.0 Hz, 3H), 0.92 (s, 9H). ^13^C NMR (151 MHz, DMSO-*d*_6_) δ 171.5, 170.5, 169.8, 169.3, 154.4, 154.1, 151.4, 149.1, 149.1, 147.6, 147.6, 144.6, 138.3, 137.3, 131.0, 131.0, 130.0, 129.8, 129.6, 129.1, 128.7, 128.7, 128.6, 126.8, 126.3, 125.4, 123.8, 122.2, 121.4, 120.2, 114.5, 111.9, 69.7, 69.4, 68.9, 68.6, 67.7, 66.8, 58.4, 56.3, 56.2, 55.6, 47.6, 37.6, 35.6, 35.4, 35.2, 26.3, 22.3, 15.9. HR-MS (ESI) m/z: calc. for C_56_H_63_N_6_O_9_S [M+H]^+^: 995.4377, found: 995.4361.

### 2-(2-methoxy-4-(6-oxo-2-phenyl-6,7-dihydro-5*H*-benzo[*b*]pyrido[2,3-*d*]azepin-4-yl)phenoxy)acetic acid (46)

Compound **46** was synthesized using crude compound **45** in a similar manner as described for compound **30**. White solid, 95% yield. ^1^H NMR (600 MHz, DMSO-*d*_6_) δ 13.04 (s, 1H), 10.41 (s, 1H), 8.23 (ddd, *J* = 14.1, 8.3, 1.6 Hz, 3H), 7.97 (s, 1H), 7.53 (qd, *J* = 7.0, 6.2, 1.6 Hz, 3H), 7.50 – 7.44 (m, 1H), 7.37 (td, *J* = 7.6, 1.3 Hz, 1H), 7.32 (d, *J* = 2.2 Hz, 1H), 7.26 (dd, *J* = 8.1, 1.2 Hz, 1H), 7.19 (dd, *J* = 8.2, 2.1 Hz, 1H), 7.05 (d, *J* = 8.3 Hz, 1H), 4.78 (s, 2H), 3.53 (m, 5H).

### *tert*-butyl 2-(2-methoxy-4-(6-oxo-2-phenyl-6,7-dihydro-5*H*-benzo[*b*]pyrido[2,3-*d*]azepin-4-yl)phenoxy)acetate (47)

Compound **47** was synthesized in a similar manner as described for compound **13**. White solid, 60% yield. ^1^H NMR (600 MHz, DMSO-*d*_6_) δ 10.44 (s, 1H), 8.22 (d, *J* = 8.1 Hz, 3H), 7.92 (s, 1H), 7.56 – 7.50 (m, 3H), 7.47 (t, *J* = 7.2 Hz, 1H), 7.37 (t, *J* = 7.6 Hz, 1H), 7.32 – 7.18 (m, 4H), 4.83 (s, 2H), 3.88 (s, 3H), 3.27 (br s, 2H), 1.39 (s, 9H).

### 2-(2-methoxy-4-(6-oxo-2-phenyl-6,7-dihydro-5*H*-benzo[*b*]pyrido[2,3-*d*]azepin-4-yl)phenoxy)acetic acid (48)

Compound **48** was synthesized in a similar manner as described for compound **30**. White solid, 93% yield. ^1^H NMR (600 MHz, DMSO-*d*_6_) δ 10.47 (s, 1H), 8.26 – 8.21 (m, 3H), 7.96 (s, 1H), 7.57 – 7.51 (m, 3H), 7.51 – 7.45 (m, 1H), 7.38 (td, *J* = 7.6, 1.2 Hz, 1H), 7.29 (dd, *J* = 8.2, 2.1 Hz, 1H), 7.25 (dd, *J* = 8.1, 1.2 Hz, 1H), 7.23 – 7.18 (m, 2H), 4.90 (s, 2H), 3.89 (s, 3H), 3.47 (br s, 2H).

### *N*-(2-(2-((2-(2,6-dioxopiperidin-3-yl)-1,3-dioxoisoindolin-4-yl)amino)ethoxy)ethyl)-2-(2-methoxy-4-(6-oxo-2-phenyl-6,7- dihydro-5*H*-benzo[*b*]pyrido[2,3-*d*]azepin-4-yl)phenoxy)acetamide (YZ-11)

Compound **46** (9.3 mg, 0.02 mmol), **49** (9.0 mg, 0.025 mmol), HATU (11.4 mg, 0.03 mmol), and diisopropylethylamine (7.8 mg, 0.06 mmol) were dissolved in N, N-dimethylformamide (3 ml) and stirred at 25 °C for 8 h. After the reaction was complete (monitored by TLC), water (100 mL) was added. The organic layer was extracted with ethyl acetate and washed with brine, dried over Na_2_SO_4_, filtered and then concentrated. The residue was purified further by silica gel column chromatography (DCM:methanol = 20:1) to obtain compound **YZ-11** (9.0 mg, 50%) as a yellow solid. ^1^H NMR (600 MHz, DMSO-*d*_6_) δ 11.08 (s, 1H), 10.40 (s, 1H), 8.22 (ddd, *J* = 7.4, 6.2, 1.6 Hz, 3H), 8.02 (t, *J* = 5.8 Hz, 1H), 7.94 (s, 1H), 7.57 – 7.49 (m, 4H), 7.49 – 7.44 (m, 1H), 7.37 (td, *J* = 7.5, 1.2 Hz, 1H), 7.32 (d, *J* = 2.1 Hz, 1H), 7.26 (dd, *J* = 8.1, 1.2 Hz, 1H), 7.18 (dd, *J* = 8.2, 2.1 Hz, 1H), 7.11 (dd, *J* = 16.6, 8.5 Hz, 2H), 7.00 (d, *J* = 7.0 Hz, 1H), 6.62 (t, *J* = 5.8 Hz, 1H), 5.03 (dd, *J* = 12.8, 5.4 Hz, 1H), 4.59 (s, 2H), 3.93 (s, 3H), 3.63 (t, *J* = 5.5 Hz, 2H), 3.54 (t, *J* = 5.8 Hz, 2H), 3.47 (q, *J* = 5.7 Hz, 2H), 3.38 – 3.35 (m, 4H), 2.85 (ddd, *J* = 16.9, 13.8, 5.5 Hz, 1H), 2.59 – 2.52 (m, 1H), 2.47 (dd, *J* = 13.1, 4.4 Hz, 1H), 1.97 (dtd, *J* = 13.0, 5.2, 3.1 Hz, 1H). ^13^C NMR (151 MHz, DMSO-*d*_6_) δ 172.8, 171.5, 170.1, 168.9, 167.8, 167.3, 154.5, 154.2, 149.1, 148.8, 147.6, 146.4, 138.4, 137.5, 136.2, 132.0, 131.3, 131.1, 131.1, 129.9, 129.2, 128.8, 126.9, 125.5, 123.9, 121.9, 121.6, 120.3, 117.4, 114.2, 114.1, 110.7, 109.3, 68.7, 68.6, 68.2, 55.7, 48.5, 41.7, 38.2, 35.5, 31.0, 22.1. HR-MS (ESI) m/z: calc. for C_45_H_41_N_6_O_9_ [M+H]^+^: 809.2935, found: 809.2909.

### *N*-(2-(2-(2-((2-(2,6-dioxopiperidin-3-yl)-1,3-dioxoisoindolin-4-yl)amino)ethoxy)ethoxy)ethyl)-2-(2-methoxy-4-(6-oxo-2- phenyl-6,7-dihydro-5*H*-benzo[*b*]pyrido[2,3-*d*]azepin-4-yl)phenoxy)acetamide (YZ-12)

Compound **YZ-12** was synthesized in a similar manner as described for compound **YZ-11**. Yellow solid, 65% yield. ^1^H NMR (600 MHz, DMSO-*d*_6_) δ 11.08 (s, 1H), 10.41 (s, 1H), 8.25 – 8.19 (m, 3H), 7.96 (d, *J* = 12.4 Hz, 2H), 7.57 – 7.49 (m, 4H), 7.49 – 7.43 (m, 1H), 7.37 (td, *J* = 7.6, 1.2 Hz, 1H), 7.33 (d, *J* = 2.1 Hz, 1H), 7.26 (dd, *J* = 8.2, 1.2 Hz, 1H), 7.20 (dd, *J* = 8.2, 2.1 Hz, 1H), 7.11 (t, *J* = 8.3 Hz, 2H), 7.01 (d, *J* = 7.0 Hz, 1H), 6.59 (t, *J* = 5.8 Hz, 1H), 5.04 (dd, *J* = 12.8, 5.4 Hz, 1H), 4.58 (s, 2H), 3.94 (s, 3H), 3.62 (t, *J* = 5.5 Hz, 2H), 3.60 – 3.54 (m, 4H), 3.51 – 3.48 (m, 2H), 3.44 (q, *J* = 5.6 Hz, 2H), 3.36 (s, 2H), 3.31 (s, 2H), 2.86 (ddd, *J* = 16.9, 13.8, 5.4 Hz, 1H), 2.57 (dt, *J* = 16.9, 3.3 Hz, 1H), 2.52 (d, *J* = 4.7 Hz, 1H), 2.00 (dtd, *J* = 13.0, 5.4, 2.2 Hz, 1H). ^13^C NMR (151 MHz, DMSO-*d*_6_) δ 173.3, 172.0, 170.5, 169.4, 168.2, 167.7, 155.0, 154.7, 149.6, 149.3, 148.0, 146.8, 138.9, 137.9, 136.7, 132.5, 131.9, 131.6, 131.5, 130.4, 129.7, 129.2, 127.3, 125.9, 124.4, 122.4, 122.0, 120.7, 117.8, 114.8, 114.5, 111.1, 109.7, 70.2, 70.1, 69.4, 69.3, 68.7, 56.2, 49.0, 42.1, 38.8, 35.9, 31.4, 22.6. HR-MS (ESI) m/z: calc. for C_47_H_45_N_6_O_10_ [M+H]^+^: 853.3197, found: 853.3178.

### *N*-(2-(2-(2-(2-((2-(2,6-dioxopiperidin-3-yl)-1,3-dioxoisoindolin-4-yl)amino)ethoxy)ethoxy)ethoxy)ethyl)-2-(2-methoxy-4- (6-oxo-2-phenyl-6,7-dihydro-5*H*-benzo[*b*]pyrido[2,3-*d*]azepin-4-yl)phenoxy)acetamide (YZ-13)

Compound **YZ-13** was synthesized in a similar manner as described for compound **YZ-11**. Yellow solid, 52% yield. ^1^H NMR (600 MHz, DMSO-*d*_6_) δ 11.09 (s, 1H), 10.41 (s, 1H), 8.23 (td, *J* = 8.4, 7.8, 1.6 Hz, 3H), 7.98 (d, *J* = 15.1 Hz, 2H), 7.58 – 7.50 (m, 4H), 7.49 – 7.45 (m, 1H), 7.37 (td, *J* = 7.6, 1.3 Hz, 1H), 7.34 (d, *J* = 2.1 Hz, 1H), 7.26 (dd, *J* = 8.1, 1.2 Hz, 1H), 7.21 (dd, *J* = 8.2, 2.1 Hz, 1H), 7.12 (dd, *J* = 9.7, 8.4 Hz, 2H), 7.02 (d, *J* = 7.0 Hz, 1H), 6.58 (t, *J* = 5.8 Hz, 1H), 5.05 (dd, *J* = 12.9, 5.4 Hz, 1H), 4.59 (s, 2H), 3.96 (s, 3H), 3.59 (t, *J* = 5.4 Hz, 2H), 3.56 – 3.51 (m, 8H), 3.48 (t, *J* = 5.8 Hz, 2H), 3.43 (q, *J* = 5.6 Hz, 2H), 3.34 - 3.32 (m, 4H), 2.87 (ddd, *J* = 17.1, 13.9, 5.4 Hz, 1H), 2.61 – 2.52 (m, 1H), 2.01 (dtd, *J* = 12.9, 5.3, 2.3 Hz, 1H). ^13^C NMR (151 MHz, DMSO-*d*_6_) δ 173.3, 172.0, 170.5, 169.4, 168.2, 167.8, 155.0, 154.7, 149.6, 149.4, 148.0, 146.8, 138.9, 137.9, 136.7, 132.5, 131.9, 131.6, 131.5, 130.4, 129.7, 129.2, 127.3, 125.9, 124.4, 122.5, 122.0, 120.7, 117.9, 114.8, 114.5, 111.1, 109.7, 70.3, 70.3, 70.2, 70.1, 69.3, 69.3, 68.8, 56.2, 49.0, 42.1, 38.8, 36.0, 31.4, 22.6. HR-MS (ESI) m/z: calc. for C_49_H_49_N_6_O_11_ [M+H]^+^: 897.3459, found: 897.3436.

### *N*-(2-(2-((2-(2,6-dioxopiperidin-3-yl)-1,3-dioxoisoindolin-4-yl)amino)ethoxy)ethyl)-2-(2-methoxy-5-(6-oxo-2-phenyl-6,7- dihydro-5*H*-benzo[*b*]pyrido[2,3-*d*]azepin-4-yl)phenoxy)acetamide (YZ-14)

Compound **YZ-14** was synthesized in a similar manner as described for compound **YZ-11**. Yellow solid, 73% yield. ^1^H NMR (600 MHz, DMSO-*d*_6_) δ 11.08 (s, 1H), 10.43 (s, 1H), 8.21 (t, *J* = 7.6 Hz, 3H), 7.93 (d, *J* = 13.1 Hz, 2H), 7.56 – 7.50 (m, 4H), 7.46 (t, *J* = 7.3 Hz, 1H), 7.39 – 7.34 (m, 1H), 7.33 (dd, *J* = 8.3, 1.9 Hz, 1H), 7.29 – 7.20 (m, 3H), 7.06 (d, *J* = 8.6 Hz, 1H), 7.00 (d, *J* = 7.0 Hz, 1H), 6.60 – 6.54 (m, 1H), 5.03 (dd, *J* = 12.9, 5.4 Hz, 1H), 4.68 (s, 2H), 3.89 (s, 3H), 3.52 (dt, *J* = 30.6, 5.6 Hz, 6H), 3.40 – 3.32 (m, 4H), 2.90 – 2.81 (m, 1H), 2.60 – 2.51 (m, 2H), 1.99 – 1.94 (m, 1H). ^13^C NMR (151 MHz, DMSO-*d*_6_) δ 173.2, 172.0, 170.5, 169.4, 168.3, 167.7, 155.0, 154.7, 149.9, 149.4, 147.5, 146.8, 138.9, 137.9, 136.6, 132.5, 131.6, 131.5, 130.4, 130.4, 129.7, 129.2, 127.3, 125.9, 124.4, 124.0, 122.0, 120.7, 117.8, 116.6, 112.7, 111.1, 109.7, 69.2, 69.1, 68.7, 56.3, 49.0, 42.1, 38.7, 35.9, 31.4, 22.6. HR-MS (ESI) m/z: calc. for C_45_H_41_N_6_O_9_ [M+H]^+^: 809.2935, found: 809.2919.

### *N*-(2-(2-(2-((2-(2,6-dioxopiperidin-3-yl)-1,3-dioxoisoindolin-4-yl)amino)ethoxy)ethoxy)ethyl)-2-(2-methoxy-5-(6-oxo-2- phenyl-6,7-dihydro-5*H*-benzo[*b*]pyrido[2,3-*d*]azepin-4-yl)phenoxy)acetamide (YZ-15)

Compound **YZ-15** was synthesized in a similar manner as described for compound **YZ-11**. Yellow solid, 66% yield. ^1^H NMR (600 MHz, DMSO-*d*_6_) δ 11.08 (s, 1H), 10.42 (s, 1H), 8.25 – 8.18 (m, 3H), 7.94 (s, 1H), 7.85 (t, *J* = 5.7 Hz, 1H), 7.57 – 7.49 (m, 4H), 7.46 (t, *J* = 7.3 Hz, 1H), 7.39 – 7.34 (m, 1H), 7.32 (dd, *J* = 8.3, 1.9 Hz, 1H), 7.28 – 7.23 (m, 2H), 7.21 (d, *J* = 8.4 Hz, 1H), 7.08 (d, *J* = 8.6 Hz, 1H), 7.03 – 6.98 (m, 1H), 6.56 (t, *J* = 5.7 Hz, 1H), 5.04 (dd, *J* = 12.8, 5.4 Hz, 1H), 4.66 (s, 2H), 3.89 (s, 3H), 3.55 (t, *J* = 5.5 Hz, 2H), 3.49 – 3.43 (m, 6H), 3.40 (q, *J* = 5.5 Hz, 2H), 3.32 – 3.28 (m, 4H), 2.86 (ddd, *J* = 17.0, 13.9, 5.4 Hz, 1H), 2.61 – 2.51 (m, 1H), 2.05 – 1.93 (m, 1H). ^13^C NMR (151 MHz, DMSO-*d*_6_) δ 173.3, 172.0, 170.5, 169.4, 168.2, 167.7, 155.0, 154.7, 149.9, 149.4, 147.5, 146.8, 138.9, 137.9, 136.7, 132.5, 131.6, 131.5, 130.5, 130.4, 129.7, 129.2, 127.3, 125.9, 124.4, 124.0, 122.0, 120.7, 117.8, 116.6, 112.7, 111.1, 109.7, 70.1, 70.0, 69.4, 69.3, 68.8, 56.3, 49.0, 42.1, 38.8, 35.9, 31.4, 22.6. HR-MS (ESI) m/z: calc. for C_47_H_45_N_6_O_10_ [M+H]^+^: 853.3197, found: 853.3180.

### *N*-(2-(2-(2-(2-((2-(2,6-dioxopiperidin-3-yl)-1,3-dioxoisoindolin-4-yl)amino)ethoxy)ethoxy)ethoxy)ethyl)-2-(2-methoxy-5- (6-oxo-2-phenyl-6,7-dihydro-5*H*-benzo[*b*]pyrido[2,3-*d*]azepin-4-yl)phenoxy)acetamide (YZ-16)

Compound **YZ-16** was synthesized in a similar manner as described for compound **YZ-11**. Yellow solid, 62% yield. ^1^H NMR (600 MHz, DMSO-*d*_6_) δ 11.08 (s, 1H), 10.43 (s, 1H), 8.25 – 8.20 (m, 3H), 7.94 (s, 1H), 7.85 (t, *J* = 5.7 Hz, 1H), 7.57 – 7.50 (m, 4H), 7.46 (t, *J* = 7.3 Hz, 1H), 7.39 – 7.34 (m, 1H), 7.32 (dd, *J* = 8.3, 1.9 Hz, 1H), 7.27 (d, *J* = 1.9 Hz, 1H), 7.25 (d, *J* = 8.0 Hz, 1H), 7.22 (d, *J* = 8.4 Hz, 1H), 7.10 (d, *J* = 8.6 Hz, 1H), 7.02 (d, *J* = 7.0 Hz, 1H), 6.57 (t, *J* = 5.7 Hz, 1H), 5.04 (dd, *J* = 12.9, 5.4 Hz, 1H), 4.66 (s, 2H), 3.90 (s, 3H), 3.57 (t, *J* = 5.4 Hz, 2H), 3.52 – 3.50 (m, 2H), 3.47 (dd, *J* = 5.7, 3.2 Hz, 2H), 3.45 – 3.40 (m, 8H), 3.32 – 3.27 (m, 4H), 2.86 (ddd, *J* = 17.1, 14.0, 5.4 Hz, 1H), 2.61 – 2.51 (m, 2H), 2.00 (dt, *J* = 12.8, 3.1 Hz, 1H). ^13^C NMR (151 MHz, DMSO-*d*_6_) δ 173.3, 172.0, 170.5, 169.4, 168.2, 167.8, 155.0, 154.7, 149.9, 149.4, 147.5, 146.8, 138.9, 137.9, 136.7, 132.5, 131.6, 131.5, 130.5, 130.4, 129.7, 129.2, 127.3, 125.9, 124.4, 124.0, 122.0, 120.7, 117.9, 116.6, 112.7, 111.1, 109.7, 70.2, 70.2, 70.2, 70.0, 69.3, 68.8, 56.3, 49.0, 42.1, 38.8, 35.9, 31.4, 22.6. HR-MS (ESI) m/z: calc. for C_49_H_49_N_6_O_11_ [M+H]^+^: 897.3459, found: 897.3433

### (2*S*,4*S*)-1-((*S*)-2-(3-(2-(2-bromoethoxy)ethoxy)propanamido)-3,3-dimethylbutanoyl)-4-hydroxy-*N*-(4-(4-methylthiazol-5-yl)benzyl)pyrrolidine-2-carboxamide (53)

Compound **53** was synthesized in a similar manner as described for compound **32**. White solid, 61% yield. ^1^H NMR (600 MHz, Chloroform-*d*) δ 8.69 (s, 1H), 7.49 (t, *J* = 5.8 Hz, 1H), 7.41 – 7.34 (m, 4H), 6.87 (d, *J* = 8.7 Hz, 1H), 5.52 – 5.48 (m, 1H), 4.74 (d, *J* = 9.0 Hz, 1H), 4.65 (dd, *J* = 14.9, 7.1 Hz, 1H), 4.49 (td, *J* = 9.1, 4.5 Hz, 2H), 4.30 (dd, *J* = 14.9, 5.0 Hz, 1H), 3.95 (dd, *J* = 10.9, 4.2 Hz, 1H), 3.79 (t, *J* = 6.5 Hz, 3H), 3.75 – 3.69 (m, 2H), 3.68 – 3.64 (m, 4H), 3.46 (t, *J* = 6.4 Hz, 2H), 2.53 (s, 3H), 2.49 (dt, *J* = 5.9, 4.6 Hz, 2H), 2.38 (d, *J* = 14.1 Hz, 1H), 2.18 (ddd, *J* = 14.0, 9.1, 4.9 Hz, 1H), 0.92 (s, 9H).

### (2*S*,4*S*)-4-hydroxy-1-((*S*)-2-(3-(2-(2-(2-methoxy-4-(6-oxo-2-phenyl-6,7-dihydro-5*H*-benzo[*b*]pyrido[2,3-*d*]azepin-4-yl)phenoxy)ethoxy)ethoxy)propanamido)-3,3-dimethylbutanoyl)-*N*-(4-(4-methylthiazol-5-yl)benzyl)pyrrolidine-2- carboxamide (YZ-6 NC)

Compound **YZ-6 NC** was synthesized in a similar manner as described for compound **YZ-6**. White solid, 64% yield. ^1^H NMR (600 MHz, DMSO-*d*_6_) δ 10.41 (s, 1H), 8.97 (s, 1H), 8.63 (t, *J* = 6.0 Hz, 1H), 8.26 – 8.20 (m, 3H), 7.98 – 7.91 (m, 2H), 7.56 – 7.50 (m, 3H), 7.47 (t, *J* = 7.3 Hz, 1H), 7.43 – 7.35 (m, 5H), 7.29 (d, *J* = 1.8 Hz, 1H), 7.26 (d, *J* = 8.1 Hz, 1H), 7.21 (dd, *J* = 8.2, 1.8 Hz, 1H), 7.16 (d, *J* = 8.3 Hz, 1H), 5.43 (d, *J* = 7.2 Hz, 1H), 4.48 (d, *J* = 8.9 Hz, 1H), 4.43 (dd, *J* = 15.8, 6.3 Hz, 1H), 4.36 (dd, *J* = 8.5, 6.2 Hz, 1H), 4.25 (dd, *J* = 15.8, 5.3 Hz, 1H), 4.23 – 4.19 (m, 1H), 4.19 – 4.15 (m, 2H), 3.91 (s, 4H), 3.81 – 3.76 (m, 2H), 3.61 (p, *J* = 4.2 Hz, 4H), 3.53 (dt, *J* = 11.0, 5.1 Hz, 2H), 3.44 (dd, *J* = 10.0, 5.3 Hz, 1H), 3.30 (br s, 2H), 2.54 (dd, *J* = 14.5, 6.8 Hz, 1H), 2.43 (s, 3H), 2.41 – 2.29 (m, 2H), 1.74 (dt, *J* = 12.3, 6.0 Hz, 1H), 0.95 (s, 9H). ^13^C NMR (151 MHz, DMSO-*d*_6_) δ 172.4, 171.5, 170.2, 169.8, 154.5, 154.2, 151.5, 149.3, 148.7, 148.2, 147.7, 139.2, 138.4, 137.5, 131.1, 131.1, 131.1, 130.3, 129.9, 129.7, 129.2, 128.8, 128.6, 127.4, 126.9, 125.5, 123.9, 122.1, 121.6, 120.3, 113.9, 113.1, 69.9, 69.5, 69.1, 69.0, 68.0, 66.9, 59.7, 58.5, 56.6, 55.6, 41.8, 36.9, 35.5, 35.5, 34.8, 26.3, 15.9. HR-MS (ESI) m/z: calc. for C_55_H_61_N_6_O_9_S [M+H]^+^: 981.4221, found: 981.4188.

### (2*S*,4*R*)-1-((*S*)-2-(6-(4-(4-(3,4-dimethoxyphenyl)-6-oxo-6,7-dihydro-5*H*-benzo[*b*]pyrido[2,3-*d*]azepin-2-yl)phenoxy)hexanamido)-3,3-dimethylbutanoyl)-4-hydroxy-*N*-((*S*)-1-(4-(4-methylthiazol-5-yl)phenyl)ethyl)pyrrolidine-2- carboxamide (YZ-17)

Compound **YZ-17** was synthesized in a similar manner as described for compound **YZ-6**. White solid, 44% yield. ^1^H NMR (600 MHz, DMSO-*d*_6_) δ 10.38 (s, 1H), 8.98 (s, 1H), 8.36 (d, *J* = 7.8 Hz, 1H), 8.22 – 8.14 (m, 3H), 7.89 (s, 1H), 7.82 (d, *J* = 9.3 Hz, 1H), 7.52 (td, *J* = 7.7, 1.6 Hz, 1H), 7.45 – 7.40 (m, 2H), 7.40 – 7.32 (m, 3H), 7.31 – 7.17 (m, 3H), 7.15 (d, *J* = 8.3 Hz, 1H), 7.08 – 7.02 (m, 2H), 5.10 (d, *J* = 3.5 Hz, 1H), 4.96 – 4.86 (m, 1H), 4.53 (d, *J* = 9.3 Hz, 1H), 4.42 (t, *J* = 8.0 Hz, 1H), 4.28 (q, *J* = 3.6 Hz, 1H), 4.03 (t, *J* = 6.5 Hz, 2H), 3.90 (s, 3H), 3.85 (s, 3H), 3.64 – 3.57 (m, 2H), 3.35 (br s, 2H), 2.45 (s, 3H), 2.30 (dt, *J* = 14.6, 7.6 Hz, 1H), 2.17 (ddd, *J* = 14.2, 8.0, 6.4 Hz, 1H), 2.05 – 1.97 (m, 1H), 1.82 – 1.72 (m, 3H), 1.57 (dt, *J* = 15.7, 7.0 Hz, 2H), 1.47 – 1.41 (m, 2H), 1.37 (d, *J* = 7.0 Hz, 3H), 0.94 (s, 9H). ^13^C NMR (151 MHz, DMSO-*d*_6_) δ 173.0, 172.5, 171.5, 170.5, 160.6, 155.2, 154.8, 152.4, 150.1, 149.9, 149.4, 148.6, 145.5, 138.3, 132.1, 132.0, 132.0, 131.6, 131.1, 130.7, 130.6, 129.7, 129.1, 127.3, 125.6, 124.8, 122.9, 122.4, 120.3, 115.5, 114.5, 112.7, 70.7, 69.6, 68.4, 59.4, 57.3, 57.1, 56.5, 56.5, 48.6, 38.6, 36.1, 35.7, 29.3, 27.3, 26.1, 23.3, 16.8. HR-MS (ESI) m/z: calc. for C_56_H_63_N_6_O_8_S [M+H]^+^: 979.4428, found: 979.4402.

### (2*S*,4*R*)-1-((*S*)-2-(8-(4-(4-(3,4-dimethoxyphenyl)-6-oxo-6,7-dihydro-5*H*-benzo[*b*]pyrido[2,3-*d*]azepin-2-yl)phenoxy)octanamido)-3,3-dimethylbutanoyl)-4-hydroxy-*N*-((*S*)-1-(4-(4-methylthiazol-5-yl)phenyl)ethyl)pyrrolidine-2- carboxamide (YZ-18)

Compound **YZ-18** was synthesized in a similar manner as described for compound **YZ-6**. White solid, 51% yield. ^1^H NMR (600 MHz, DMSO-*d*_6_) δ 10.38 (s, 1H), 8.98 (s, 1H), 8.37 (d, *J* = 7.8 Hz, 1H), 8.19 (tt, *J* = 10.2, 2.4 Hz, 3H), 7.88 (s, 1H), 7.79 (d, *J* = 9.3 Hz, 1H), 7.52 (ddd, *J* = 8.4, 7.4, 1.6 Hz, 1H), 7.46 – 7.40 (m, 2H), 7.40 – 7.31 (m, 3H), 7.29 – 7.23 (m, 2H), 7.21 (dd, *J* = 8.2, 2.1 Hz, 1H), 7.15 (d, *J* = 8.3 Hz, 1H), 7.07 – 7.01 (m, 2H), 5.10 (s, 1H), 4.92 (p, *J* = 7.0 Hz, 1H), 4.53 (d, *J* = 9.4 Hz, 1H), 4.43 (t, *J* = 8.0 Hz, 1H), 4.28 (s, 1H), 4.03 (t, *J* = 6.5 Hz, 2H), 3.91 (s, 3H), 3.85 (s, 3H), 3.65 – 3.57 (m, 2H), 3.37 (s, 2H), 2.45 (s, 3H), 2.27 (dt, *J* = 14.8, 7.6 Hz, 1H), 2.13 (ddd, *J* = 14.0, 8.0, 6.1 Hz, 1H), 2.01 (ddd, *J* = 11.8, 7.5, 2.8 Hz, 1H), 1.80 (ddd, *J* = 12.9, 8.5, 4.7 Hz, 1H), 1.73 (p, *J* = 6.7 Hz, 2H), 1.55 – 1.51 (m, 1H), 1.48 (t, *J* = 6.6 Hz, 1H), 1.42 (p, *J* = 7.1 Hz, 2H), 1.37 (d, *J* = 7.0 Hz, 3H), 1.34 – 1.23 (m, 4H), 0.94 (s, 9H). ^13^C NMR (151 MHz, DMSO-*d*_6_) δ 172.5, 172.1, 171.1, 170.1, 160.2, 154.8, 154.4, 151.9, 149.7, 149.5, 149.0, 148.2, 145.1, 137.9, 131.7, 131.6, 131.2, 130.7, 130.2, 130.2, 129.3, 128.7, 126.8, 125.1, 124.3, 122.5, 122.0, 119.9, 115.1, 114.1, 112.3, 69.2, 68.0, 59.0, 56.8, 56.7, 56.1, 56.1, 48.2, 38.2, 35.9, 35.7, 35.3, 29.1, 29.1, 29.0, 26.9, 25.9, 25.8, 22.9, 16.4. HR-MS (ESI) m/z: calc. for C_58_H_67_N_6_O_8_S [M+H]^+^: 1007.4741, found: 1007.4719.

### (2*S*,4*R*)-1-((*S*)-2-(10-(4-(4-(3,4-dimethoxyphenyl)-6-oxo-6,7-dihydro-5*H*-benzo[*b*]pyrido[2,3-*d*]azepin-2-yl)phenoxy)decanamido)-3,3-dimethylbutanoyl)-4-hydroxy-*N*-((*S*)-1-(4-(4-methylthiazol-5-yl)phenyl)ethyl)pyrrolidine-2- carboxamide (YZ-19)

Compound **YZ-19** was synthesized in a similar manner as described for compound **YZ-6**. White solid, 38% yield. ^1^H NMR (600 MHz, DMSO-*d*_6_) δ 10.38 (s, 1H), 8.98 (s, 1H), 8.36 (d, *J* = 7.8 Hz, 1H), 8.22 – 8.14 (m, 3H), 7.88 (s, 1H), 7.78 (d, *J* = 9.3 Hz, 1H), 7.52 (ddd, *J* = 8.6, 7.2, 1.6 Hz, 1H), 7.45 – 7.40 (m, 2H), 7.40 – 7.31 (m, 3H), 7.29 – 7.23 (m, 2H), 7.21 (dd, *J* = 8.2, 2.1 Hz, 1H), 7.15 (d, *J* = 8.3 Hz, 1H), 7.08 – 7.00 (m, 2H), 5.09 (d, *J* = 3.5 Hz, 1H), 4.91 (p, *J* = 7.2 Hz, 1H), 4.52 (d, *J* = 9.3 Hz, 1H), 4.42 (t, *J* = 8.0 Hz, 1H), 4.28 (q, *J* = 4.1, 3.6 Hz, 1H), 4.03 (q, *J* = 6.7 Hz, 2H), 3.90 (s, 3H), 3.85 (s, 3H), 3.64 – 3.56 (m, 2H), 3.36 (br s, 2H), 2.45 (s, 3H), 2.25 (dt, *J* = 15.3, 7.7 Hz, 1H), 2.14 – 2.07 (m, 1H), 2.04 – 1.97 (m, 1H), 1.82 – 1.70 (m, 3H), 1.52 (dd, *J* = 14.1, 7.3 Hz, 1H), 1.47 (d, *J* = 6.9 Hz, 1H), 1.45 – 1.41 (m, 2H), 1.37 (d, *J* = 7.0 Hz, 3H), 1.35 – 1.31 (m, 2H), 1.26 (d, *J* = 26.7 Hz, 6H), 0.93 (s, 9H). ^13^C NMR (151 MHz, DMSO-*d*_6_) δ 172.5, 172.1, 171.1, 170.1, 160.2, 154.7, 154.4, 151.9, 149.7, 149.5, 149.0, 148.2, 145.1, 137.9, 131.7, 131.6, 131.2, 130.7, 130.2, 130.2, 129.3, 128.7, 126.8, 125.1, 124.3, 122.5, 122.0, 119.9, 115.1, 114.1, 112.3, 69.2, 68.0, 59.0, 56.8, 56.7, 56.1, 56.1, 48.2, 38.2, 35.9, 35.7, 35.4, 29.4, 29.2, 29.2, 29.1, 27.0, 26.9, 26.0, 25.9, 22.9, 16.5. HR-MS (ESI) m/z: calc. for C_60_H_71_N_6_O_8_S [M+H]^+^: 1035.5054, found: 1035.5032.

### (2*S*,4*R*)-1-((*S*)-2-(3-(2-(2-(4-(4-(3,4-dimethoxyphenyl)-6-oxo-6,7-dihydro-5*H*-benzo[*b*]pyrido[2,3-*d*]azepin-2-yl)phenoxy)ethoxy)ethoxy)propanamido)-3,3-dimethylbutanoyl)-4-hydroxy-*N*-((*S*)-1-(4-(4-methylthiazol-5- yl)phenyl)ethyl)pyrrolidine-2-carboxamide (YZ-20)

Compound **YZ-20** was synthesized in a similar manner as described for compound **YZ-6**. White solid, 51% yield. ^1^H NMR (600 MHz, DMSO-*d*_6_) δ 10.39 (s, 1H), 8.98 (s, 1H), 8.37 (d, *J* = 7.8 Hz, 1H), 8.20 (dd, *J* = 7.4, 5.1 Hz, 3H), 7.91 – 7.85 (m, 2H), 7.52 (t, *J* = 8.3 Hz, 1H), 7.43 (d, *J* = 8.2 Hz, 2H), 7.36 (dd, *J* = 10.6, 7.9 Hz, 3H), 7.29 – 7.24 (m, 2H), 7.22 (dd, *J* = 8.2, 1.9 Hz, 1H), 7.16 (d, *J* = 8.3 Hz, 1H), 7.08 (d, *J* = 8.8 Hz, 2H), 5.13 – 5.09 (m, 1H), 4.92 (p, *J* = 7.0 Hz, 1H), 4.54 (d, *J* = 9.4 Hz, 1H), 4.43 (t, *J* = 8.0 Hz, 1H), 4.28 (s, 1H), 4.19 – 4.14 (m, 2H), 3.91 (s, 3H), 3.86 (s, 3H), 3.80 – 3.76 (m, 2H), 3.65 – 3.58 (m, 6H), 3.53 (dt, *J* = 15.7, 4.9 Hz, 2H), 3.37 (br s, 2H), 2.55 (dt, *J* = 14.1, 6.8 Hz, 1H), 2.45 (s, 3H), 2.40 – 2.34 (m, 1H), 2.05 – 1.98 (m, 1H), 1.79 (ddd, *J* = 12.9, 8.5, 4.7 Hz, 1H), 1.37 (d, *J* = 7.0 Hz, 3H), 0.94 (s, 10H). ^13^C NMR (151 MHz, DMSO-*d*_6_) δ 171.6, 170.6, 169.9, 169.4, 159.5, 154.2, 154.0, 151.5, 149.2, 149.0, 148.5, 147.8, 144.7, 137.4, 131.2, 131.1, 130.9, 130.2, 129.8, 129.7, 128.8, 128.2, 126.4, 124.7, 123.9, 122.0, 121.5, 119.4, 114.63, 113.61, 111.78, 69.9, 69.5, 68.9, 68.8, 67.3, 67.0, 58.6, 56.4, 56.3, 55.6, 55.6, 47.7, 37.7, 35.7, 35.4, 35.3, 26.4, 22.4, 16.0. HR-MS (ESI) m/z: calc. for C_60_H_71_N_6_O_8_S [M+H]^+^: 1025.4483, found: 1025.4448.

### 2-(4-(4-(3,4-dimethoxyphenyl)-6-oxo-6,7-dihydro-5*H*-benzo[*b*]pyrido[2,3-*d*]azepin-2-yl)phenoxy)acetic acid (55)

Compound **55** was synthesized using crude compound **54** in a similar manner as described for compound **30**. White solid, 92% yield. ^1^H NMR (600 MHz, DMSO-*d*_6_) δ 13.09 (s, 1H), 10.38 (s, 1H), 8.23 – 8.16 (m, 3H), 7.90 (s, 1H), 7.53 (td, *J* = 7.6, 1.6 Hz, 1H), 7.36 (td, *J* = 7.6, 1.3 Hz, 1H), 7.28 (d, *J* = 2.1 Hz, 1H), 7.25 (dd, *J* = 8.1, 1.2 Hz, 1H), 7.22 (dd, *J* = 8.2, 2.1 Hz, 1H), 7.16 (d, *J* = 8.3 Hz, 1H), 7.08 – 7.02 (m, 2H), 4.76 (s, 2H), 3.88 – 3.19 (m, 8H).

### 2-(4-(4-(3,4-dimethoxyphenyl)-6-oxo-6,7-dihydro-5*H*-benzo[*b*]pyrido[2,3-*d*]azepin-2-yl)phenoxy)-*N*-(2-(2-((2-(2,6-dioxopiperidin-3-yl)-1,3-dioxoisoindolin-4-yl)amino)ethoxy)ethyl)acetamide (YZ-21)

Compound **YZ-21** was synthesized in a similar manner as described for compound **YZ-11**. White solid, 50% yield. ^1^H NMR (600 MHz, DMSO-*d*_6_) δ 11.08 (s, 1H), 10.38 (s, 1H), 8.19 (td, *J* = 7.6, 1.8 Hz, 3H), 8.13 (t, *J* = 5.8 Hz, 1H), 7.89 (s, 1H), 7.56 – 7.49 (m, 2H), 7.38 – 7.33 (m, 1H), 7.29 – 7.23 (m, 2H), 7.21 (dd, *J* = 8.2, 2.1 Hz, 1H), 7.15 (d, *J* = 8.4 Hz, 1H), 7.11 (d, *J* = 8.6 Hz, 1H), 7.09 – 7.05 (m, 2H), 7.01 (d, *J* = 7.1 Hz, 1H), 6.60 (t, *J* = 5.8 Hz, 1H), 5.03 (dd, *J* = 12.8, 5.5 Hz, 1H), 4.56 (s, 2H), 3.91 (s, 3H), 3.85 (s, 3H), 3.61 (t, *J* = 5.4 Hz, 2H), 3.52 (t, *J* = 5.9 Hz, 2H), 3.45 (q, *J* = 5.6 Hz, 2H), 3.37 – 3.34 (m, 4H), 2.85 (ddd, *J* = 16.9, 13.8, 5.5 Hz, 1H), 2.55 (dt, *J* = 16.8, 3.2 Hz, 1H), 2.47 (dd, *J* = 13.1, 4.5 Hz, 1H), 2.01 – 1.94 (m, 1H). ^13^C NMR (151 MHz, DMSO-*d*_6_) δ 173.2, 172.1, 170.5, 169.4, 168.1, 167.7, 159.1, 154.6, 154.5, 149.7, 149.5, 149.0, 146.9, 137.9, 136.7, 132.5, 132.0, 131.7, 131.6, 130.7, 130.3, 128.6, 125.3, 124.4, 122.5, 122.0, 120.0, 117.9, 115.4, 114.1, 112.3, 111.1, 109.7, 69.2, 69.1, 67.4, 56.1, 56.1, 49.0, 42.1, 38.7, 35.9, 31.4, 22.6. HR-MS (ESI) m/z: calc. for C_46_H_43_N_6_O_10_ [M+H]^+^: 839.3041, found: 839.3024.

### 2-(4-(4-(3,4-dimethoxyphenyl)-6-oxo-6,7-dihydro-5*H*-benzo[*b*]pyrido[2,3-*d*]azepin-2-yl)phenoxy)-*N*-(2-(2-(2-((2-(2,6-dioxopiperidin-3-yl)-1,3-dioxoisoindolin-4-yl)amino)ethoxy)ethoxy)ethyl)acetamide (YZ-22)

Compound **YZ-22** was synthesized in a similar manner as described for compound **YZ-11**. White solid, 58% yield. ^1^H NMR (600 MHz, DMSO-*d*_6_) δ 11.08 (s, 1H), 10.38 (s, 1H), 8.22 – 8.16 (m, 3H), 8.10 (t, *J* = 5.8 Hz, 1H), 7.89 (s, 1H), 7.57 – 7.49 (m, 2H),7.36 (td, *J* = 7.5, 1.2 Hz, 1H), 7.29 – 7.23 (m, 2H), 7.21 (dd, *J* = 8.2, 2.1 Hz, 1H), 7.15 (d, *J* = 8.4 Hz, 1H), 7.11 – 7.05 (m, 3H), 7.01 (d, *J* = 7.0 Hz, 1H), 6.58 (t, *J* = 5.8 Hz, 1H), 5.04 (dd, *J* = 12.8, 5.5 Hz, 1H), 4.56 (s, 2H), 3.91 (s, 3H), 3.85 (s, 3H), 3.60 (t, *J* = 5.5 Hz, 2H), 3.57 – 3.51 (m, 5H), 3.48 (t, *J* = 5.9 Hz, 2H), 3.43 (q, *J* = 5.6 Hz, 2H), 3.33 – 3.29 (m, 4H), 2.86 (ddd, *J* = 17.0, 13.8, 5.5 Hz, 1H), 2.61 – 2.52 (m, 2H), 2.00 (dtd, *J* = 12.9, 5.3, 2.2 Hz, 1H). ^13^C NMR (151 MHz, DMSO-*d*_6_) δ 172.8, 171.6, 170.1, 168.9, 167.6, 167.3, 158.7, 154.1, 154.0, 149.2, 149.0, 148.5, 146.4, 137.4, 136.2, 132.1, 131.6, 131.2, 131.1, 130.2, 129.8, 128.2, 124.8, 123.9, 122.0, 121.5, 119.5, 117.4, 114.9, 113.6, 111.8, 110.7, 109.2, 69.7, 69.6, 68.9, 67.0, 55.6, 55.6, 48.6, 41.7, 38.3, 35.4, 31.0, 22.1. HR-MS (ESI) m/z: calc. for C_48_H_47_N_6_O_11_ [M+H]^+^: 883.3303, found: 883.3277.

### 2-(4-(4-(3,4-dimethoxyphenyl)-6-oxo-6,7-dihydro-5*H*-benzo[*b*]pyrido[2,3-*d*]azepin-2-yl)phenoxy)-*N*-(2-(2-(2-(2-((2-(2,6-dioxopiperidin-3-yl)-1,3-dioxoisoindolin-4 yl)amino)ethoxy)ethoxy)ethoxy)ethyl)acetamide (YZ-23)

Compound **YZ-23** was synthesized in a similar manner as described for compound **YZ-11**. White solid, 65% yield. ^1^H NMR (600 MHz, DMSO-*d*_6_) δ 11.09 (s, 1H), 10.39 (s, 1H), 8.23 – 8.18 (m, 3H), 8.12 (t, *J* = 5.8 Hz, 1H), 7.90 (s, 1H), 7.58 – 7.50 (m, 2H), 7.36 (td, *J* = 7.5, 1.3 Hz, 1H), 7.30 – 7.24 (m, 2H), 7.22 (dd, *J* = 8.2, 2.1 Hz, 1H), 7.16 (d, *J* = 8.4 Hz, 1H), 7.13 – 7.07 (m, 3H), 7.01 (d, *J* = 7.1 Hz, 1H), 6.58 (t, *J* = 5.8 Hz, 1H), 5.05 (dd, *J* = 12.8, 5.5 Hz, 1H), 4.57 (s, 2H), 3.91 (s, 3H), 3.86 (s, 3H), 3.59 (t, *J* = 5.5 Hz, 2H), 3.54 – 3.50 (m, 8H), 3.44 (dt, *J* = 16.3, 5.8 Hz, 4H), 3.34 – 3.28 (m, 4H), 2.88 (ddd, *J* = 17.1, 13.9, 5.5 Hz, 1H), 2.61 – 2.52 (m, 2H), 2.48 (d, *J* = 4.5 Hz, 1H), 2.01 (dtd, *J* = 13.0, 5.3, 2.4 Hz, 1H). ^13^C NMR (151 MHz, DMSO-*d*_6_) δ 173.3, 172.1, 170.5, 169.4, 168.0, 167.8, 159.1, 154.6, 154.5, 149.7, 149.5, 149.0, 146.9, 137.9, 136.7, 132.5, 132.0, 131.7, 131.6, 130.7, 130.3, 128.7, 125.3, 124.4, 122.5, 122.0, 120.0, 117.9, 115.4, 114.1, 112.3, 111.1, 109.7, 70.3, 70.2, 70.1, 69.3, 69.3, 67.5, 56.1, 56.1, 49.0, 42.1, 38.8, 35.9, 31.4, 22.6. HR-MS (ESI) m/z: calc. for C_50_H_51_N_6_O_12_ [M+H]^+^: 927.3565, found: 927.3542.

### (2*S*,4*R*)-1-((*S*)-2-acetamido-3,3-dimethylbutanoyl)-4-hydroxy-*N*-((*S*)-1-(4-(4-methylthiazol-5-yl)phenyl)ethyl)pyrrolidine-2-carboxamide (56, VHL-L)

Compound **31** (88.7 mg, 0.2 mmol), acetic anhydride (30.6 mg, 0.3 mmol), and NEt_3_ (60.7 mg, 0.6 mmol) were dissolved in DCM (5 ml) and stirred at 25 °C for 2 h. After the reaction was complete (monitored by TLC), water (20 mL) was added. The organic layer was extracted with DCM and washed with brine, dried over Na_2_SO_4_, filtered, and then concentrated. The residue was purified further by silica gel column chromatography (DCM:methanol = 25:1) to obtain compound **56** (75.9 mg, 78%) as a white solid. ^1^H NMR (600 MHz, Chloroform-*d*) δ 8.68 (s, 1H), 7.41 (d, J = 8.2 Hz, 3H), 7.37 (d, J = 8.2 Hz, 2H), 6.14 (d, J = 8.6 Hz, 1H), 5.11 – 5.05 (m, 1H), 4.74 (t, J = 7.9 Hz, 1H), 4.54 (d, J = 8.7 Hz, 2H), 4.12 (d, J = 11.5 Hz, 1H), 3.59 (dd, J = 11.4, 3.7 Hz, 1H), 2.58 (ddd, J = 13.4, 7.5, 4.7 Hz, 1H), 2.53 (s, 3H), 2.10 – 2.05 (m, 1H), 2.02 (s, 3H), 1.48 (d, J = 6.9 Hz, 3H), 1.05 (s, 9H).

#### Cell lines

HEK 293T (CRL-3216) cell line was bought from the American Type Culture Collection (ATCC). NCI-H226 cell line was a gift from Dr. Maria Zajac-Kaye’s lab at the University of Florida. The Huh7 cell line used in the study was generously provided by Dr. Liya Pi’s lab at Tulane University. All cells were grown at 37 °C with 5% CO_2_ in media as recommended by the supplier.

#### Luciferase reporter assay

HEK 293T cells were plated at 2×10^4^ cells/well in a 96 well microplate and were transfected with 50 ng of 8xGTIIC-luciferase plasmid (Addgene #34615) and 0.5 ng of pRL-CMV (Promega #E226A) by using lipofectamine 3000 (Life Technologies #L3000). After 6 h, cells were treated with various concentrations of compounds or DMSO control for 24 h. Luciferase signals were measured using the Dual Luciferase Reporter Assay System (Promega, E1960) according to the manufacturer’s instructions. Experiments were performed in triplicate and repeated at least three times.

#### Competition, Proteasome Inhibition, and Neddylation Inhibition Experiments

Between 2.0 x 10^5^ and 4.0 x 10^5^ cells were seeded into 6-well plates. The next day cells were pretreated with DMSO, 1 mM or 2 mM VHL ligand, 300 nM MG132, 200 nM or 1 μM MLN4924, or 100 μM NSC682769 for 2 h. Media was then removed, and cells were treated with DMSO, 10 μM or 20 μM **YZ-6**, 10 μM or 20 μM **YZ-6 NC**. Huh7 cells were treated for 24 h, after which cells were lysed by scraping in RIPA buffer supplemented as described previously. For an individual experiment conducted on a given day, two separate wells of cells were treated identically for every condition and harvested side-by-side

#### Western blot analysis

Treated cells were harvested and lysed in cold RIPA lysis buffer containing proteasome and phosphatase inhibitors. The protein concentrations were determined using the BCA Protein Assay kit. The prepared protein samples were loaded on and separated by 10% SDS-PAGE gel and then were transferred to PVDF membrane. Blots were blocked for 1 h at RT in blocking buffer and incubated with corresponding primary antibodies, anti-YAP (1:1000, Cell Signaling #14074), anti-CTGF (1:1000, Cell Signaling #86641), and anti-β-Actin (1:1000, Cell Signaling #4970), at 4 °C overnight. Then the blots were incubated with HRP-conjugated anti-rabbit secondary antibody in blocking buffer for 1.5 h at RT. The membranes were imaged with the ChemiDoc MP imaging system.

#### In vitro antiproliferative Assay

CCK8 assay kit was used to determine antiproliferative activity of synthesized compounds against Huh7 cell line and NCI-H226 cell line. First, cells were seeded into 96-well plates at a density of 3000 cells/well and incubated overnight. Cells were then co-incubated with compounds at various concentrations for 72 h or 96 h. Then, the cell proliferation was determined by the CCK8 kit according to the standard protocol. Experiments were performed in triplicate.

### Animal Experimentation

All animal protocols were approved by the Animal Care and Usage Committee at Tulane University and were conducted in compliance with their guidelines. *In vivo* xenograft study, immunodeficient NSG male mice (n=10, 6-week old) were purchased from Charles River Laboratories (Wilmington, MA) and were raised in an environment free of specific pathogens. Tumors were implanted by subcutaneous injection of 5×10^^6^ Huh7 cells in 100 μL of PBS containing 10% matrix gel (Corning, NY, USA), into the right flank of the mice. When the tumor volume reached about 85 mm^3^, mice were randomized into the vehicle control group (n = 5) and a treatment group (n = 5). The vehicle group (n=5) received the vehicle only (50% phosal 50 PG, 45% miglyol 810 N and 5% polysorbate 80), while the treatment group (n=5) was given compound **YZ-6** (35 mg/kg in 50% phosal 50 PG, 45% miglyol 810 N and 5% polysorbate 80) via IP injection every three days. After 24 days, mice were euthanized. Tumor size and body weight were measured every 3 days, and tumor volume was estimated using the formula volume = 1/2 × length × width^2^. Tumor growth inhibition value (TGI) was calculated as TGI (%) = [1 – Vt/Vc] × 100, where Vc and Vt were tumor volumes before and after administration.

#### Statistical analysis

Statistical analyses were conducted in Prism 9 software. Results are reported as mean ± SD. Student’s t-test was utilized to compare individual data with control values for the analysis of statistical significance. *, P < 0.05; **, P <0.01; ***, P < 0.001; ****, P < 0.0001; ns, P > 0.05.

## Acknowledgement.

This project is partially supported by the UFHCC Shands cancer fund and the Bodor Professorship Endowment fund. We would like to thank Daniel Schultz for his assistance in refining the manuscript. Special thanks are extended to Jim Rocca for providing guidance in NMR spectroscopy and to Maria Zajac-Kaye for generously providing the NCI-H226 cancer cell line. Additionally, we would like to thank Guangrong Zheng for providing us with 300 mg of compound **31**, as well as 10 mg each of compounds **42**, **43**, **44**, **49**, **50**, **51**, which greatly facilitated the synthesis process.

